# Identification of brain-like complex information architectures in embryonic tissue of *Xenopus laevis* organoids

**DOI:** 10.1101/2024.12.05.627037

**Authors:** Thomas F. Varley, Vaibhav P. Pai, Caitlin Grasso, Jeantine Lunshof, Michael Levin, Josh Bongard

**Affiliations:** Vermont Complex Systems Center, University of Vermont, Burlington, VT; Department of Computer Science, University of Vermont, Burlington, VT; Allen Discovery Institute, Tufts University, Medford, MA; Wyss Institute for Biologically Inspired Engineering at Harvard, Boston, MA; Department of Genetics, Harvard Medical School, Boston, MA, USA

**Author notes:** Co-first author (Thomas F. Varley, Vaibhav P. Pai).

**Keywords:** Developmental biology, neuroscience, functional connectivity, emergence, synergy, multivariate information theory

## Abstract

Understanding how populations of cells collectively coordinate activity to produce the complex structures and behaviors that characterize multicellular organisms, and which coordinated activities, if any, survive processes that reshape cells and tissues into organoids, are fundamental issues in modern biology. Here we show how techniques from complex systems and multivariate information theory provide a framework for inferring the structure of collective organization in non-neural tissue. Many of these techniques were developed in the context of theoretical neuroscience, where these statistics have been found to be altered during different cognitive, clinical, or behavioral states, and are generally thought to be informative about the underlying dynamics linking biology to cognition. Here we show that these same patterns of coordinated activity are also present in the aneural tissues of evolutionarily distant biological systems: preparations of embryonic *Xenopus laevis* tissue (known as “basal Xenobots”). These similarities suggest that such patterns of activity either arose independently in these two systems (epithelial constructs and brains); are epiphenomenological byproducts of other dynamics conserved across vastly different configurations of life; or somehow directly support adaptive behavior across diverse living systems. Finally, these results provide unambiguous support for the hypothesis that, despite their apparent simplicity as collections of non-neural epithelial cells, Xenobots are in fact integrated, complex systems in their own right, with sophisticated internal information structures.

## I. INTRODUCTION

One of the hallmarks of biological systems is what cybernetician Warren Weaver called “organized complexity” [1]. In contrast to simple, physical systems with just a few elements that can be well-described by formal mathematical models, biological systems display highly non-trivial structures that manifest complexity at multiple scales [2, 3]. Beyond being merely complex in the sense of having many interacting component parts, biological systems additionally have a clear capacity to engage in sophisticated self-assembly and repair of complex anatomical structures: structural complexity co-exists with dynamical and adaptive behavioral complexity. Collectives of cells and tissues homeostatically navigate the space of anatomical states during embryogenesis, regeneration, and cancer suppression [4, 5]. These states occur at scales much larger than the cells and molecular networks that implement them, requiring global coordination and virtual governor dynamics that adaptively manipulate subsystems to reach the correct species-specific target morphology and physiological states despite a range of unpredictable circumstances [4, 6]. While biologists intuitively recognize the presence of this high-level organization and control, and have noted its importance for applications in biomedicine [7, 8] and bioengineering [9], quantifying it for rigorous study has proven difficult, due to the inherent difficulties in understanding high-dimensional, multi-scale processes [10]. Consequently, major knowledge gaps exist with respect to what kind of information patterns are being implemented in neural and non-neural tissues, and how the patterns facilitate complex, adaptive self-organization and behavior.

### Identifying emergent structures in biological systems

The challenge, then, is how can we infer the structure of these high-dimensional processes given limited, and often noisy, empirical data? One of the most well-developed approaches is studying how information is distributed over the component elements of the system, and the structure of the dependencies that form the “scaffold” of on-going, spontaneous dynamics [11]. These approaches have been most well-developed in the field of computational neuroscience, where large datasets of high-temporal and high-spatial resolution time series are frequently recorded using methodologies like fMRI, EEG, and MEG [12]. From these data, a whole library of tools have been developed that characterize the structure of multivariate interactions and describe them with a variety of formal statistics. In the context of neuroscience, these techniques have revealed a rich architecture of different kinds of interactions, between neurons, neuronal networks, and whole regions of cortex [13, 14].

#### 1. Complex structures in brain activity correlates with cognitive and clinically meaningful differences

Being the most well-developed test-bed of higher-order information measures, we will begin with a brief introduction to the various measures used in this study, and what they have revealed about the information dynamics of the nervous system (a more formal review of all the measures can be found in the Materials and Methods). The first, and most fundamental inference is that of signed, functional connectivity networks, which describe the architecture of the whole system (see Sec. II D). In the brain, consistent functional connectivity networks have been found in all neuroimaging modalities [15, 16], and include a complex distribution of positively and negatively signed dependencies between brain regions, indicating correlated and anti-correlated patterns of activity. These interactions between positive and negative edges are thought to play a key role in the maintenance of non-trivial structure [17–19], and alterations to the distribution of positive and negative edges has been associated with cognitive and clinical differences [20–23]. The networks themselves are organized hierarchically, with edges arranged into a modular, meso-scale structure of communities with high internal co-fluctuation (associated with positively-signed edges) and anticorrelations between communities [14, 24], and changes in this modular structure are associated with behavioral and cognitive differences as well [25–27]. A limitation of all of these analyses is that they are built on *pairwise* models of interaction: in the functional connectivity framework, the fundamental unit is the bivariate edge, and higher-order interactions can only be inferred indirectly by considering patterns of lower-order interactions [28, 29]. To get around this limitation, there has recently been an increase in interest in truly higher-order interactions in complex systems [30], using techniques from information theory to asses polyadic synergies and redundancies directly. Analysis of neural data has found that these higher-order interactions are widespread throughout the human brain [28, 29, 31], and changes in higher-order interactions have been observed in the context of aging [32–34], neurodegeneration [35], autism [36], schizophrenia [37], and sequel to neurosurgery [38]. All of these statistics are summary statistics, which describe some feature of the *average* structure of the system. To deepen our analysis, we also can also explore time-resolved metrics. The first was the variance of the instantaneous co-fluctuation, which describes frame-resolved changes in integration-segregation balance over time [39–42], and the second was the integrated information [43], which measures how much more effectively an observer can predict the future of a whole based on a model of it’s true joint statistics versus a disintegrated model. Changes in integrated information have been found to be most closely associated with changes to consciousness following anesthesia or brain injury [44, 45]. This literature from neuroscience shows that patterns of collective behavior are not merely statistical “noise” or some kind mathematical shell game, but rather that they reflect meaningful differences in the underlying processes that generate complex cognition and behavior, sometimes with clinical or quality-of-life implications.

### A. Comparing neural data and calcium data from basal Xenobots

Despite the obvious success of functional multivariate information theory in neuroscience, however, there has been little cross-pollination with other areas of biology, leaving it unclear to what extent the patterns observed in the brain are “brain-specific”, or if these patterns of brain activity reflect larger features of information-processing dynamics across biological systems. The answer to this question would be a contribution to the emerging field of *diverse intelligence* research [46–48], which seeks to understand how information-processing capacities arise and are adaptively implemented in a wide range of material substrates and problem-spaces. To do this, we compare temporal information dynamics in two, radically diverse systems: resting state fMRI data from human brains [49], and calcium signaling data recorded from preparations of embryonic *Xenopus* tissue known colloquially as “Xenobots” [50, 51]. The term “Xenobot” is very broad and refers to a number of different of preparations: here we specifically deal with so-called “basal Xenobots”, which represent a kind of default configuration, lacking some of the additional features seen in [50, 51]. The latter (i.e. augmented or non-basal) are autonomously-motile, self-assembling constructs derived from ectoderm (epidermal progenitor) cells from the frog *Xenopus laevis*, and which have been used in the field of biorobotics as a chassis in which competencies of living materials in novel configurations can be assessed, as well as with which techniques to induce novel useful behaviors can be developed [52]. Because of their morphogenetic and behavioral properties, their lack of nervous system, their synthetic (ecologically-unique and novel) nature, and their amenability to optical interrogation, they are an interesting model system to compare to the dynamics observed in neural constructs, which have been under long periods of selection for specific functionality in a fixed configuration. One currently outstanding question in the emerging field of Xenobot bio-engineering is the extent to which the Xenobots themselves (basal or otherwise) represent truly integrated, complex systems. Given that creation of even a basal Xenobot requires fairy extreme interventions on embryonic tissues early in development, it is unclear to what extent the bots themselves maintain coherent, non-trivial structure, or if they are better understood as “blobs” of largely-independent embryonic cells that are mechanically bound together, but without any coordinated, emergent structure.

Here we systematically explore the presence of higher-order interactions in the basal Xenobot functional structure, and show that those patterns are similar to those found in the dynamics of the adult human brain. Recent prior work on calcium signaling has shown that xenopus epidermal organoid systems (with no autonomous motion behavior) do show signs of non-trivial functional organization [53], however an in-depth, comparative analysis remains an outstanding question. Multicellular patterns of calcium signaling are widespread in biology, and complex patterns (waves, bursts, and oscillations) have been found in diverse systems, including animal epithelial tissue [54, 55], plant tissue [56], and fungal mycelial networks [57]. These coordinated dynamics have been found to be crucial for many dynamical and self-organizing processing in biology; examples include skin development [58], regeneration [59], wound healing [60], and cell-type differentiation [61]. While the role of calcium transients in diverse biological processes is well-developed, and the apparent complexity of multi-cellular patterns of calcium activity have been qualitatively described in terms of waves, oscillations, and other large-scale patterns, a rigorous, formal analysis of multivariate information-processing structures in networks of signaling cells has not been done.

#### 1. Is this a fair comparison?

A natural objection to this line of work is whether the comparison of calcium signals from basal Xenobots and fMRI time series from humans is a “fair” comparison; these are different organs, different organisms, different scales, and different imaging modalities. We believe that, first, this difference is as much a feature as it is a bug, as it provides an opportunity to test the extent to which these patters, widespread in brain activity, are actually modality-specific or system-specific, and to think about what reasons could prevent applications of information metrics to specific kinds of materials. Second, these two datatypes are not as different as they appear on first blush. The key similarity is that they both describe a “whole” unit, parcellated into component “parts. “In the basal Xenobot, we have coverage of calcium activity from the camera-facing surface of the bot, while in the fMRI, we have coverage from the entire surface of the brain (the cortex). Contrast this with calcium recordings of spiking neurons (e.g. [62]), which typically only sample a small number of neurons from the whole brain. The “whole” is not recorded from in this case, making the structure of part-whole-relationships inaccessible. The data themselves also have similarities that help facilitate comparison: recordings are generally short (on the order of a few hundred frames), the signals are real-valued (unlike spiking neurons), and generally autocorrelated, which means that a common set of information-theoretic estimators and a common null model can be appropriately used in both cases. Taken together, while we acknowledge the significant differences between the two systems under study, and take pains not to over-interpret the results that follow, we also argue that it is exactly these large differences that make this kind of comparative diverse intelligences work informative.

In more typical comparative analyses, it is standard to look for and compare homologous structures (structures that descend from a common evolutionary ancestry and/or developmental origin. This is emphatically not what we are doing here. Instead of structural homologies, we are searching for what might be called “statistical” or “organizational” homologies; patterns of interactions between component parts and their collective wholes that are conserved across different species or systems. Being statistical patterns rather than physically instantiated organs or morphologies, these organizational homologies can be substrate-independent, even across phylogenically distant systems.

### B. Null models and hypothesis testing

In this paper, we are not comparing brains and basal Xenobots directly (e. g. we are not asking “do brains or basal Xenobots more total correlation” or “which system can process more information”): differences in system sizes, the properties of the recordings, and variable first order dynamics can introduce complex and hard-to-untangle biases in the estimation of information-theoretic quantities [11]. Instead, we show that both systems consistently display greater integration than you would expect if the elements were independent. Given that it is unclear to what extent basal Xenobots can develop integrated, higher-order dependencies, a null model that forces each component to be independent of all others is a natural choice. It is well known that certain lower-order features of a time series can complicate analysis of higher-order interactions [63], requiring careful consideration to avoid Type 1 errors [64].

Doing this requires constructing a bespoke null model for reach dataset that preserves all first-order features of the individual elements (autocorrelation, frequency spectrum, etc.) while disrupting any interactions between them. To do this, we use a circular-shift null model (for details see Materials and Methods) to preserve first order features while disrupting higher-order interactions between elements. We should note that this is a purely statistical null model; it makes no assumptions about the biological “hardware” running in either brains or basal Xenobots and instead only injects noise into pre-recorded data. While such a causal model would undoubtedly be useful for a more fine-grained analysis of structure-function coupling, it would require significant developments in our understanding of the structural biology of both systems, beyond what is available at present.

We hypothesized that, despite being radically different kinds of systems, that the basal Xenobots would show non-trivial information structure, indicative of complex, higher-order organization, despite the comparative “simplicity” of the system (relative to the brain). Our results largely bear this hypothesis out: we found that, for all measures tested, both the basal Xenobots and the brains showed significantly greater higher-order integration than would be expected in the null case of independent elements with preserved first-order statistics. For visualization of a sample of empirical and null covariance matrices, see Supplementary Material.

## II. MATERIALS AND METHODS

### A. Xenobot preparation

All experiments were approved by the Tufts University Institutional Animal Care and Use Committee (IACUC) under protocol number M2023-18. Xenopus Laevis embryos were fertilized in vitro according to standard protocols (Sive, 2000) and reared in 0.1X Marc’s Modified Ringer’s solution and were housed at 14oC and staged according to Nieuwkoop and Faber (Nieuwkoop and Faber, 1994). Embryos were injected with GCamp6s calcium indicator (1-2 ngs per injected blastomere) and lifeAct-mCherry cell membrane and cytoskeletal indicator (0.2-0.5 ngs per injected blastomere) mRNA in both blastomeres at 2-cell stage for uniform expression throughout the embryo. mCherry-Lifeact-7 plasmid was a gift from Michael Davidson (Addgene plasmid # 54491, http://n2t.net/addgene:54491, Accessed on 12 August 2022, RRID:Addgene 54491). The most basic version of Xenobots (no sculpting or mixing of different tissues) were generated as described previously. Briefly, stage 9 fertilized injected embryos were placed in a petri dish coated with 1% agarose made in 0.75X MMR and containing 0.75X MMR solution. Vitelline membrane was removed from the embryos using surgical forceps followed by cutting of the animal cap epidermal progenitor cells. The animal cap explants were placed on the agarose with inside surface facing up. The explants round up into spherical tissue over two hours. These explants were provided fresh dish and 0.75X MMR solution daily. Over 7 days the explant tissue differentiates and transforms into an autonomously moving synthetic epidermal entity aka Xenobot. The day 7 mature autonomously moving Xenobots were used for calcium imaging.

### B. Calcium imaging and video processing

Xenobots were imaged using Leica Stellaris Sp8 confocal microscope. Each Xenobot was imaged for 15 mins with a capture rate of one Z-stack projection every 10 secs. GCamp6s and LifeAct-mCherry were imaged in parallel with separate excitation wavelengths and separate detectors.

Individual cells in the videos were identified and segmented using Cellpose, a generalized deep learning algorithm [65]. Segmentation was performed on a representative image constructed by averaging the frames in the video converted to grayscale. As there was little movement between frames, this was sufficient to provide a good representation of the cell boundaries over the course of the video. The resulting cell segmentations were overlayed on the frames from the fluorescence channels, which captured GCaMP6s expression (there were no frame-to-frame differences in the F-actin channel, as the actin was static), and the average pixel intensity within each cell boundary was extracted to pro-duce a time series of calcium signal intensities localized to individual cells over the duration of the videos. Code for video preprocessing can be found at https://github.com/caitlingrasso/calcium-signal-extraction. Finally, the multivariate calcium traces underwent global signal regression to remove image-wide features

### C. fMRI acquisition and pre-processing

These data have been previously published on [28, 29, 66], and so we will present a minimal discussion here and refer to earlier references for the full details of the pre-processing pipeline. The data were taken from the Humman Connectome Project dataset [49]. All participants provided informed consent, and the Washington University IRB approved all protocols and procedures. A Siemens 3T Connectom Skyra equipped with a 32-channel head coil was used to collect data. Resting-state functional MRI data was acquired during four scans on two separate days. This was done with a gradient-echo echo-planar imaging (EPI) sequence (scan duration: 14:33 min; eyes open). Acquisition parameters of TR = 720 ms, TE = 33.1 ms, 52° flip angle, isotropic voxel resolution = 2 mm, with a multiband factor of 8 were used for data collection. A parcellation scheme covering the cerebral cortex developed in [67] was used to map functional data to 200 regions.

Of the 100 unrelated subjects considered in the original dataset, 50 were retained for inclusion in empirical analysis in this study. Exclusion criteria were established before the present study was conducted. They included the mean and mean absolute deviation of the relative root mean square (RMS) motion across either four resting-state MRI scans or one diffusion MRI scan, resulting in four summary motion measures. Subjects that exceeded 1.5 times the interquartile range (in the adverse direction) of the measurement distribution in two or more of these measures were excluded.

### D. Functional connectivity network inference

To build bivariate network models of the brains and the basal Xenobots, we followed a standard functional connectivity inference pipeline [13]. For every pair of elements *X*_*i*_, *X*_*j*_, we computed the Pearson correlation coefficient between their associated z-scored timeseries, which defines an *N* × *N* matrix **W** that can be interpreted as the adjacency matrix of a weighted, undirected graph **G**. We opted not to threshold the resulting covariance matrices for several reasons. The first is to maintain the link between functional connectivity matrices and Gaussian probability distributions leveraged by the higher-order information analysis, and the second was to avoid adding biases to the networks that can result from *ad hoc* thresholding procedures [68–70].

To construct a null network, we took a circular-shifting approach: each individual time-series was randomly shifted to the left or right, independently of each other, to construct a surrogate dataset that preserved all first-order features of each signal (autocorrelation, frequency spectrum, etc), but disrupted all coupling between them. Given a time series of length *T*, shifts were randomly selected integers on the range [−*T*/2 − *T*/2], sampled with uniform probability. A null functional connectivity network was then constructed, and averaged, over 1000 such instances, to create a single null “twin” for each basal Xenobot and fMRI recording. This null network contains all *apparent* higher-order information that is actually attributable to lower-order features such as autocorrelation, with the *true* higher-order information being erased.

### E. Community detection

To explore the meso-scale network structure of brains and basal Xenobots, we clustered the signed functional connectivity networks, following standard pipelines in computational neuroscience [24]. For a graph **G** = {**V, W**}, where **V** is the vertex set and **W** is the adjacency matrix of weighted edges, the standard approach to community detection involves finding the partition *σ* of the system **G** that maximizes the modularity quality function. We used the Leiden algorithm with a constant Potts model [71] to optimize the quality function:

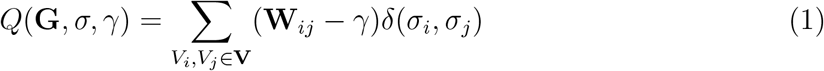

where **W**_*ij*_ is weight of the edge connecting vertices *V*_*i*_ and *V*_*j*_ in the graph **G**, and *δ*(*σ*_*i*_, *σ*_*j*_) is a delta function that returns 1 if *V*_*i*_ and *V*_*j*_ are in the same partition, and 0 otherwise. Finally, *γ* is a constant resolution parameter that determines whether the modularity function is biased towards large communities or small ones [72]. While many papers present the modularity as an intrinsic feature of a network, like the average degree, or clustering coefficinet, it is important to note that modularity is actually a feature of a partition *on* a graph, and subject to the resolution parameter. As such there is not a single “optimal” modularity or partition, but rather a space of viable partitions that must be summarized in a reasonable way. To this end we took a multi-resolution, consensus-based approach.

Ordinarily, *γ* is a free parameter that must be set in an *ad hoc* fashion, however here we adapt a modified version of the multiresolution consensus clustering algorithm [73]: we swept the value of *γ* in the range from the minimum edge weight in **W** to the maximum edge weight in **W** in 100 steps, optimizing the partition *σ* each time, and then computed a consensus partition [74] on the distribution of optimal partitions. Finally, this consensus matrix was clustered using the classic Newman-Girvan modularity with *γ* = 1 [75]. This returns a map of the optimal community structure of the network that accounts for every meaningful scale. All optimizations were done using the LeidenAlg package in Python [76].

### F. Co-fluctuation analysis

The functional connectivity analyses and their higher-order generalizations describe the expected statistical structure of a system over the duration of the entire recording. While the spatial features (e. g. cells, brain regions, etc) are preserved, the dynamic features are collapsed across time into a single summary statistic [77]. To assess whether the time-resolved dynamics between brains and basal Xenobots were similar, in addition to their long-term structures, we considered the patterns of instantaneous pairwise co-fluctuations. For each pair of cells, *X*_*i*_ and *X*_*j*_, we constructed an “edge timeseries” [39, 78]:

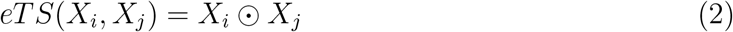

where ⊙ is the Hadamard product between the two timeseries. By doing this for all pairs, we construct a new set of of *N* ^2^ *− N/*2 edge timeseries, which tracks the instantaneous co-fluctuations of each pair of cells. If, at time *t*, both cells activity traces are both above or below their means, then the edge timeseries at time *t* will be positive, while if the cells are fluctuating in opposite directions, the co-fluctuation will be negative.

To construct a time resolved summary statistic of the global edge timeseries, we computed the root-sum-of-squares for each frame [66]:

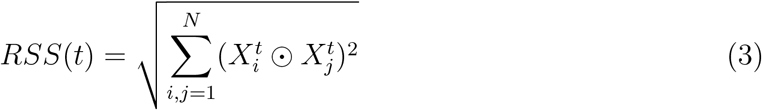

This produces a univariate timeseries that tracks the global magnitude of the co-fluctuations at each moment. We then computed the variance of the RSS timeseries, which acts as a kind of coefficient of variation, as applied to a variable continuous signal. If every *X*_*i*_⊥*X*_*j*_, then the variance of the RSS timeseries will be low (as there will be few large co-fluctuations), while if the whole system is totally synchronized, the variance will also be low, as everything is fluctuating together. When the variance is high, there is a dynamic mixture of global integration (high collective co-fluctuation) and segregation (cells are activating independently, with low co-fluctuations).

We used the same circular-shift nulls described above to compute null edge timeseries and null RSS timeseries that preserve all lower-order patterns of activity, but disrupt higher-order coordination.

### G. Higher-order information

Most statistical approaches to functional architecture in biological systems have been based on bivariate network models. While powerful, these models are limited by the fact that they can only directly assess interactions between pairs of two regions (e. g. the Pearson correlation or mutual information). Interactions between sets of three or more variables can only be inferred obliquely, as they must be constructed out of lower-order, bivariate relationships. To address this limitation, we also explore higher-order interactions between elements directly using several multivariate extensions of the classic Shannon mutual information.

#### 1. Total correlation

The first is the total correlation [79]:

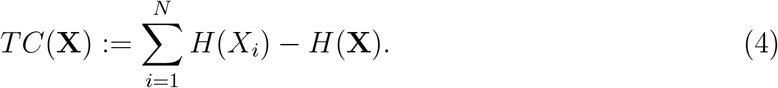

Where *H*() is the Shannon entropy function. The total correlation is a measure of how strongly a given system **X** deviates from the null case of total independence among all it’s constituent elements. It is zero if ever *X*_*i*_⊥*X*_*j*_, and it is maximal if every *X*_*i*_ is a deterministic function of some other *X*_*j*_ (for example, if all elements are perfectly synchronized). The total correlation, dual total correlation, and O-information all assume that every frame is an independent draw from from a distribution *P* (**X**), which for autocorrelated signals may not be the case, necessitating the autocorrelation-preserving null model to account for unobserved temporal dependencies.

#### 2. Dual total correlation

The second measure is the dual total correlation [80]:

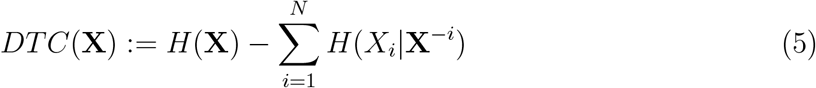

where **X**^*−i*^ refers to the joint state of every element of **X** *excluding X*_*i*_. Interestingly, the dual total correlation is low both when all the elements are independent, but also when they are all synchronized. It is high in an interstitial region where information is shared between many elements simultaneously, but not redundantly duplicated over all of them. Due to this, it has been referred to as a “genuine” higher-order interaction [35].

#### 3. O-information

The third measure of higher-order interaction we considered is the O-information [81]:

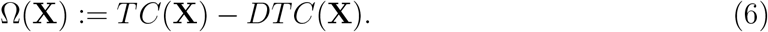

The O-information is an unusual measure in that its sign can reveal global features of the organization of **X**. If Ω(**X**) *<* 0, then the structure of **X** is dominated by synergistic interactions (i. e. information is present in the joint state of **X** and not any subset), while if Ω(**X**) *>* 0, the **X** is dominated by redundant interactions. Recently, Varley, Pope et al. [29], derived an alternative formalization of the O-information that clarifies the nature of the measure:

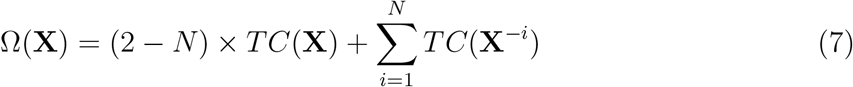

which shows that the O-information can be seen as a measure of whether the deviation from independence in **X** is better understood as occurring globally, or in some subset of **X**.

#### 4. Tononi-Sporns-Edelman Complexity

Finally, the last measure that we used was the Tononi-Sporns-Edelman complexity [82], which provides a global measure of integration/segregation balance:

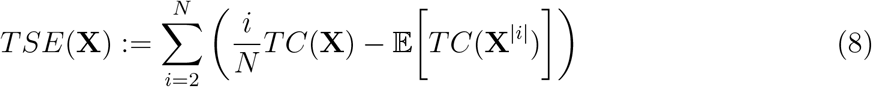

where 𝔼 [*TC*(**X**^|*i*|^)] refers to the expected total correlation of all subsets of **X** of size *i*. Like the DTC, the TSE is a non-monotonic measure: it is low if **X** is totally disintegrated and if **X** is totally synchronized, although unlike the DTC, it sweeps all possible scales of the system in sequence. Analytic work has found that the TSE complexity has a nuanced relationship with redundancy and synergy: synergy positively contributes to the complexity, while redundancy is penalized, pointing to a rich link between integration/segregation balance and redundancy/synergy balance [83].

Since computing the full TSE complexity requires brute-forcing every possible bipartition of **X**, which is impossible for large systems, we took a sub-sampling approach, as in [29, 84]. If the number of subsets of **X** of size *i* was greater than 100, we randomly sampled 100 subsets to estimate the expected value.

We used the same circular-shift nulls generated above in these analyses as well. The null functional connectivity matrices also double as null covariance matrices, and so each randomly sampled subset of brain regions was also compared to that same subset in the associated null covariance matrix.

### H. Dynamic measures

All of the results that we have discussed here have been static measures of statistical structure, assuming that joint states are drawn at random from some underlying mulitvariate probability distribution. These approaches don’t assess the dynamic nature of biological organisms: the past state of an organism informs on the future state in non-trivial ways.

### I. Bivariate transfer entropy

The bivariate transfer entropy quantifies the degree to which knowledge of source variable’s past reduces the uncertainty about a target variable’s immediate future above and beyond what could be learned by observing the target’s own past [85, 86]. Formally, for a source variable *X* and a target variable *Y* :

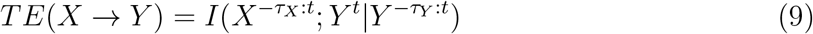

Where 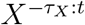 is the embedded, potentially multivariate vector of past states of 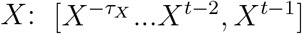. The Python-esque notation taken from [87]. The depth of histories *τ*_*X*_ and *τ*_*Y*_ are free parameters. We set *τ*_*X*_ = 5 and *τ*_*Y*_ = 1 to account for the possibility of conduction delays between brain regions and basal Xenobot cells. Note that *TE*(*X → Y*)*/*= *TE*(*Y → X*), being a temporal measure, order matters.

For every pair of regions or cells (*X, Y*), we computed *TE*(*X → Y*), and then built a distribution of 1000 circular-shifted null transfer entropies. A link *X → Y* was only considered significant if the empirical transfer entropy was greater than all of the associated autocorrelation-preserving nulls. Future work exploring the effects of the history parameters, as well as multivariate extensions (e.g. [88]).

### J. Dynamic integrated information

The space of possible dynamic measures is vast, however, in keeping with our focus on multivariate information integration, we chose to use the integrated information coefficient, which measures the degree to which the future of a system **X**^*t*+1^ is better predicted based on its own statistics, versus a null model that assumes that all *X*^*i*^ *∈* **X** are independent. Formally:

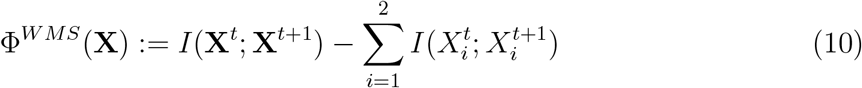

where **X**^*t*^ is the present state of **X, X**^*t*+1^ is the state of **X** after one timestep, and *WMS* refers to the “whole-minus-sum” interpretation given above.

Intuitively, the Φ function can be understood as quantifying the extent to which the “whole” **X** is greater than the sum of it’s parts. If Φ(**X**) = 0, then there is no higher-order information in the joint-state of the whole that is not learnable when considering the parts individually. **X** could be modeled just as effectively as a set of disintegrated independent processes as opposed to a coherent “whole. “A peculiar feature of the Φ function is that it can be negative, suggesting that a system is somehow less than the sum of its parts. Interpreting this stood as a long-term challenge for information theorists until Mediano et al. [89] showed that this occurred when *X*_1_ and *X*_2_ are highly correlated: i. e. that there is so much redundancy that it “swamps” the higher-order interaction. To address this, they introduced a correction:

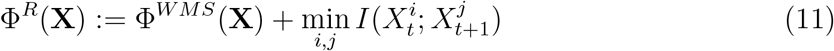

which is guaranteed to be non-negative and captures only those information-sharing modes that engage the whole system.

#### 1. Estimating the minimum information bipartition

Since the Φ^*W MS*^ is only defined for bivariate systems, computing it for the multi-cellular basal Xenobots required finding an appropriate bipartition of the system. Different bipartitions may return different values of Φ^*W MS*^, and so it is generally required that the statistic be computed with respect to the minimum information bipartition [43, 90]: that bipartition that minimizes the integrated information. This is a non-trivial optimization problem (equivalent to minimal cut problems in graph theory), and it is generally infeasible to brute-force search all possible cuts. As such, a number of heuristics have been applied over the years. Here we use the Kernighan-Lin bipartition algorithm [91], as implemented in the Net-workx package [92]. Following [93], we began with the bivariate effective connectivity graph: for each pair of regions *X*_*i*_, *X*_*j*_, we computed the mutual information between 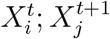 to construct weighted, undirected graph, which was then bipartitioned. The Kernighan-Lin algorithm was selected due to it’s computational efficiency and because it produces equally sized bipartitions, ensuring a balanced bisection of the bots.

Finally, we averaged the timeseries of all elements in each half to extract the global signal for each partition. These were the two time series on which we computed Φ^*W MS*^ and Φ^*R*^. As a null model, we computed a distribution of 1000 autocorrelation-preserving nulls using a circular shift randomization, and re-computed the Φ^*W MS*^ and Φ^*R*^ statistics to compare the empirical values to.

### K. Gaussian information theory

Information theory is typically used for discrete random variables, however, closed-form, analytic estimators of all the quantities used here exist for continuous, Gaussian random variables [31]. Here we will briefly introduce the basic definitions. The mutual information between two univariate variables is given by:

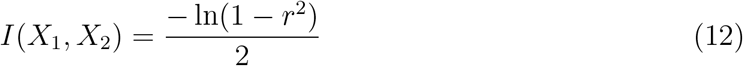

where *r* is the Pearson correlation coefficient between *X*_1_ and *X*_2_. The mutual information between two multivariate random variables is given by:

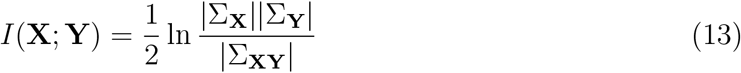

where |Σ_**X**_| is the determinant of the covariance matrix of **X**, and Σ_**XY**_ is the joint covariance matrix of **X** and **Y**. If **X** is composed of zero-mean and unit-variance variables (e.g. following a z-score transformation), the total correlation is equivalent to:

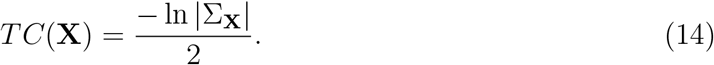

These identities are sufficient to compute all information-theoretic measures reported herein.

## III. RESULTS

In order to determine whether basal Xenobots showed patterns of information integration and emergent dynamics in a manner similar to the human brain, we computed a battery of measures on both calcium signals recorded from basal Xenobots (N=28) and fMRI BOLD signals recording from resting adult human brains (N=50). We found that, despite their comparatively simple biological structure the basal Xenobots displayed every sign of emergent complexity at significantly greater intensities than was observed in surrogate null data that preserved first-order features. This indicates that the these complex patterns are truly “emergent” and not trivially reducible to lower-order features of the data.

### A. Basal Xenobot calcium activity shows significant autocorrelation

The most basic measure of structure we studied is the first-order autocorrelation of individual cells and brain regions. For all cells and cortical parcels, in all data sets, the associated time series showed highly significant lag-one autocorrelation (quantified by the mutual information 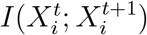, also known as the *active information storage* [11], see Fig.2). The average lag one autocorrelation for the Xenobots was 0.78 ± 0.26 nat (all p-values less than 10^*−*22^). The average autocorrelation from the fMRI scans was 1.72 ± 0.09 nat (all p-values less than 10^*−*100^). For visusalization, see Figure 2. This is significant as first-order autocorrelation can artificially bias measures of interaction strength between elements [63] and motivates the importance of the autocorrelation-preserving null model, although as we will see below, the presence of first-order autocorrelation is not sufficient to explain observed complexity in either basal Xenobots or fMRI scans.

**Figure 1.**
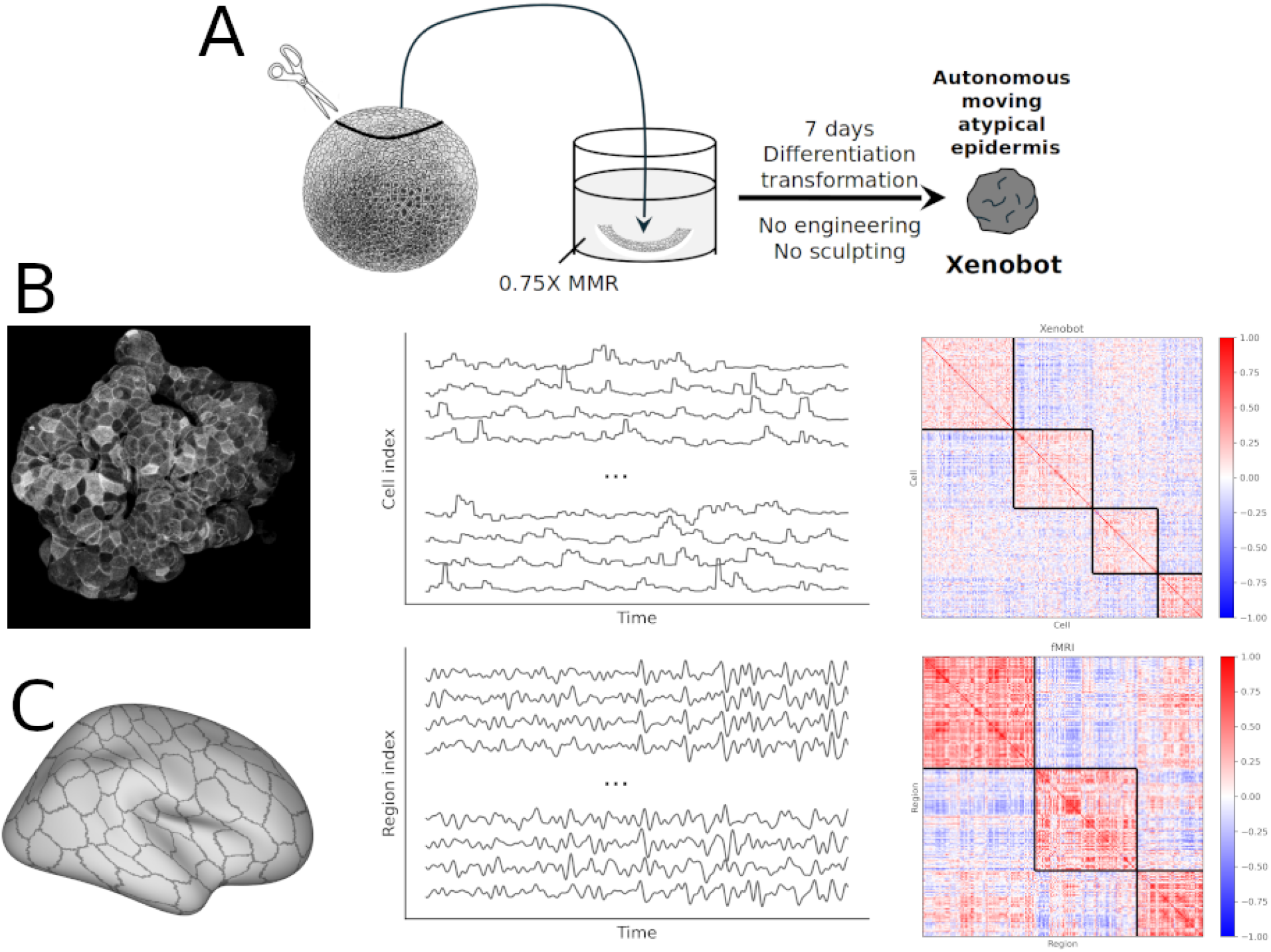
Data preparation and matrix construction. **A:** This cartoon describes the production of basal Xenobots from frog embryonic tissue. Caps are excised by hand and cultured for seven days without perturbation; the resulting autonomously moving epidermal cell structure is a “basal” (i. e. unmodified) Xenobot. **B:** The pipeline for learning functional connectivity patterns from basal Xenobots. Videos of the bots are segmented into individual cells, from which average calcium traces are extracted. The covariance matrix defines the functional connectivity, which can be organized into meso-scale communities following clustering. **C:** The same pipeline, but for human brain data recorded with an fMRI; in this case, the elements are cortical regions (corresponding to the Schaefer 200 parcellation [67]) and the signal is BOLD rather than calcium, but the overall pipeline remains the same.

**Figure 2.**
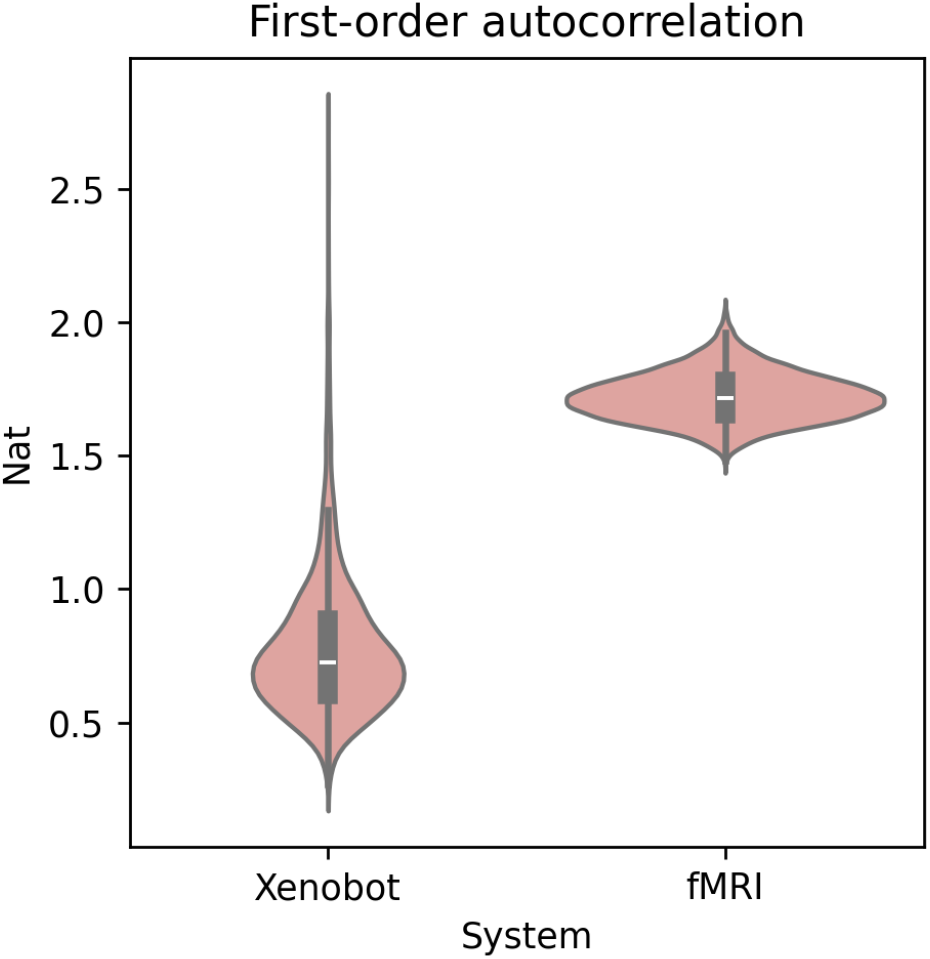
Autocorrelation. Violin plots of the distribution of autocorrelation over all brain regions (for the fMRI scans) and all cell calcium signals (for the basal Xenobot recordings. All cells and all brain regions had statistically significant autocorrelation.

### B. Basal Xenobots show non-trivial, spatially-embedded functional connectivity networks

When constructing functional connectivity networks, in every single data set, we found that positive and negative functional correlations were widely present, while in the autocorrelation-preserving nulls, spurious connections were generally weaker. These correlations form the functional connectivity network: every element in the system (cortical parcels from brains, cells for basal Xenobots) forms a vertex, and the weight of the edge between them is the Pearson correlation between their respective time series. When considering the spatial embedding of edges, we found that basal Xenobots displayed a stereotyped pattern also typically found in the brain: a negative relationship between the Euclidean distance between two elements and the strength of the dependency between them.

All twenty eight bots had statistically significant negative correlations between signed functional edge weight and distance (average *r* = *−*0.11 ± 0.05, all *p*-values less than 10^*−*9^). The same pattern was observed in the fMRI data: all brains showed a negative correla-tion between functional edge weight and Euclidean distance (average *r* = *−*0.237 ± 0.04, all *p*-values less than 10^*−*100^). For visualization see Figure 4A. These results show that the functional connectivity structure of the basal Xenobots is plausible given their spatial embedding and consistent with other spatially embed, complex biological systems. While the distance/functional connectivity relationship was certainly weaker in the basal Xenobots than in the brains, we should note that the brain regions are represented in three-dimensional space, while the basal Xenobot cells are only represented in two-dimensional space, reducing the available degrees of freedom. These results also replicate findings first observed in [53], in a closely-related organoid model system.

Multi-resolution community detection [73] finds that, in both brains and bots, these positive and negative correlations are arranged into a meso-scale structure of multiple distinct communities that are internally positively correlated, with anticorrelations between them. These communities group vertices (cortical parcels in brains, cells in basal Xenobots) into sub-systems that are more strongly connected *within* themselves than they are *between* other groups of vertices.

For visualization of example modular covarince matrices, see Figure 3, and visualizations of all communities are available in Supplementary Materials. If we consider the spatial embedding of modules, we find that, in basal Xenobots, cells that are assigned to the same modules to be closer in space than cells grouped into different modules (average Δ = *−*0.02±0.02), with 23 of 28 bots showing a statistically significant difference (*p <* 0.01, Bonferonni corrected). The relationship between spatial embedding and modules was more consistent in the fMRI scans, but followed the same pattern: average Δ = *−*0.047 ± 0.02, and all fifty recordings showed a statistically significant difference (*p <* 10^*−*15^, Bonferonni corrected). For visualization, see Figure 4B.

**Figure 3.**
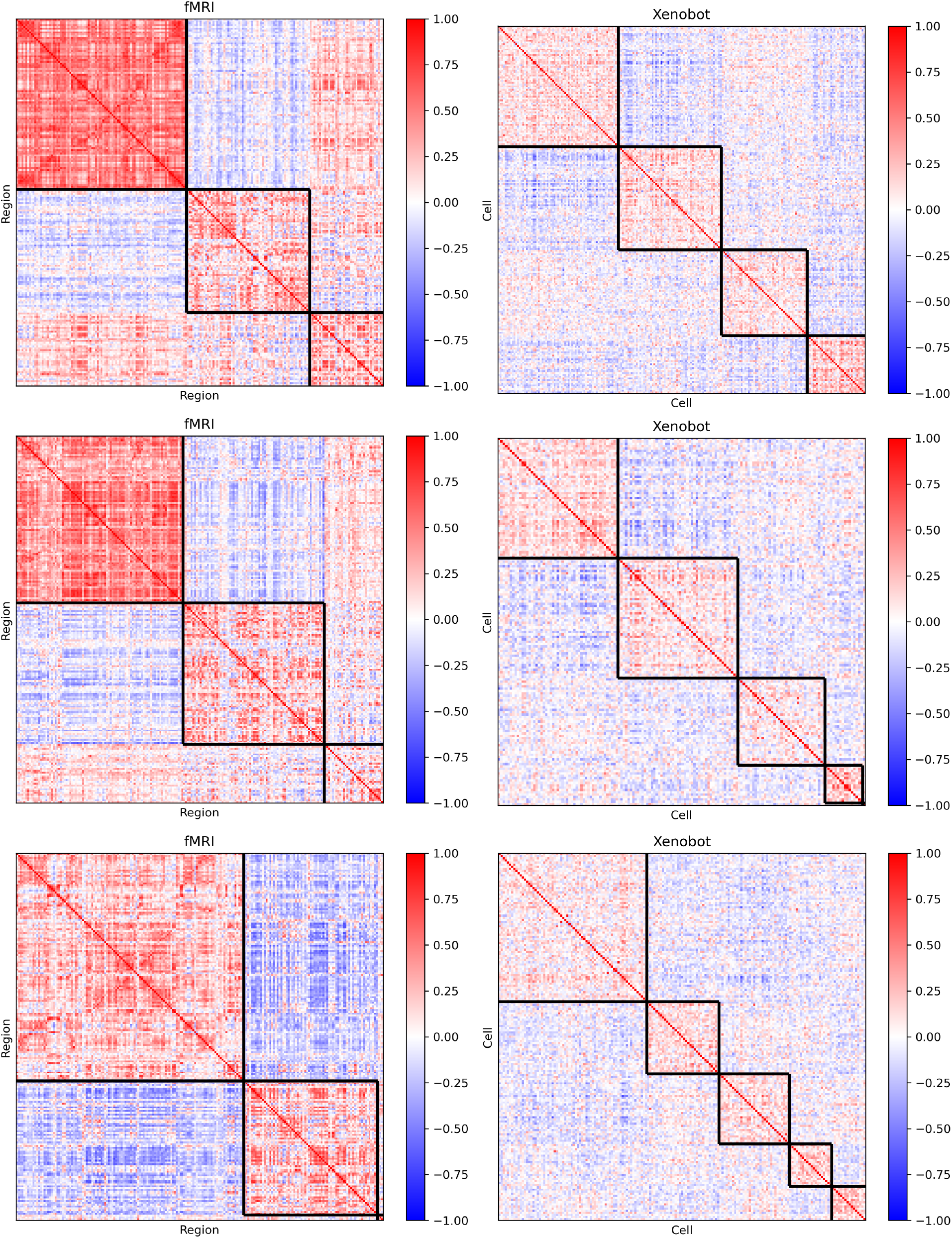
Sample communities. A set of sample functional covariance matrices, organized by the communities detected by the multi-resolution community detection algorithm [73] (see Materials and Methods II E). The on-diagonal squares (outlined in black) correspond to the communities. Three fMRI and three basal Xenobot matrices are shown. The full set of all clustered functional connectivity matrices, for basal Xenobots and fMRI scans Supplementary Material.

**Figure 4.**
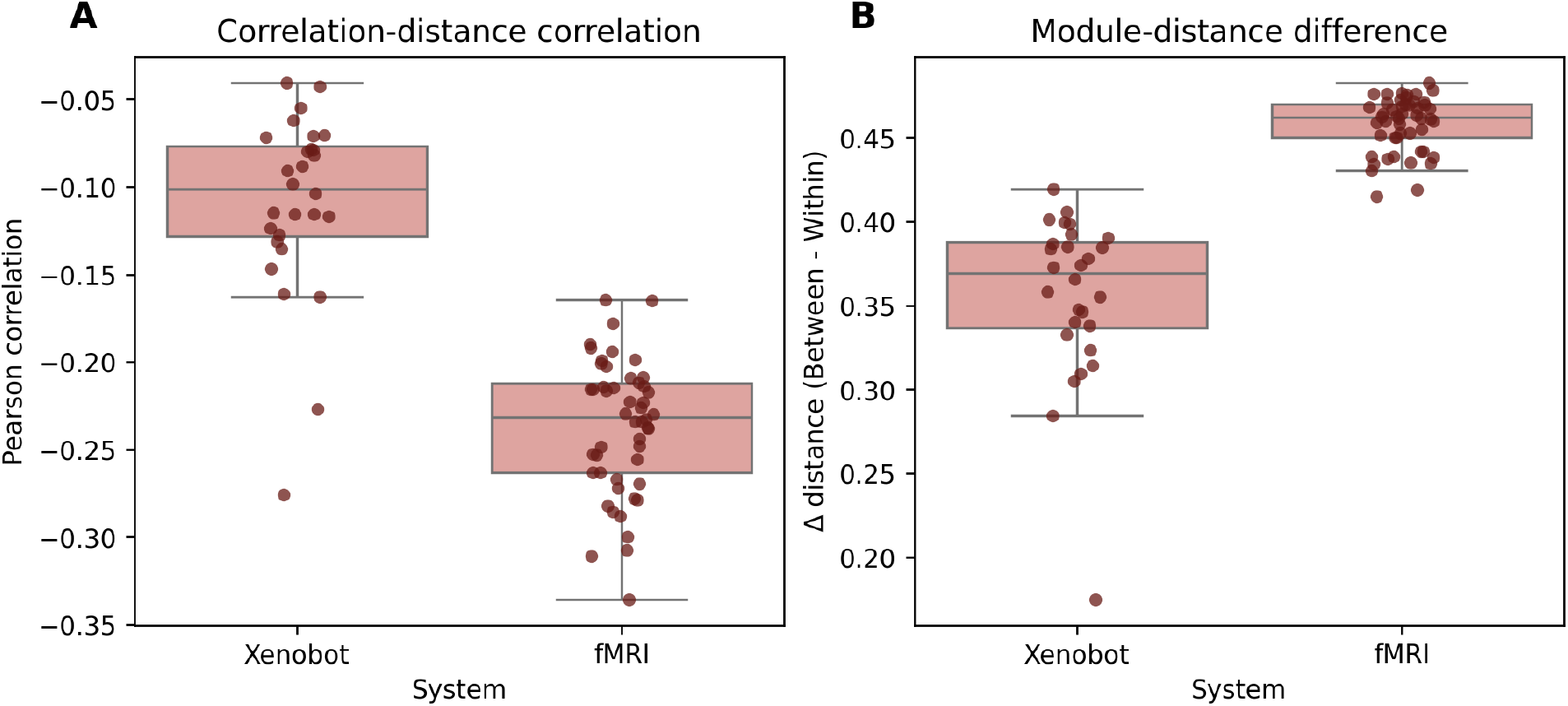
Spatial embedding of functional connections. **A** The Pearson correlation coefficient between the strength of a functional edge between two brain elements, and the Euclidean distance between them. Note that all correlations are negative, across all systems, showing a consistent drop off of functional integration with distance. **B.** The difference between the average distance between elements that both reside within a single module and the average distance between elements in different modules. All differences were positive, showing that elements grouped within a functional module tend to be closer together than elements assigned to different modules. Collectively, these results show that the functional statistics are embedded in space in plausible ways: cells that are closer together tend to be more integrated and are more likely to belong to the same meso-scale community.

### C. Time-resolved analysis shows dynamic integration and segregation balance

The functional connectivity results describe the *average* dependency between two elements. To assess whether the patterns of co-fluctuation were similar at the highest possible level of temporal resolution, we used an edge time series analysis [39, 78], which decomposes the average correlation into a series of instantaneous couplings (see Materials and Methods). To construct a summary statistic from the edge time series, we considered the variance of the root-sum-squares (RSS) for every frame across every node [39]. The variance of the RSS can be thought of as a measure of complexity balancing synchrony and independence; if all elements are independent, the variance of the RSS will be zero since there will be no periods of unusually high correlation or independence. Likewise if all time series are copies of each-other, the variance will be low, as all time series will maintain a consistent level of integration. Variance of RSS is highest when the system combines moments of instantaneous integration and segregation at different times. Both the basal Xenobots and the fMRI scans had a significantly higher empirical variance in the RSS of co-fluctuations than their respective nulls. Every single fMRI scan, and all but one of the basal Xenobots had significantly greater variance of RSS compared to the set of 1000 autocorrelation preserving nulls. The variance for the basal Xenobots was 1330.29 ± 987.5, significantly greater than the basal Xenobot nulls, which had variance of 443.65 ± 300.19 (Cohen’s D = 1.21, *p <* 10^*−*^8). In the fMRI scans, the variance was 4667.169 ± 1630.249, which was significantly greater than the null variance: 211.797 ± 14.641 (Cohen’s D = 3.865, *p <* 10^*−*^14). For visualization, see Figure 5. These results suggest that both the brains and the basal Xenobots dispslay a rich temporal dynamic, combining a mixture of transient integration and segregation. In some frames all the cells are co-fluctuating together, while in others they are behaving as if they are independent.

**Figure 5.**
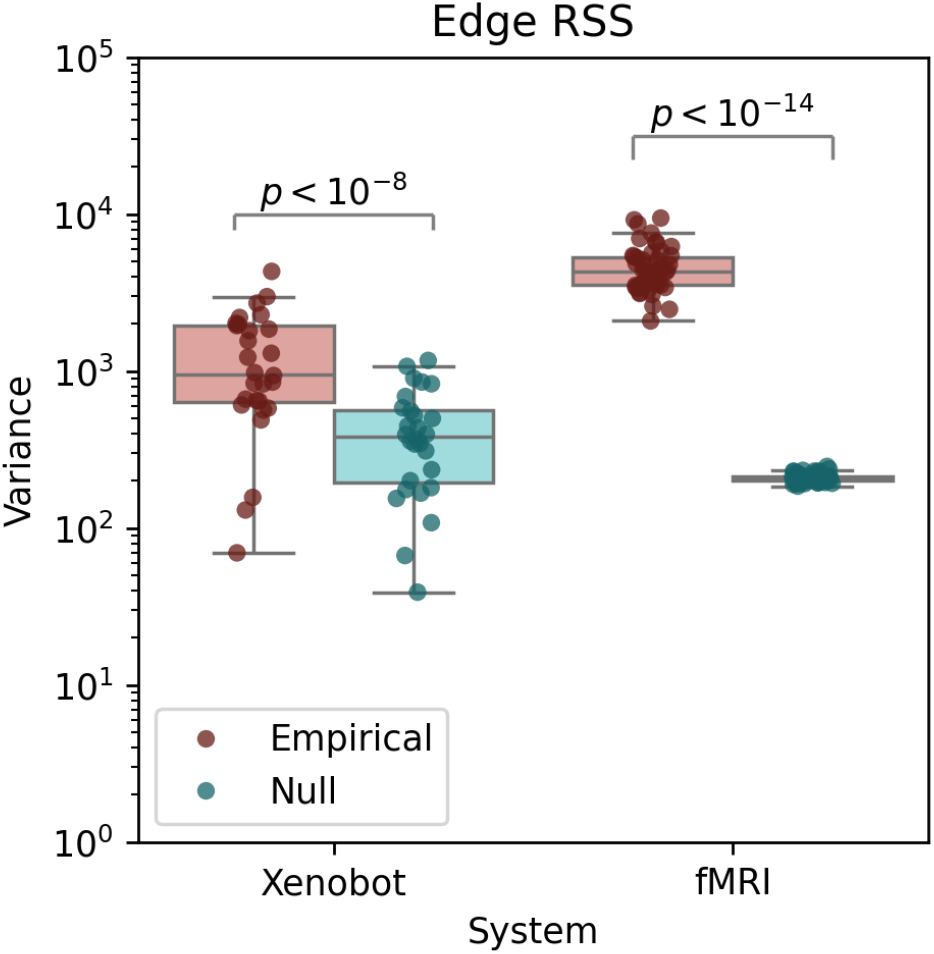
Variance in pairwise co-fluctuation magnitude. For both fMRI scans and basal Xenobot recordings, the variance of the RSS of co-fluctuation amplitude across all pairs of elements is greater than in the circular-shifted, autocorrelation-preserving nulls. These show that both fMRI scans and basal Xenobot calcium imaging data show transient, periods of both increased integration and increased segregation, indicating a kind of dynamical “richness.”

### D. Basal Xenobots show emergent, higher-order information dependencies

All of the measures discussed so far have been pairwise: describing some feature of the interaction between pairs of elements (functional connections, pairwise cofluctuations, etc). To explore genuine higher-order interactions comprising sets of three or more elements, we turned to multiviariate information, and specifically, generalizations of the mutual information: the total correlation, the dual total correlation, and the O-information [81, 87]. The total correlation and dual total correlation both quantity different notions of what it means for a set of multiple variables to “share” information [79, 80], while the O-information gives a heuristic measure of whether the overall information structure is dominated by *redundant* or *synergistic* dependencies [81].

From each data set, we sampled 1000 random triads (sets of three elements), 1000 random tetrads (sets of four elements), and pentads (sets of five elements). We then computed the total correlation, dual total correlation, and O-information for each one. For each of the samples, we then generated 1000 circular-shifted nulls to construct a null distribution of the same measures. A given set of three/four/five elements was considered to be “significant” with respect to a given measure if that measure was more extreme than all of the associated nulls (i.e. no circular-shifted null ever produced surrogate that with a value greater than the true value). When considering polyadic information describing coordinated activity between multiple elements, we found that all forms of higher-order information were significantly enriched in both brains and basal Xenobots compared to their respective nulls.

#### 1. Total correlation

The total correlation can be understood as quantifying the extent to which a given system deviates from global independence: the more global dependency exists between elements, the greater the total correlation [79]. The total correlation itself is maximized in cases of total synchrony, and minimized when very element is independent of every other one.

Of the three thousand samples, 78.73 ± 6.33% of the fMRI samples were significant, while for the basal Xenobots, only 11.21 ± 13% of samples were significant. In the fMRI data, we found significantly greater total correlation in the empirical data (0.39 ± 0.062 nat) compared to the circular-shifted nulls (0.04 ± 0.003 nat, Cohen’s d=7.93, *p <* 10^*−*14^). For visualization see Figure 6A. The same pattern appeared in basal Xenobots, with the empirical data showing significantly great total correlation (0.47 ± 0.23 nat) than the nulls (0.15 ± 0.09 nat, Cohen’s *d* = 1.79, *p <* 10^*−*9^). For visualization, see Figure 6B. Returning to the interpretation of total correlation as quantifying a global measure of inter-element dependency, we can see that the basal Xenobots, like the brains, contain sets of elements displaying non-trivial “integration”: they are not simply blobs of autonomous cells that just happen to be physically proximate, but instead appear to be coordinating calcium signaling in some way.

**Figure 6.**
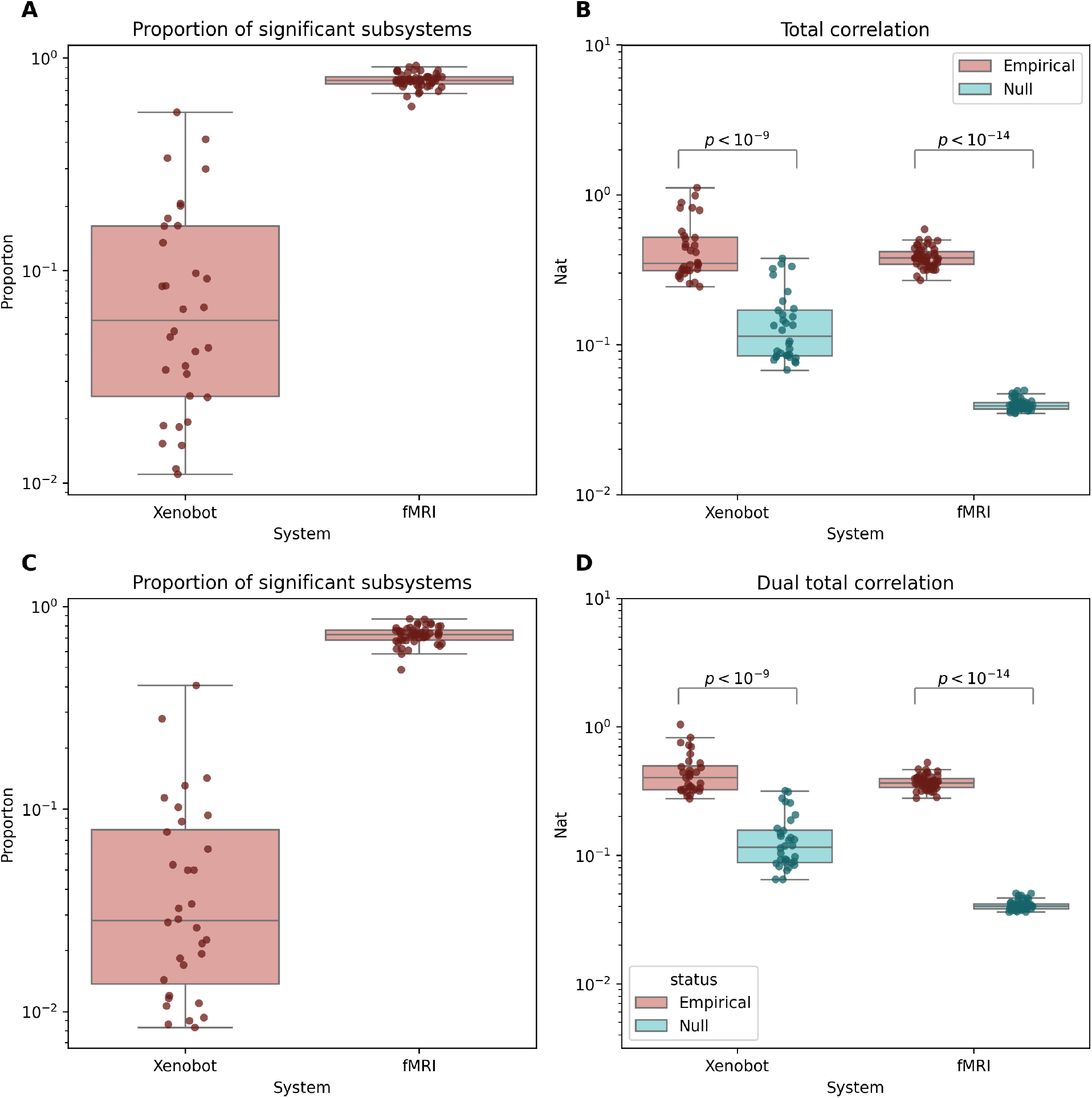
Higher-order information: total correlation and dual total correlation. **A.** The proportion of subsystems of size 3, 4, and 5 that had significant total correlation for both basal Xenobots and fMRI scans. Both systems showed significant integration for multiple samples. **B**. Both fMRI scans and basal Xenobots showed significantly greater total correlation [79] than their autocorrelation-preserving nulls. **C**. The proportion of systems of size 3, 4, and 5 that had significant dual total correlation [80] for both system types. **D**. Both fMRI scans and basal Xenobot recordings showed greater dual total correlation than their respective nulls.

To further confirm that the total correlation results (and by extension the dual total correlation and O-information results) are not attributable to the autocorrelation, we also explored the time-lagged conditional total correlation (see Supplementary Material for details).

#### 2. Dual total correlation

The dual total correlation [80] takes a different approach to higher-order information. Rather than looking at global deviation from independence, the dual total correlation quantifies the amount of information “shared” between sets of two or more elements (i.e. the amount of information that is not restricted to a single element). Of the three thousand sets of triads, tetrads, and pentads sampled, for fMRI scans, 72.64 ± 7.3% of the samples were significant, while for the basal Xenobots, only 6.2 ± 8.4% of samples were significant. For visualization see Figure 6C. In the fMRI data, we found that the empirical data showed significantly higher dual total correlation (0.372 ± 0.049 nat) than the associated nulls (0.04 ± 0.004 nat, Cohen’s D = 9.52, *p <* 10^*−*14^). In the basal Xenobot data, we similarly found that the empriical data showed significantly higher dual total correlation (0.45 ± 0.18 nat) than the nulls (0.14 ± 0.07 nat, Cohen’s d = 2.28, *p <* 10^*−*9^). For visualization, see Figure 6D. Based on the interpretation of dual total correlation as quantifying “shared information”, these results provide a different lens on multicellular coordination in basal Xenobots: not only do they form coherently integrated wholes (as evidenced by the total correlation results), but the component parts share information about each-other in non-trivial ways.

#### 3. O-information

The difference between the total correlation and the dual total correlation defines a measure called the O-information [81, 87]. Unlike the total correlation and dual total correlation, which are both strictly non-negative generalizations of mutual information, the O-information is a signed measure, which reflects the balance between different kinds of higher-order interactions. If the sign is negative, then the system in question is dominated by *synergistic* dependencies (information in the whole that is not present in the parts), while if the sign is positive, the system is dominated by *redundant* dependencies (information copied over multiple elements) [81]. Crucially, if the system is comprised of solely bivariate dependencies, then the O-information zero, and by implication, whatever information is visible must be truly polyadic - a non-zero O-information is sufficient to identify a truley higher-order relationship (although it is not necessary, as redundant and synergistic dependencies can co-exist in ways that balance out).

We first consider all sets of three, four, and five elements that have significant O-information of either sign (corresponding to those triads that have significant higher-order dependency of any kind). For the fMRI scans, we find that the average proportion of significant samples was 25% ± 14%. For the basal Xenobots, the average proportion was 2.2% ± 3.8%, a much smaller fraction. This tells us that both brains and basal Xenobots are capable of sustaining significant, higher-order/beyond pairwise dependencies. We next break the significant samples down by positive O-information (corresponding to higher-order redundancy) or negative O-information (corresponding to higher-order synergy). We find that, for the fMRI scans, of the positively-signed samples, on average 37.21 ± 9.08% were significant and of the negatively-signed samples, on average, 12.59 ± 02.3% were significant (for visualization, see Figure 7A, C). For the basal Xenobot recordings, of the positively-signed samples, on average 3.65±4.92% were significant, while for the negatively-signed samples, on average, 0.79 ± 0.87% were significant (for visualization, see Figure 7A, C). In both cases, we can see that significantly synergy-dominated samples are rarer than redundancy-dominated samples.

**Figure 7.**
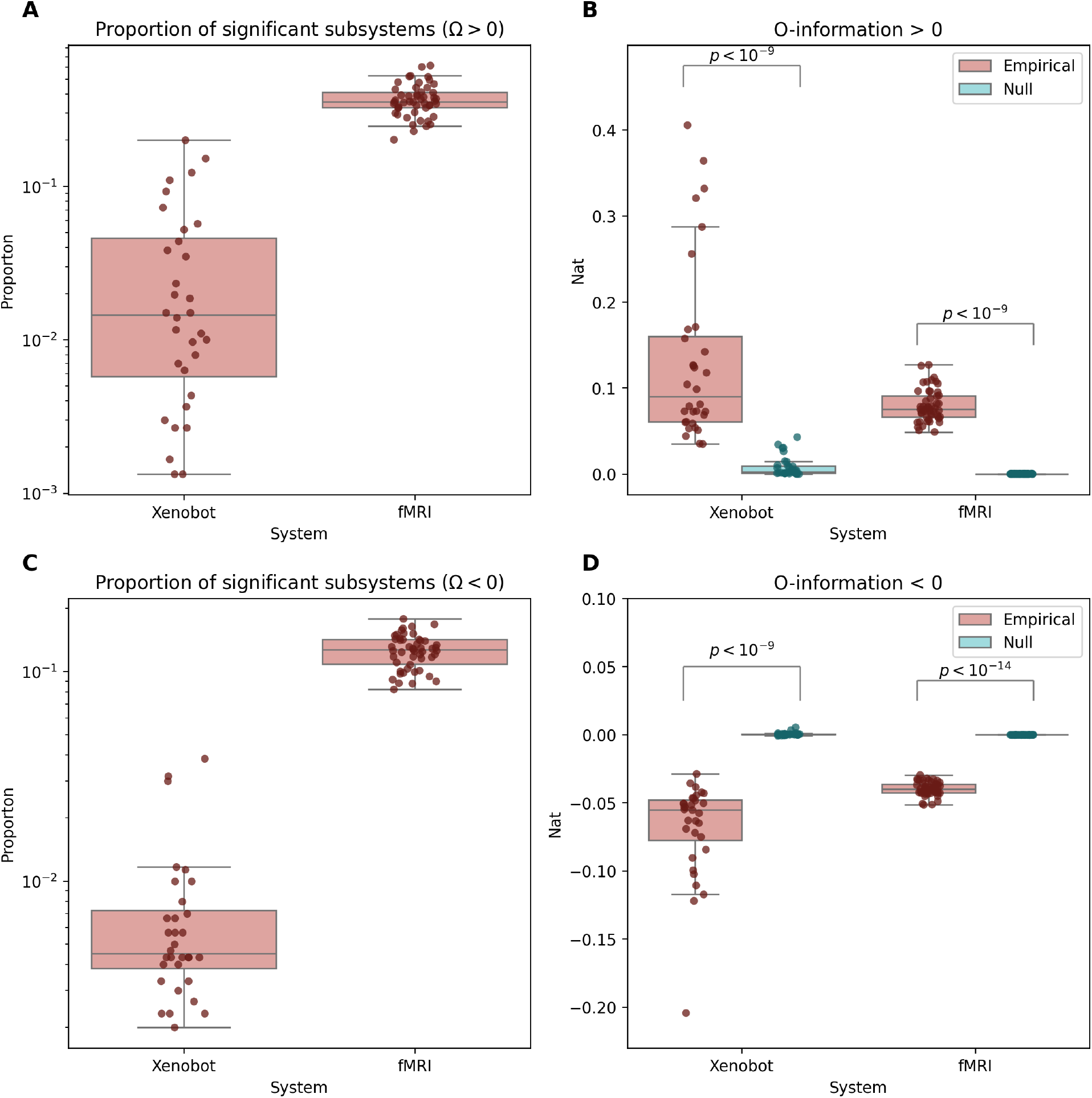
Positively- and negatively-signed O-information. **A.** The proportion of sub-systems of size 3, 4, and 5 that had significant positive O-information (i.e. were significantly redundancy-dominated) for both basal Xenobots and fMRI scans. Both systems showed significant higher-order redundancy for multiple samples. **B**. For positively-signed samples, both fMRI scans and basal Xenobots showed significantly greater O-information than their autocorrelation-preserving nulls. **C**. The proportion of systems of size 3, 4, and 5 that had significant negative O-information (i.e. were significantly synergy-dominated) for both system types. As with redundancy, both systems displayed significant synergy. **D**. For negatively-signed samples, both fMRI scans and basal Xenobot recordings showed more strongly negative O-information than their respective nulls.

In the fMRI scans, the average significant positive O-information (0.079 ± 0.019 nat) was significantly greater than the circular-shift null for those same samples (*−*0.0002 ± 7.3 *×* 10^*−*5^ nat, Cohens’d *d* = 5.98, *p <* 10^*−*9^, for visualization see Figure 7B). The average significant negative O-information was more negative (*−*0.04 ± 0.005 nat) than the nulls as well (*−*0.00019 ± 3.43 *×* 10^*−*5^ nat, Cohen’s *d* = *−*11.17, *p <* 10^*−*14^). For visualization, see Figure 7D. The same pattern was observed in the basal Xenobot recordings. The average significant positive O-information (0.13 ± 0.10 nat) was significantly greater than the associated nulls (0.008 ± 0.011 nat, Cohen’s *d* = 1.69, *p <* 10^*−*9^, for visualization see Figure 7B). The average significant negative O-information (*−*0.069 ± 0.034 nat) was significantly more negative than the circular-shift nulls (0.0003 ± 0.001 nat, Cohen’s d=-2.8, *p <* 10^*−*9^). For visualization, see Figure 7D.

These results show that, not only do basal Xenobots have statistically significant higher-order information, that they display both higher-order redundancies and higher-order synergies in different ratios, in a manner that is typical of the human brain (as recorded with fMRI). Redundancies and synergies have been found throughout the natural world; beyond brains, synergistic dependencies have also been found in sociological [94] data, climate data [95, 96], economic data [97], psychometric data [98], physical phase changes [99], and even the structure of musical scores [100]. Consequently, higher-order synergy may reflect a very general kind of “integrated sophistication” common to many complex systems.

#### 4. Tononi-Sporns-Edelman complexity

The final higher-order functional connectivity measure that we used was the Tononi-Sporns-Edelman (TSE) complexity [82], which provides a measure of the balance between integration and segregation (somewhat analogous to the O-information’s balance of redundancy and synergy). A high TSE complexity suggests that integration and segregation are co-existing: with a global deviation from independence co-existing alongside local segregation. Here, for each data set, we computed the empirical TSE complexity and a set of 1000 circular-shifted nulls, with 100 samples computed for each “level” of the TSE profile (for details, see Sec. II G 4).

For both the basal Xenobots and the fMRI scans, every single brain and bot showed significantly greater TSE complexity than their respective nulls (i.e. no individual null ever had a TSE greater than the empirical value; *p* = 0). In the fMRI data, we found that the TSE complexity was much greater in the empirical data (19183.19±493.89 nat) compared to the associated nulls (9944.57 ± 608.57, Cohen’s D = 16.76, *p* < 10^1^.7764 × 10^−15^). The same pattern is apparent in the basal Xenobots, with the empirical bots having a very high TSE complexity (25917.18 ± 36934.56 nat) compared to the associated nulls (6006.3 ± 12434.09, Cohen’s D = 0.722, *p* < 10^−5^). These results collectively show that the functional structure of the basal Xenobots show sophisticated, multi-cell coordination that is highly similar to what is seen in human brain activity. For visualization, see Figure 8.

**Figure 8.**
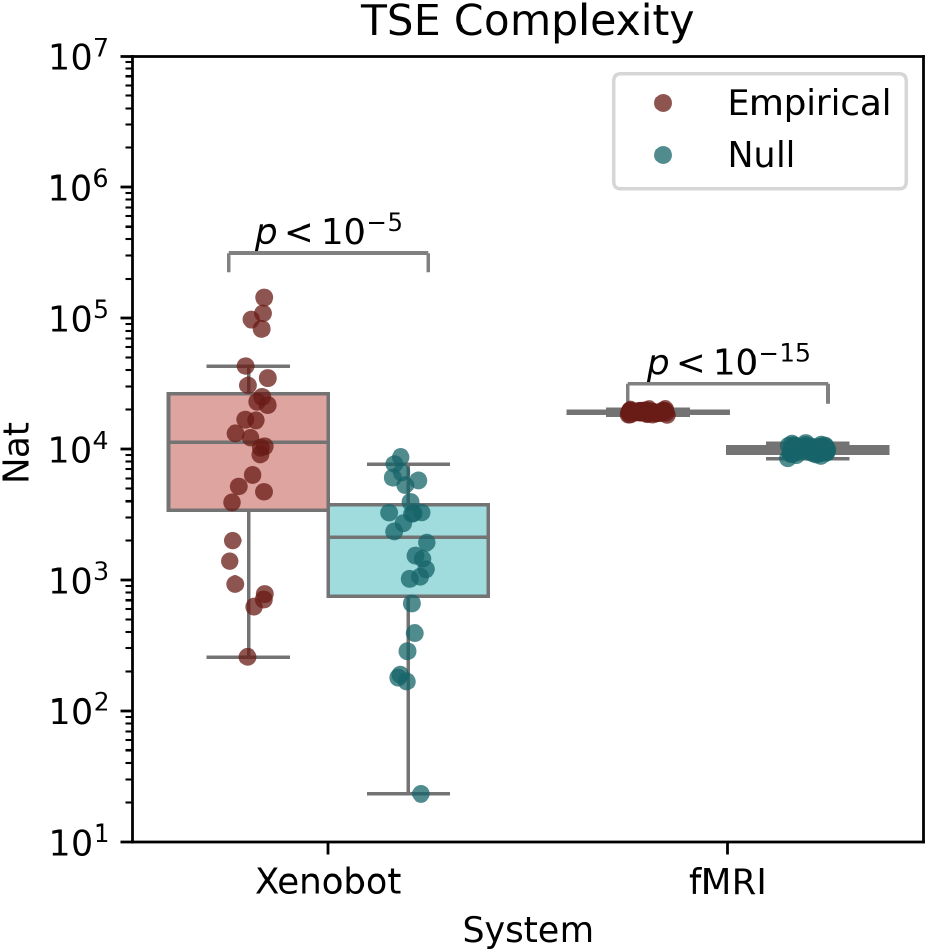
Tononi-Sporns-Edelman complexity. Both fMRI scans and basal Xenobot recordings showed higher TSE complexity [82] than their respective autocorrelation-preserving null models. These results show that the higher-order structure in brains and bots spans the whole system, involving multiple orders of interaction. The TSE complexity is understood as being maximized by a balanced combination of integration and segregation: it is low both when the whole system is synchronized, and also when all elements are independent. It is maximized when higher-order integration co-exists with lower-order independence.

### E. Dynamic measures

So far, all of the measures that we have computed are *functional* measures: they model the dynamics of the BOLD signal or calcium signal as independent frames. We also wanted to consider the *effective* connectivity, which quantifies the extent to which the past state of elements discloses information about their collective future: this is a dynamic measure that accounts for the temporal integration of the whole system. To better explore temporally extended dynamics, we used two measures common in the study of complex systems: the bivariate transfer entropy [85, 86], and the whole-minus sum integrated information (Φ^*W MS*^) [43, 101]. Both measures provide insight into how how the past of a system constrains the future: in the case of the bivariate transfer entropy, the interaction is local; between individual units (cell-to-cell or region-to-region information transfer), while Φ^*W MS*^ describes how the whole system evolves in ways that are not reducible to a trivial decomposition of the parts.

#### 1. Basal Xenobots show time-directed bivariate information transfer

The bivariate transfer entropy is a classic measure of directed information transfer [85, 86]. Intuitively, the measure quantifies the degree to which knowing the past state of a source element reduces uncertainty about the future state of a target element above and beyond the information disclosed by the targets own past. In this way, it accounts for the first-order autocorrelation in the system to provide a measure of bipartite integration not reducible to spurious correlations between elements. Here, we tested every pair of elements in each fMRI scan and each basal Xenobot recording, using five bins of source history and a single bin of target history to compute the transfer entropy. Each edge was significance tested against a set of 500 circular-shift nulls and retained only if the empirical transfer entropy was greater than all of the null transfer entropies.

The fMRI scans were very dense, with the majority of possible pairwise edges showing significant information transfer (78 ± 4.7%), while the basal Xenobots were considerably less dense (2.8 ± 2.4%). Of the 28 basal Xenobot scans analyzed, five had no significant bivariate transfer entropy at all, while 23 had significant pairwise dependencies. For visualization, see Figure 9A. All fMRI scans had significant information transfer. If we consider those basal Xenobots that did display significant transfer entropy, they showed significantly greater information transfer than their respective nulls: the average information transfer was 0.04 ± 0.01 nat, significantly higher than the null (0.014 ± 0.009 nat, Cohen’s D = 1.81, *p <* 10^*−*^6). The same pattern was true for the fMRI scans, which had an empirical average transfer entropy 4.97 ± 0.037 nat, significantly greater than the nulls, which had average transfer entropy of 1.43 ± 0.15 nat (Cohen’s D = 31.58, *p <* 10^*−*14^. For visualization, see Figure 9B. While we generally do not attempt to compare fMRI scans and basal Xenobots directly, the bivariate transfer entropy is the measure that shows the most striking difference between the two systems. Whether this is due to fundamental differences between the systems, or attributable to differences in data collection (recording duration, number of frames, etc) will be an interesting avenue of future work.

**Figure 9.**
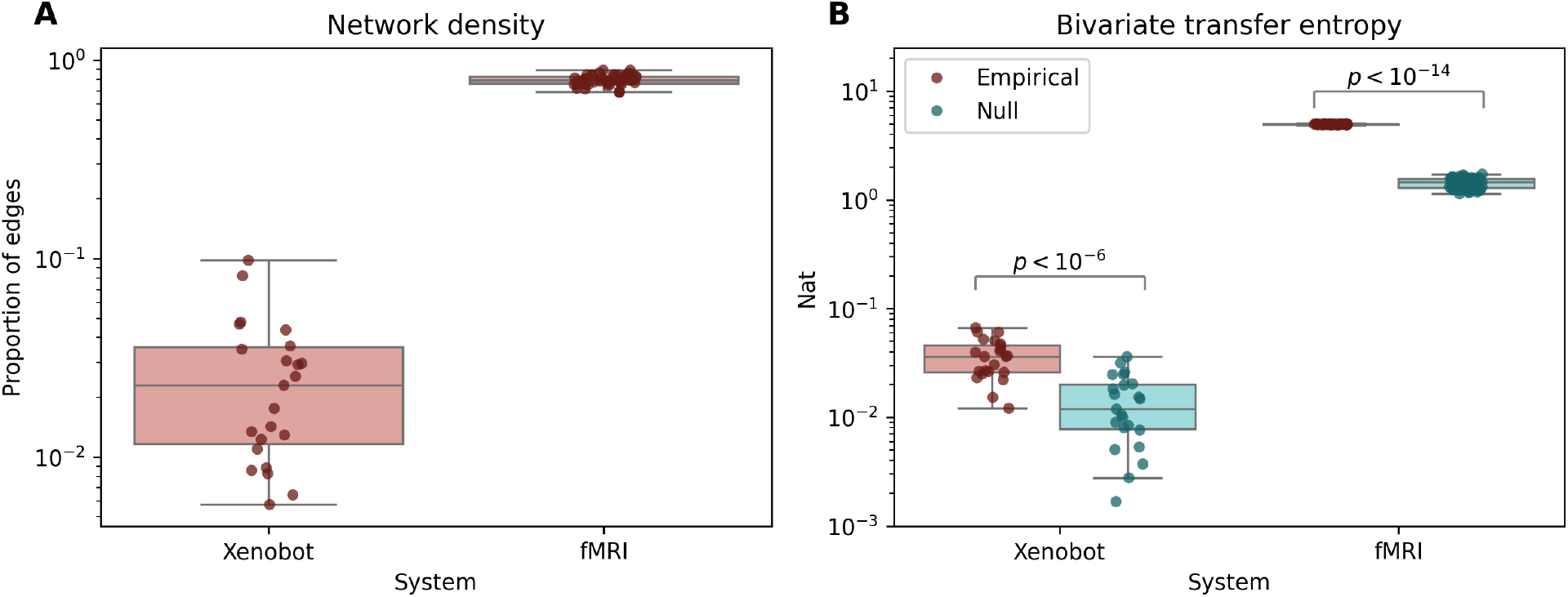
Bivariate transfer entropy. **A.** This plot shows the proportion of possible pairs (i.e. of all possible *X*_*i*_, *X*_*j*_ *∈* **X**) that showed significant transfer entropy. Both the fMRI scans and basal Xenobot recordings showed some multi-cellular directed information flows, although basal Xenobots were considerably sparser. **B**. For both fMRI scans and basal Xenobot recordings, the empirical, significant transfer entropies were significantly higher than their respective circular shifted nulls.

#### 2. Basal Xenobots integrate information over time

The whole-minus-sum integrated information [43, 101] provides a measure of how much more predictive power there is when modeling the system based on its true joint statistics compared to the sum of all the parts modeled independently. Since the whole-minus-sum integrated information can be negative (which occurs when redundant information swamps the higher-order dependency, *a la* the O-information), we computed a corrected version of the integrated information that ensures that the integrated information is non-negative [89], for details see Sec II J. We found that both the basal Xenobots and the brain had significantly greater dynamic integrated information compared to their respective nulls. In the fMRI data, the average integrated information was 0.048 ± 0.04 nat, significantly higher than the associated null 0.011 ± 0.001 (Cohen’s D = 1.22, *p <* 10^*−*11^). In the basal Xenobots, the average integrated information was 0.075 ± 0.098 nat, significantly greater than the null 0.005 ± 0.001 nat (Cohen’s D = 1.02, *p <* 10^*−*9^). For visualization, see Figure 10.

**Figure 10.**
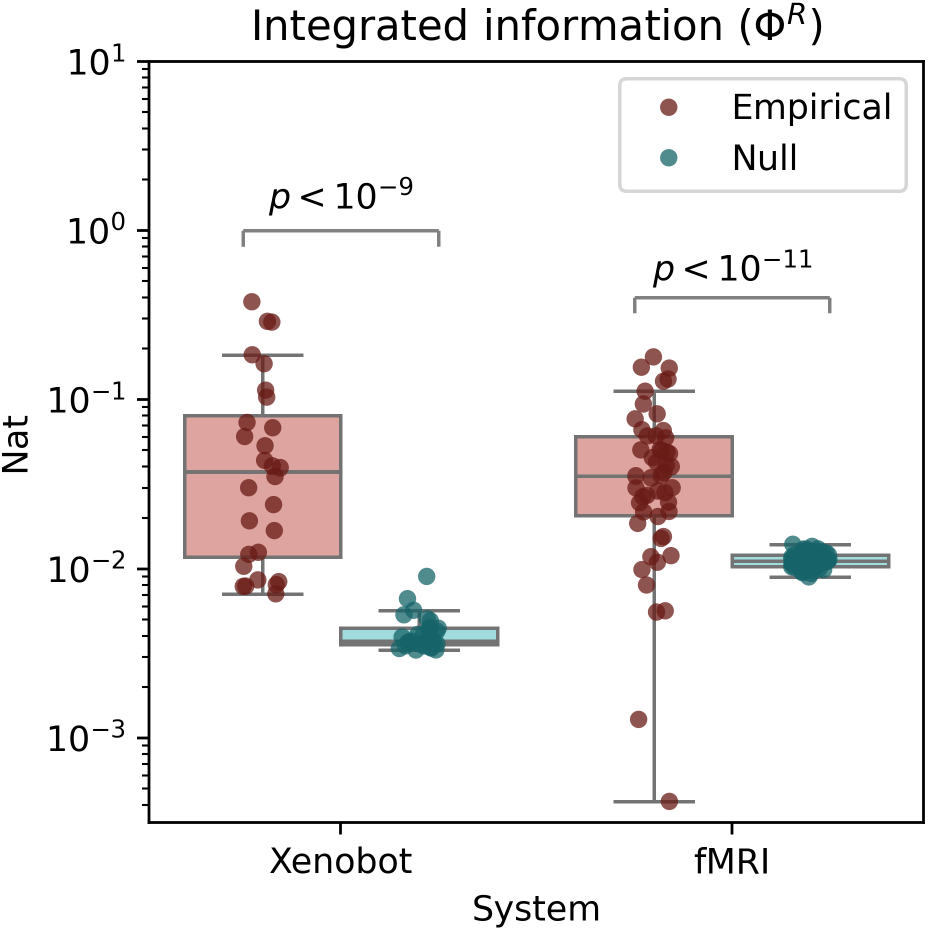
Integrated information (Φ^*R*^. Both the basal Xenobots and the fMRI scans have significantly higher global integrated information than their respective nulls do. This shows that both systems display at least some dynamical properties in the “whole” that are not trivially reducible to lower-order “parts.”

All of these results have fit a common pattern: whatever the measure of structure applied, both the brains and basal Xenobots show significantly greater structure than a null model that preserves lower-order dynamics but disrupts higher-order ones. This suggests that there is a large number of “features” of structural organization that are shared by both basal Xenobots and brains, despite their radically different biological substrates.

## IV. DISCUSSION

In this paper we have compared two radically different biological systems: adult human brains (as imaged with fMRI [49]) and preparations of embryonic frog epithelial tissue colloquially named “basal Xenobots” (expressing a fluorescent calcium reporter). In neuroscience, many of the organizational features we tested are known to be associated with meaningful differences in cognitive or clinical states (such as anesthesia, aging, and neurodegeneration), and so the question of how to interpret their presence in non-neural systems is an intriguing one. While changes in information flows in brains track changes in cognitive state, we do not claim that the presence of similar flows in basal Xenobots is necessarily indicative of “cognition” (to the degree it is seen in brains). Clearly information processing patterns and cognitive processes are correlated in brains, but any stronger claim of identity would be inappropriate. We hypothesize, instead, that these signs of similarity reflect a deeper, underlying commonality between brains and basal Xenobots, which could be probed with techniques from complex systems science, network science, and multivariate information theory. Morphogenetic systems, whether natural embryos or synthetic constructs, are distinguished by coherent, coordinated actions of individual cells toward specific outcomes, often featuring the ability to attain the target morphology despite noise or even drastic interventions. Like neurons in a brain which implement an integrated “animal” with competencies, preferences, and memories that belong to it and not to any of its parts, biochemical and bioelectrical networks among cells likewise serve a binding function [102, 103]. They enable cellular collectives to traverse the space of anatomical possibilities as a coherent unit, with all cells cooperating to achieve a large-scale outcome (e.g., the right number of fingers in a regenerating axolotl limb) that is not defined at the level of a single cell. Thus, we sought to use established methods from neuroscience to attempt to quantify the ability of non-neural cells to process information in a way that reveals a whole distinct from a collection of parts. Given the ubiquity of these self-organizing processes throughout the natural world, it seems unlikely that the patterns described here would be unique to brains and basal Xenobots: if they do, in fact, reflect conserved capacity to self-organize collectives into coherent structures, we would expect to see them in many different tissues and organisms.

We found that, across a battery of measures of statistical organization, including networks, higher-order interactions, dynamic information integration, and time-resolved fluctuation analysis that basal Xenobots display patterns of information that resemble those seen in human brains, and are universally more organized than a null model that preserves cell-level dynamics. As we will discuss below, these findings raise questions about how to think about modeling both neural and non-neural complex biological systems, and challenge how we interpret many of the statistics that have formed a core part of modern computational neuroscience.

The use of “brain-like” in this paper may seem provocative. There are, of course, a tremendous number of ways that basal Xenobots are nothing like human brains: they are different scales, cultured from different physiological systems so the cell types are completely different, and the behaviors each system can display are profoundly different as well. Despite these differences, however, we argue that the claim of brain-likeness remains insightful: almost all of the features explored here have been found to correlate with clinically or cognitively meaningful differences in mental state or behavior. The finding that they are also present in basal Xenobots raises complex philosophical and scientific questions about the nature and purpose of these patterns in Nature. All of these statistics were developed to reveal different kind of “complex” structure: organizational architectures involving the integrated interaction of many different components, largely independent of the specific nature of the individual elements [11]. This is what has motivated the applications in neuroscience, but in theory the notions of information-sharing and dependency revealed by these measures should apply to any system. If we propose that, in brains, these patterns are reflective of fundamental features of the brain’s central function, then we have to ask: what are they doing in basal Xenobots? Have we underestimated the information processing capacity of non-neural tissue? Is it possible that the fundamental machinery of cognition is more widespread than it appears and can be instantiated in non-neural tissue, devoid of spikes, neurotransmitters, and the usual biological “wetware” we assume is required for coherent behavior? Possibly. Conversely, one could argue that the presence of these patterns in (commonly-assumed to be) non-cognitive, non-conscious embryonic tissue suggests that the neuroscientists exploring these patterns in brains have been chasing statistical epiphenomena; that collective information, meso-scale modularity, and integration/segregation balance do not actually reveal anything uniquely specific to the brain at all. If this is the case, it would behoove neuroscience as a field (particularly theoretical and computational neuroscience that relies on gross-scale neuroimaging) to reflect on what it is that is being measured, and why. To tackle these questions will require considerably more detailed analysis of information dynamics in non-neural tissue, as well as more formal and rigorous definitions of what “cognition” or “cognitive processes” might look like in biological systems vastly different from the usual animal models used in neuroscience and cognitive science. The latter is the province of the emerging field of diverse intelligence [47, 48, 104].

We have no answer to these questions at present, however there these results undeniably show that basal Xenobots are themselves “complex systems”. While we remain largely agnostic about the interpretation of these results with regards to philosophically fraught topics such as “cognition”, it is clear from these results that the basal Xenobots themselves are instantiating rich and sophisticated patterns of cell-to-cell, and higher-order multi-cell coordination; patterns that could not have occurred due to chance, or due to lower-order features such as temporal autocorrelation of the individual cell time series. While this may seem obvious in retrospect, given the nature of the preparation of the Xenobots (explanting and treating embryonic epithelial tissues in disruptive and non-natural ways), it was far from given that the same kinds of sophisticated, multi-scale systems that emerge in typical development would be preserved after construction of the basal Xenobot. We can, however, say with confidence that such structure can, and do, emerge, even in these artificial preparations. The question of *how* they emerge, and by what channels cells in the basal Xenobots use to faciliate information-exchange remains mysterious, however.

The emergence of complex, non-trivial forms of statistical integration has been documented in many models of the brain as a complex system, however, these models almost universally assume that the dynamics are being run on a physical substrate of white-matter connections (e.g. see [66, 105], which used a Kuramoto model on structural graphs derived from MRI scans of white-matter tracts to explore the relationship between structural topology and functional dynamics). A question for future researchers is to what extent the basal Xenobots show a similar structural network, and whether the patterns of functional connections track the patterns of structural connections in ways similar to what is observed in the brain. It is known that collections of cells can have structural, as well as functional connections: tunneling nanotubes can connect distinct cells [106–108], and can be and reach lengths as long as multiple cell diameters [109], facilitating everything from electrical connectivity to organelle transfer. Unlike white matter connections in the brain, which are reasonably static on short to medium timescales (such as the duration of an fMRI scan), tunneling nanotubes are fragile and dynamic, and also much harder to image, making a “structure/function” analysis currently impossible. Other methods of physical cell-to-cell communication might include exosomes: vesicles containing signaling molecules that are secreted by cells into the extra-cellular space for take-up by (potentially) distant neighbors [110]. Unlike nanotubes, which form one-dimensional links between source cells and target cells, exosomes can diffuse, potentially reaching many cells simultaneously and thereby facilitating one-to-many, or broadcaster, styles of communication that may be better modeled with directed hypergraphs or other higher-order generalizations of the classic functional connectivity network [28].

Finally, this work sets the stage for future work on how perturbations can impact the structure of emergent coordination in non-neural tissue. A key conceptual driver of this work is the repeated finding that, in the brain, perturbations and state differences, such as anaethesia, brain injury, aging, neurodivergent developmental differences, and more are reflected in changes to functional connectivity structures [33, 35, 44, 45]. We expect that the same thing will be true in basal and modified Xenobots: that different perturbations or chemical environments will give rise to distinct patterns of global cell-cell coordination. Possible interventions to explore might include anaesthetics, inflammatory signaling molecules such as cytokines, purinergic singaling molecules such as ATP and ADP, or mechanical per-turbation (as was done in [53]). Currently, the results of this paper show that these patterns exist, but shed little light on exactly what they mean or how to interpret them: follow-up work detailing how they change will go a long way to making it clear what it is we are observing here.

## V. CONCLUSIONS

Here, we showed that two radically different biological systems: human brains, and basal Xenobots (self-assembling epithelial cell constructs) display a common set of organizational features and dynamics, including sophisticated functional connectivity networks, higher-order information of multiple types, and dynamic integration of information. In human brains, these features have been associated with meaningful differences in cognitive and behavioral state, raising questions about how to interpret their presence in the context of non-neural tissue. Despite their comparatively simpler make-up, and the artificial processes involved in making them, these results show that basal Xenobots are undeniably complex systems in their own right, displaying a rich information structure and emergent organizational features. Given the lack of (apparent) structural connectivity (a key difference from brains), basal Xenobots displaying complex, emergent statistical structures may have alternative ways of propagating signals and processing them. We propose that these results show that “brain-like” patterns of information processing may not be specific to the brain at all; that ideas like “information processing” and “information integration”, may be even more relevant to non-neural, biological systems than is generally appreciated and the functional similarities between different types of embodied autonomous agents can now be quantified.

## Acknowledgments

This work was supported by the Cold Regions Research and Engineering Laboratory (CRREL) under Contract No. W913E524C0012.

