## Supplemental material for "Identification of brain-like complex information architectures in embryonic tissue of *Xenopus laevis* organoids"

### Time-lagged conditional total correlation

The definition of the total correlation [1] given in the manuscript assumes that every frame of each recording is an independent draw from some high-dimensional probability distribution $P(\mathbf{X})$, which in the Gaussian case is parameterized by the covariance matrix $\Sigma_{\mathbf{X}}$:

$$TC(\mathbf{X}) = \sum_{i=1}^{N} H(X_i) - H(\mathbf{X}) \tag{1}$$

To ensure that the higher-order integration was robust to this assumption, we replicated the analysis using the time-lagged, conditional total correlation rate.

Recall that the conditional entropy rate is defined by [2]:

$$H_\mu(X^t) = H(X^t|X^{t-1}) \tag{2}$$

This quantifies how much uncertainty about the next state of $X$ remains after learning the past state. In systems with non-trivial autocorrelation $H_\mu(X^t) < H(X^t)$.

We can extend the conditional entropy rate to the conditional total correlation rate:

$$TC_\mu(\mathbf{X}) = \sum_{i=1}^{N} H(X_i^t|\mathbf{X}^{t-1}) - H(\mathbf{X}^t|\mathbf{X}^{t-1}) \tag{3}$$

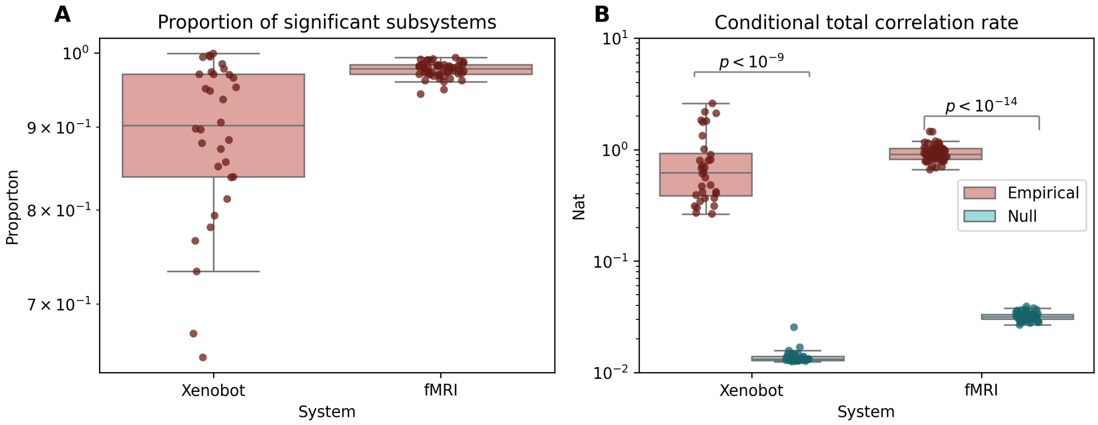


Fig. 1: **Total correlation rate. A.** The proportion of samples of size 3, 4, or 5 elements that have significant conditional total correlation rate for basal Xenobots and fMRI scans. Note that the proportion is much higher than for just the total correlation for both basal Xenobots and fMRI. **B.** Both basal Xenobots and fMRI scans show greater empirical conditional total correlation rate than their respective circular-shifted nulls.

If Figure 1 we can see that the same pattern is present: significantly greater integration in the empirical data compared to the autocorrelation-preserving nulls.

### Comparing TC and conditional TC rate.

If the apparent total correlation in $\mathbf{X}^t$ is attributable to the global state of the system at time $t-1$, then the conditional $TC(\mathbf{X}^t|\mathbf{X}^{t-1})$ should be less than $TC(\mathbf{X})$. We can see this in Figure 2, which plots CDF curves of the TC and conditional TC rate over all samples from all xenobots, as well as their associated nulls. In the case of empirical data (Panel A), the TC rate curve (orange) is greater than the TC curve. In contrast, for the circular-shifted null data, the TC rate curve is less than the TC curve.

This indicates that, in the null, circular-shifted data, any deviation from independence is explainable by autocorrelation (as expected, as the circular shift null disrupts all inter-cell dependencies). In contrast, in the empirical data, knowing the past actually illuminates *more* integration than can be identified if we model the data as independent draws from some $P(\mathbf{X})$.

The difference between the TC and the conditional TC rate quantifies how much more (or less) information that the past of the whole ($\mathbf{X}^{t-1}$) discloses about it's own future ($\mathbf{X}^t$), or the sum of it's constituent parts:

$$TC(\mathbf{X}^t) - TC(\mathbf{X}^t|\mathbf{X}^{t-1}) = \sum_{i=1}^{N} I(\mathbf{X}^{t-1}; X_i^t) - I(\mathbf{X}^{t-1}; \mathbf{X}^t) \quad (4)$$

As with the O-information, this measure can be signed, and a negative sign indicates greater information in the "whole" than the "sum of its parts" (synergy), while a positive sign indicates the opposite (redundancy).

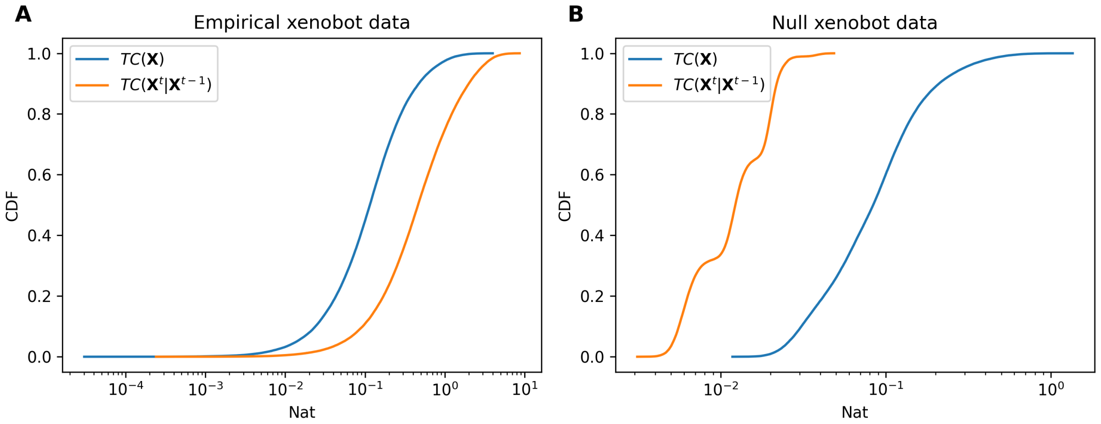


Fig. 2: **TC vs. TC rate. A.** Cumulative distribution plots for the total correlation (blue) and the conditional total correlation rate (orange) for empirical xenobot data (aggregated over all significant samples from all bots). **B.** The same as panel A, except for the circular-shifted nulls.

### Visualization of empirical and null covariance matrices

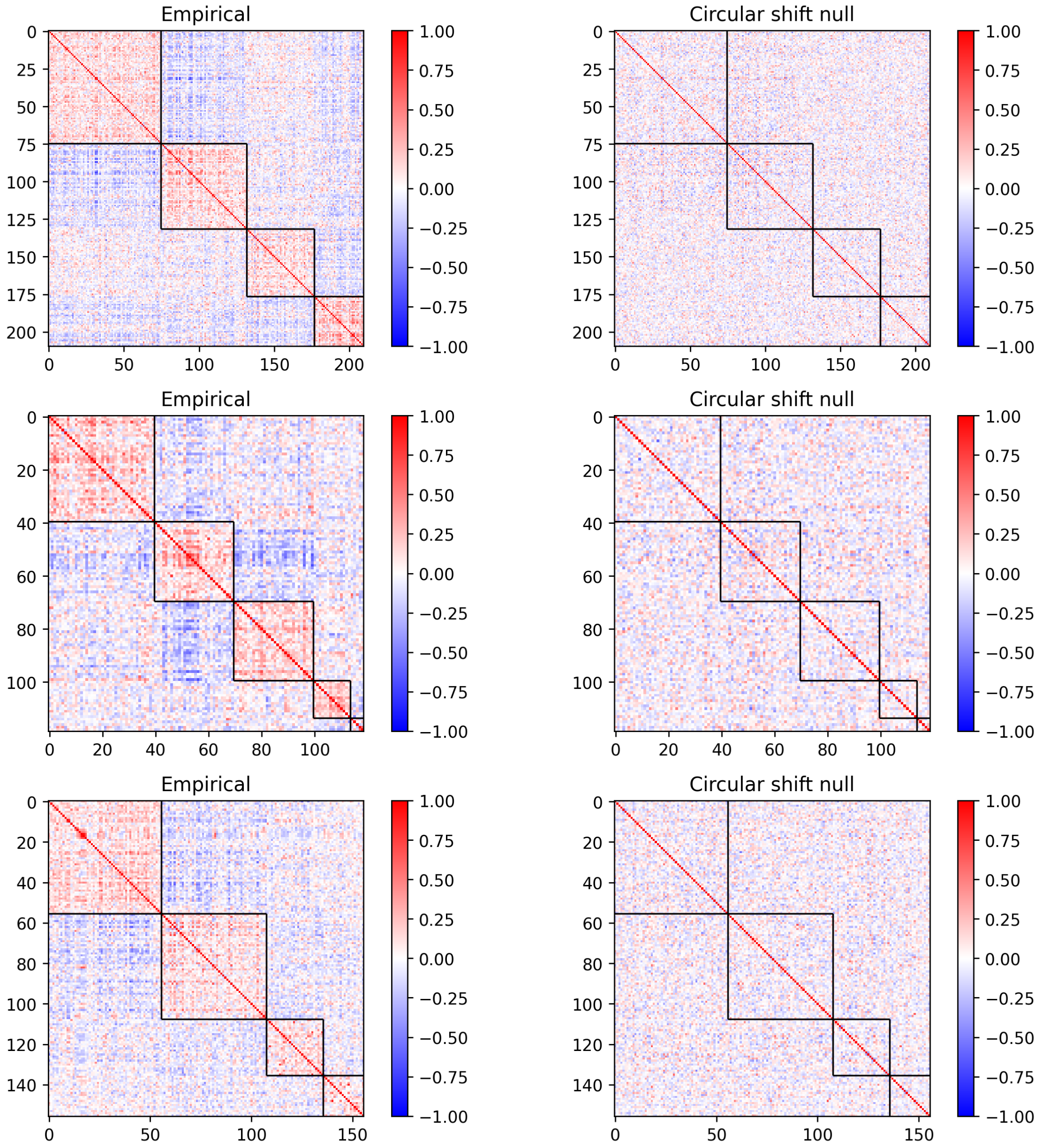


Fig. 3: **Null covariance matrices.** Empirical, clustered example of three empirical covariance matrices ordered into functional clusters (right column). Examples of null covariance matrices (left column) for each data set after applying a circular-shift null. Note that, while overall covariances are lower in magnitude and have lost the modular structure, there are still non-zero functional edges: this is the spurious dependency caused by first-order features such as autocorrelation.

**Modules in functional connectivity networks.**

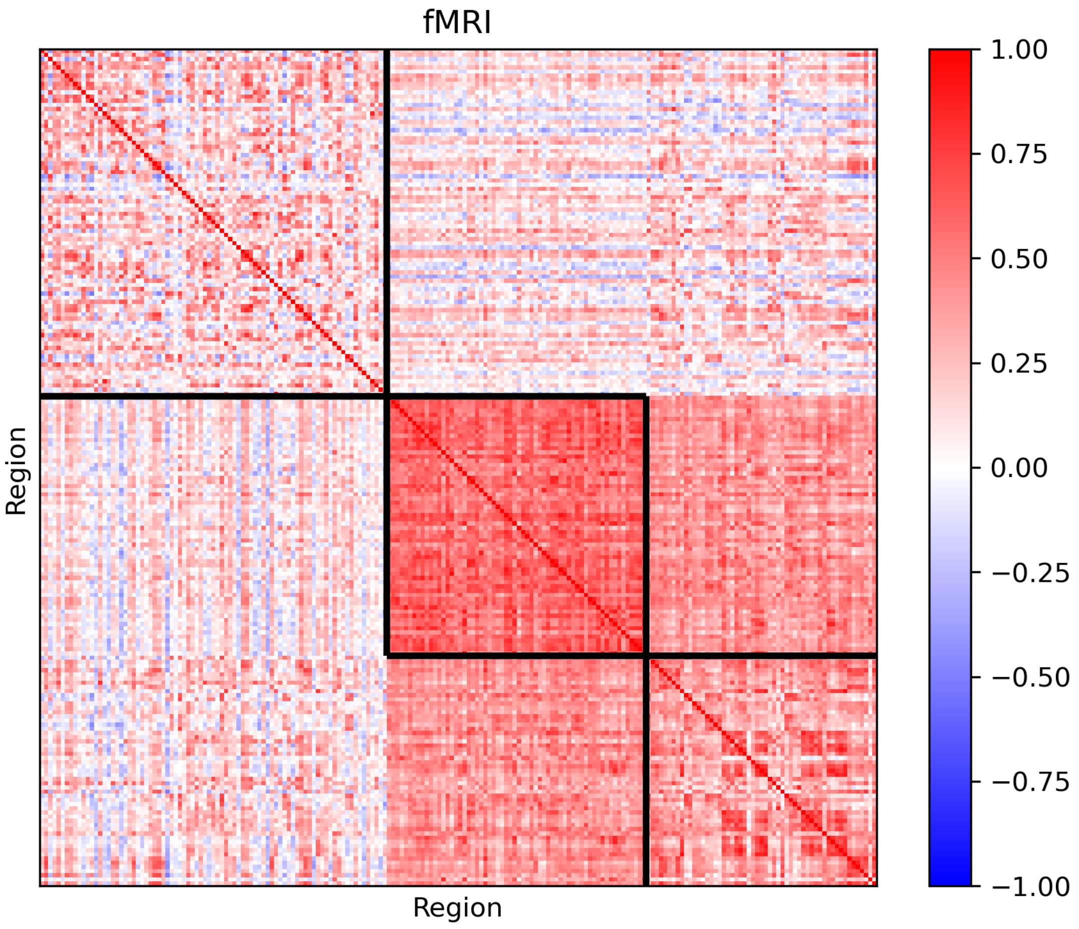
fMRI
Region
Region
1.00
0.75
0.50
0.25
0.00
−0.25
−0.50
−0.75
−1.00

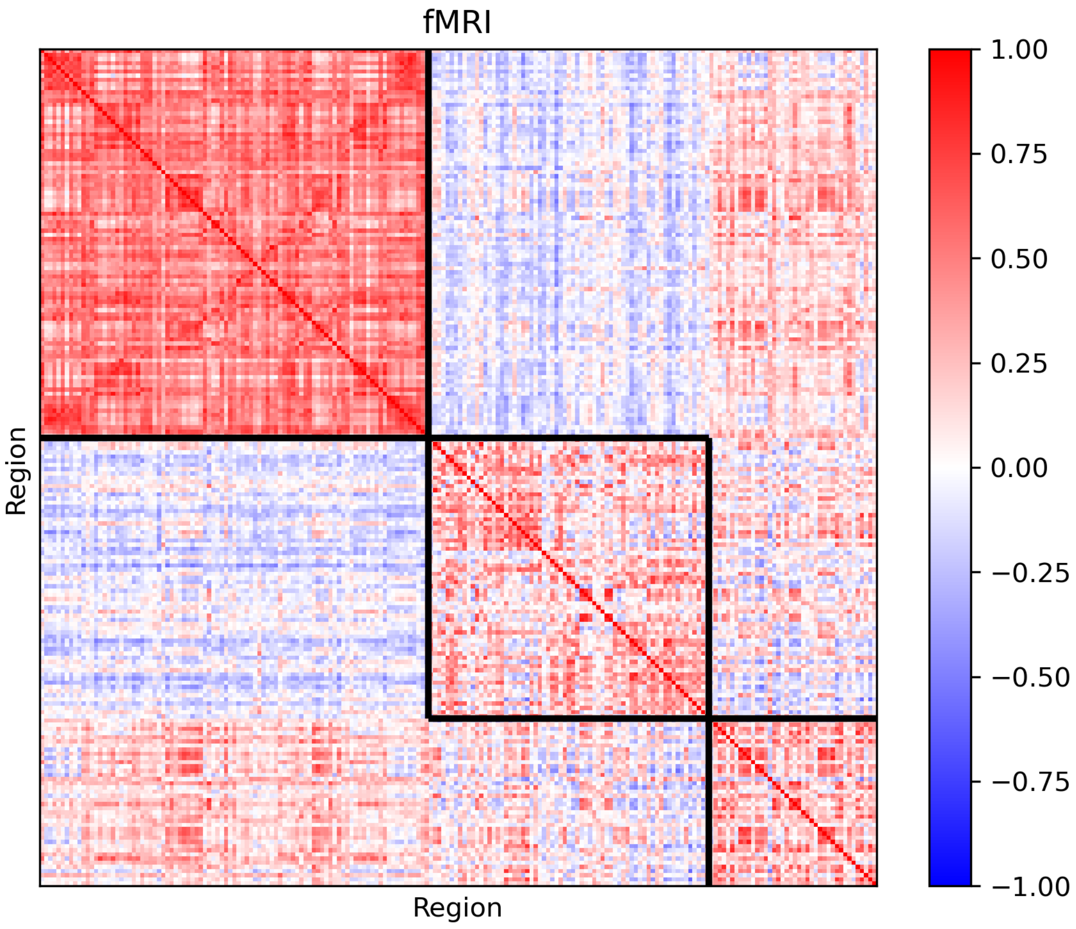
fMRI
Region
Region
1.00
0.75
0.50
0.25
0.00
−0.25
−0.50
−0.75
−1.00

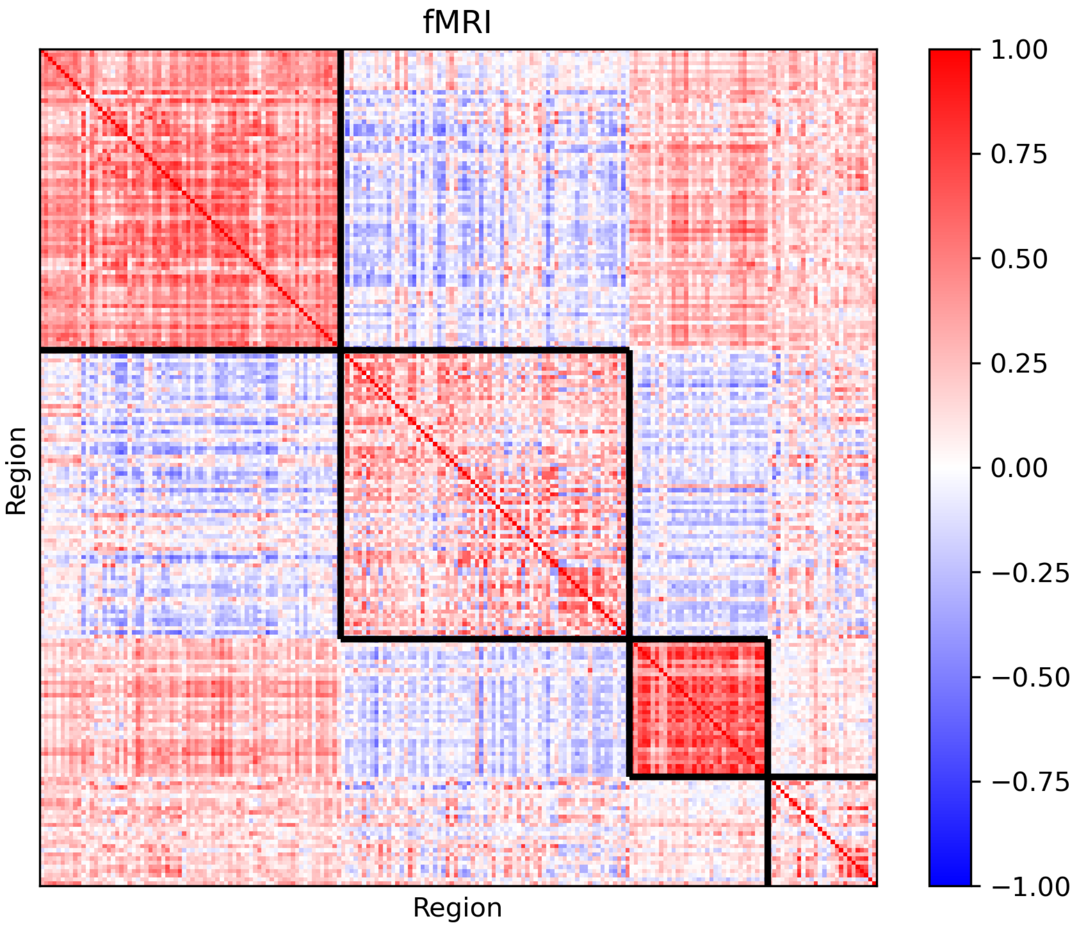
fMRI
Region
Region
1.00
0.75
0.50
0.25
0.00
−0.25
−0.50
−0.75
−1.00

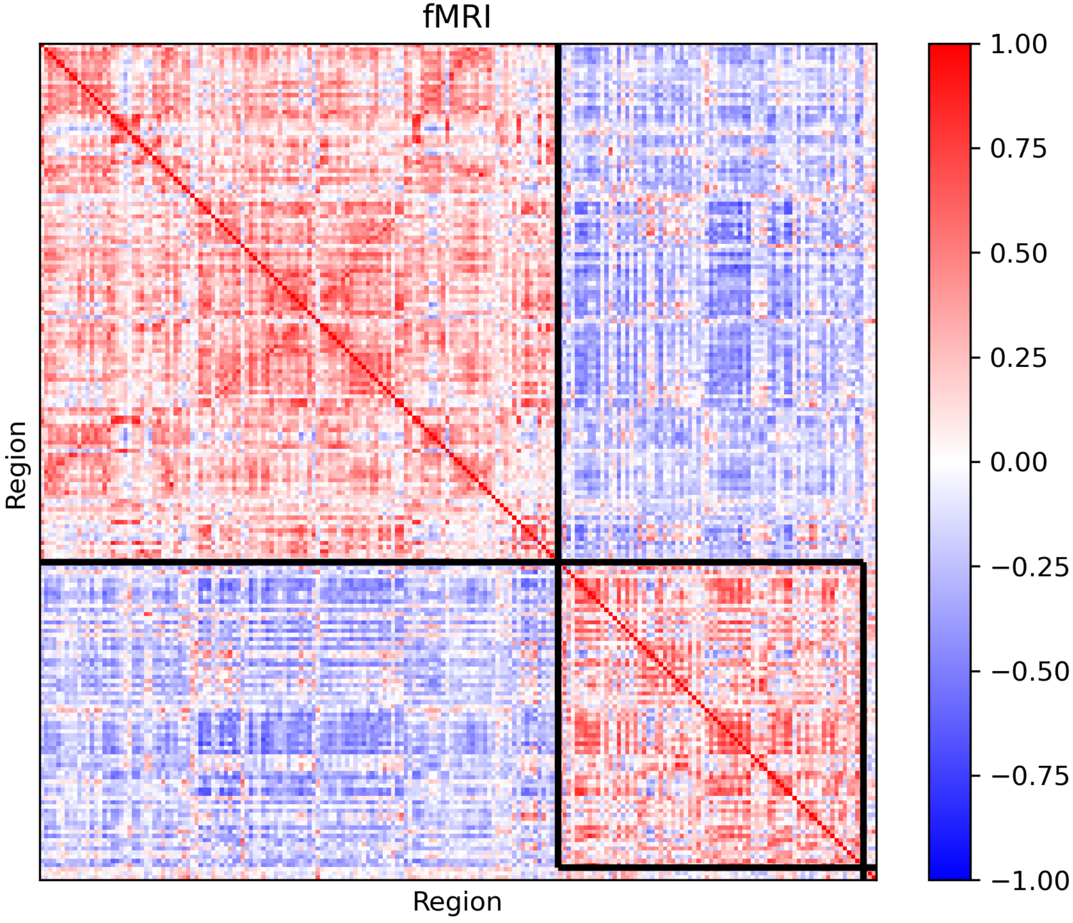

fMRI
Region
Region
1.00
0.75
0.50
0.25
0.00
−0.25
−0.50
−0.75
−1.00

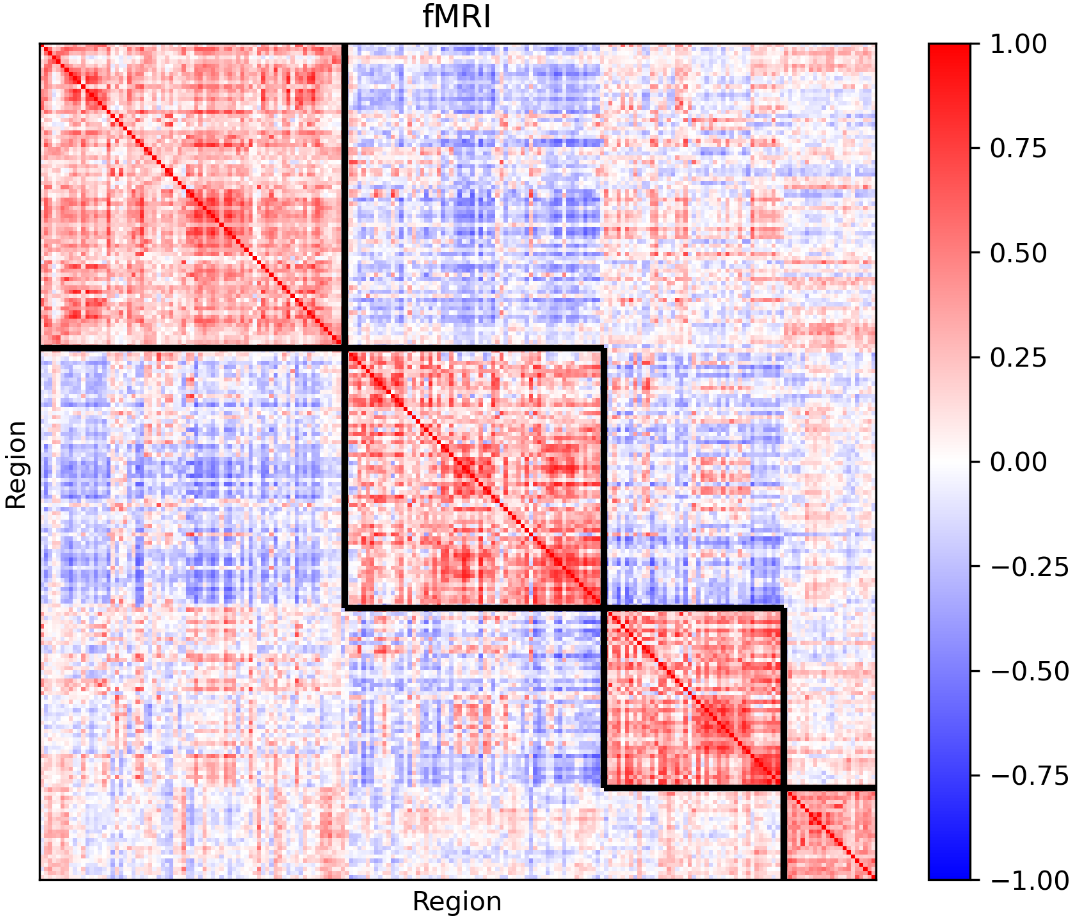
fMRI
Region
Region
1.00
0.75
0.50
0.25
0.00
−0.25
−0.50
−0.75
−1.00

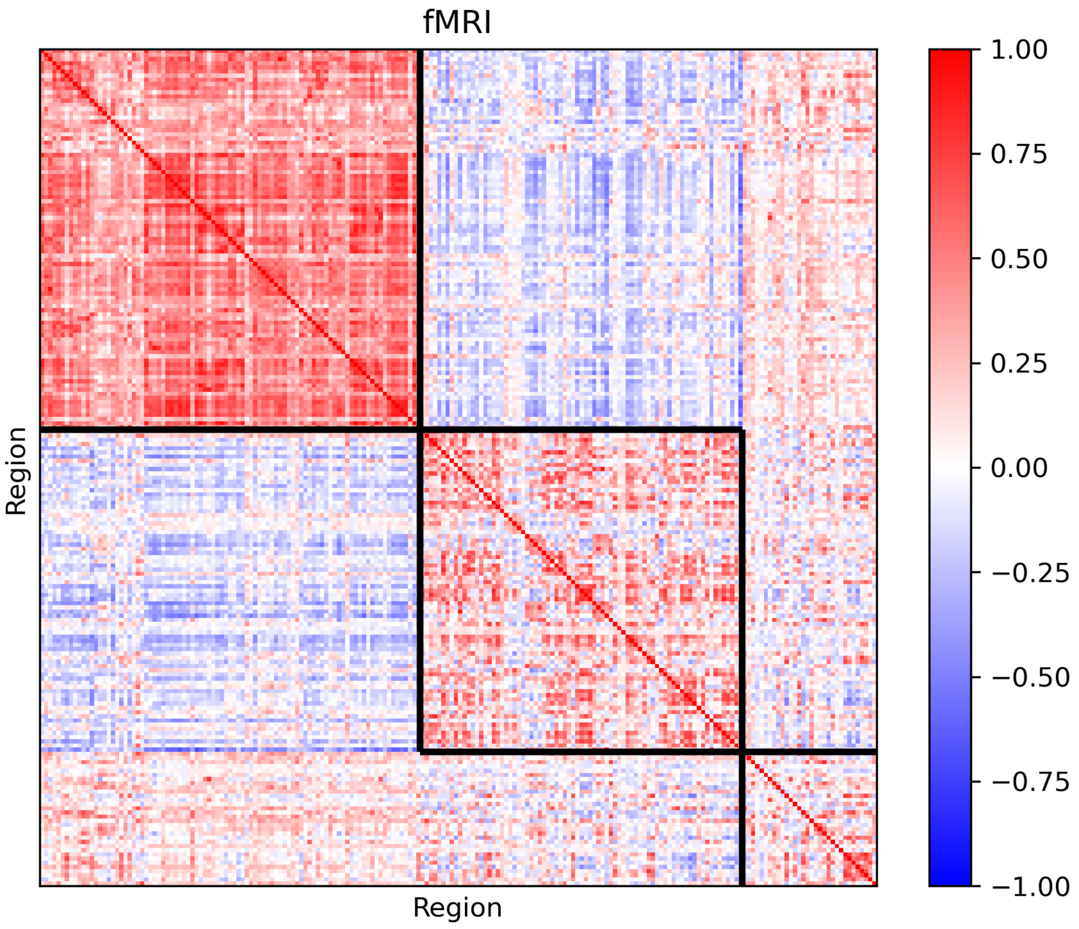
fMRI
Region
Region
1.00
0.75
0.50
0.25
0.00
−0.25
−0.50
−0.75
−1.00

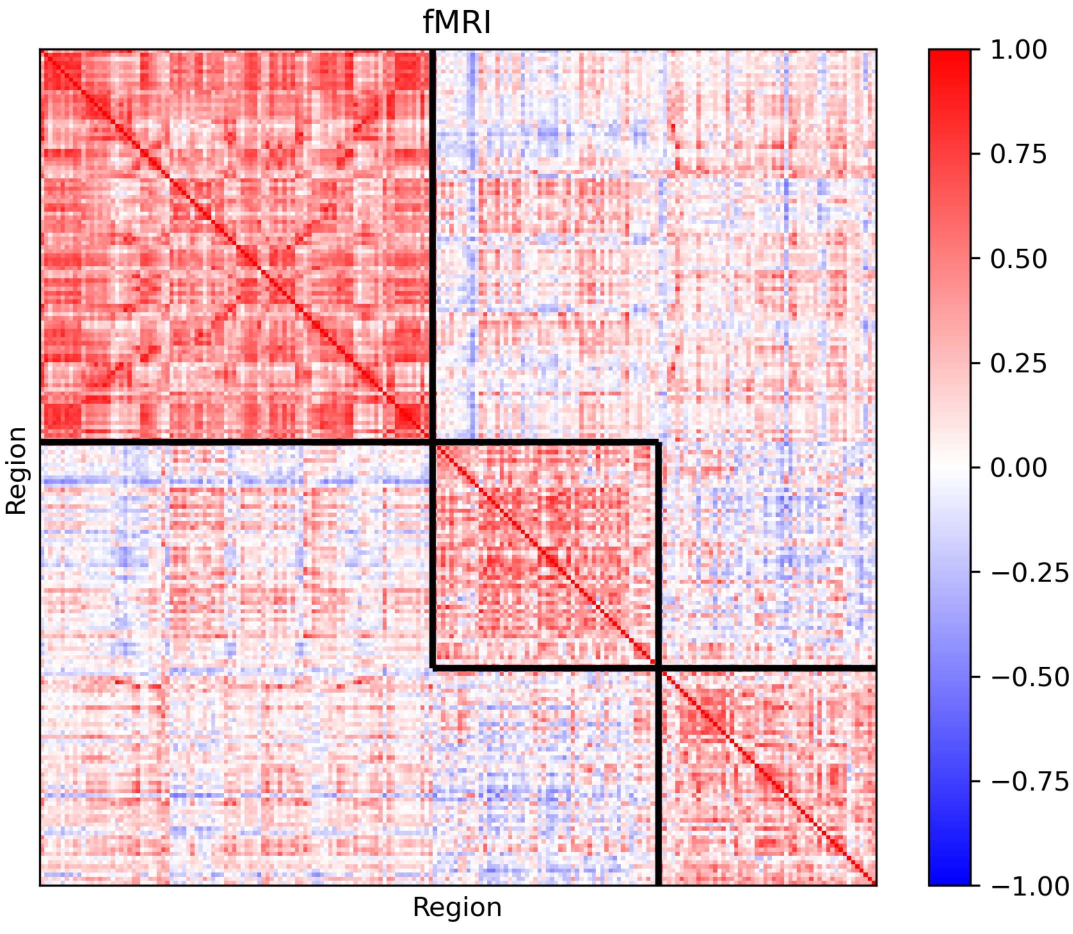
fMRI
Region
Region
1.00
0.75
0.50
0.25
0.00
−0.25
−0.50
−0.75
−1.00

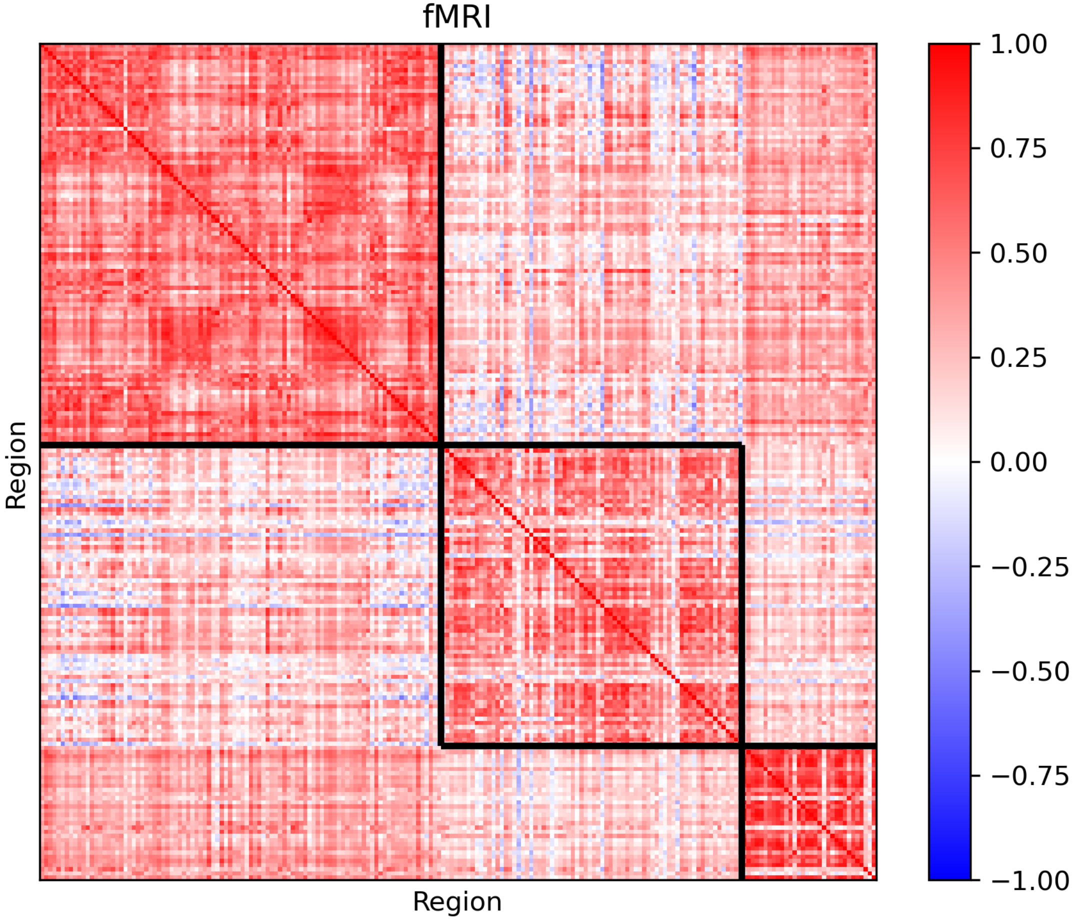
fMRI
Region
Region
1.00
0.75
0.50
0.25
0.00
−0.25
−0.50
−0.75
−1.00

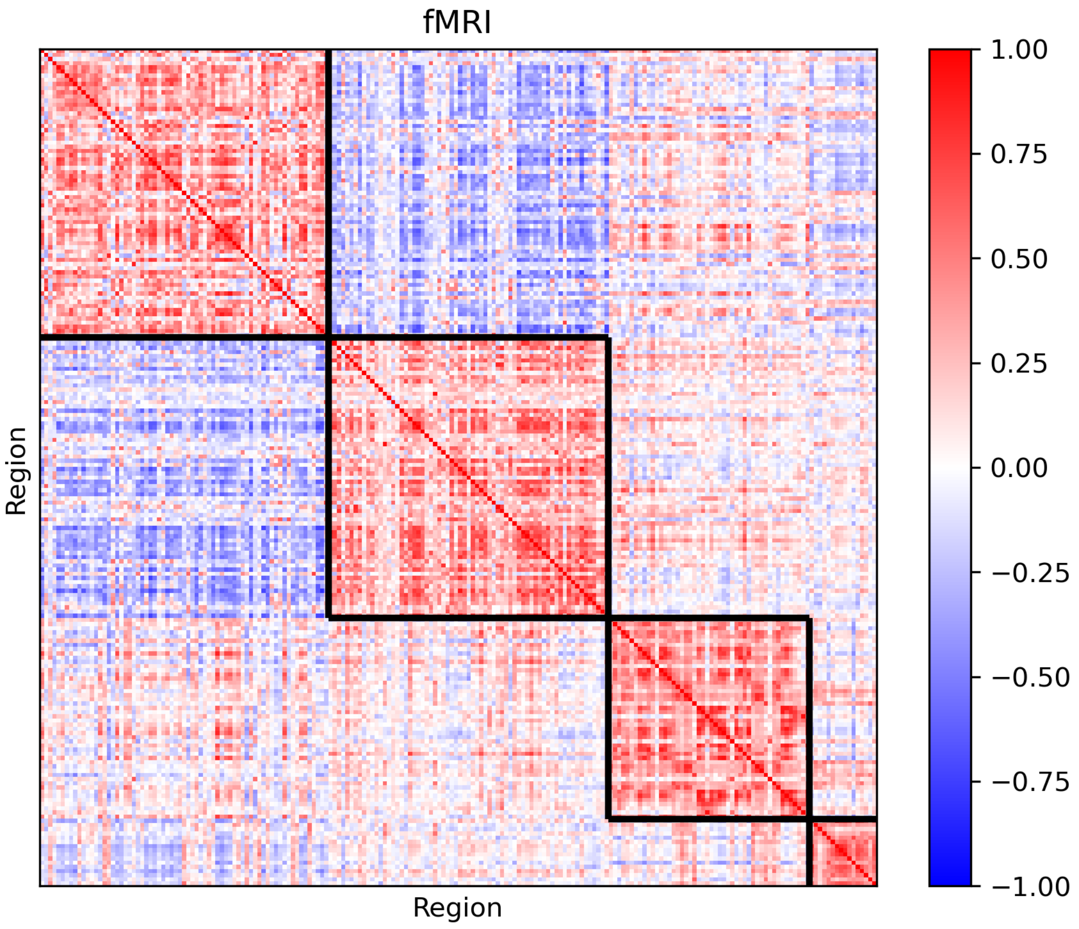

fMRI
Region
Region
1.00
0.75
0.50
0.25
0.00
−0.25
−0.50
−0.75
−1.00

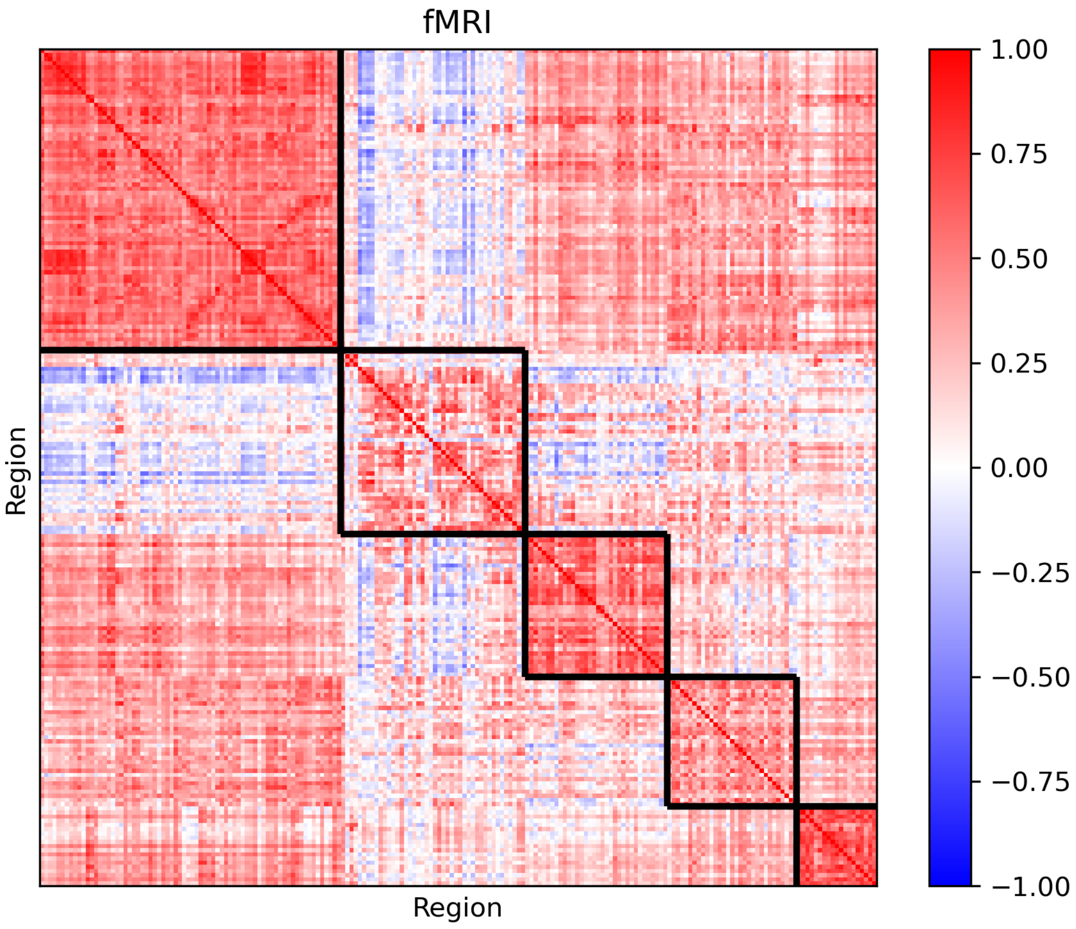
fMRI
Region
Region
1.00
0.75
0.50
0.25
0.00
−0.25
−0.50
−0.75
−1.00

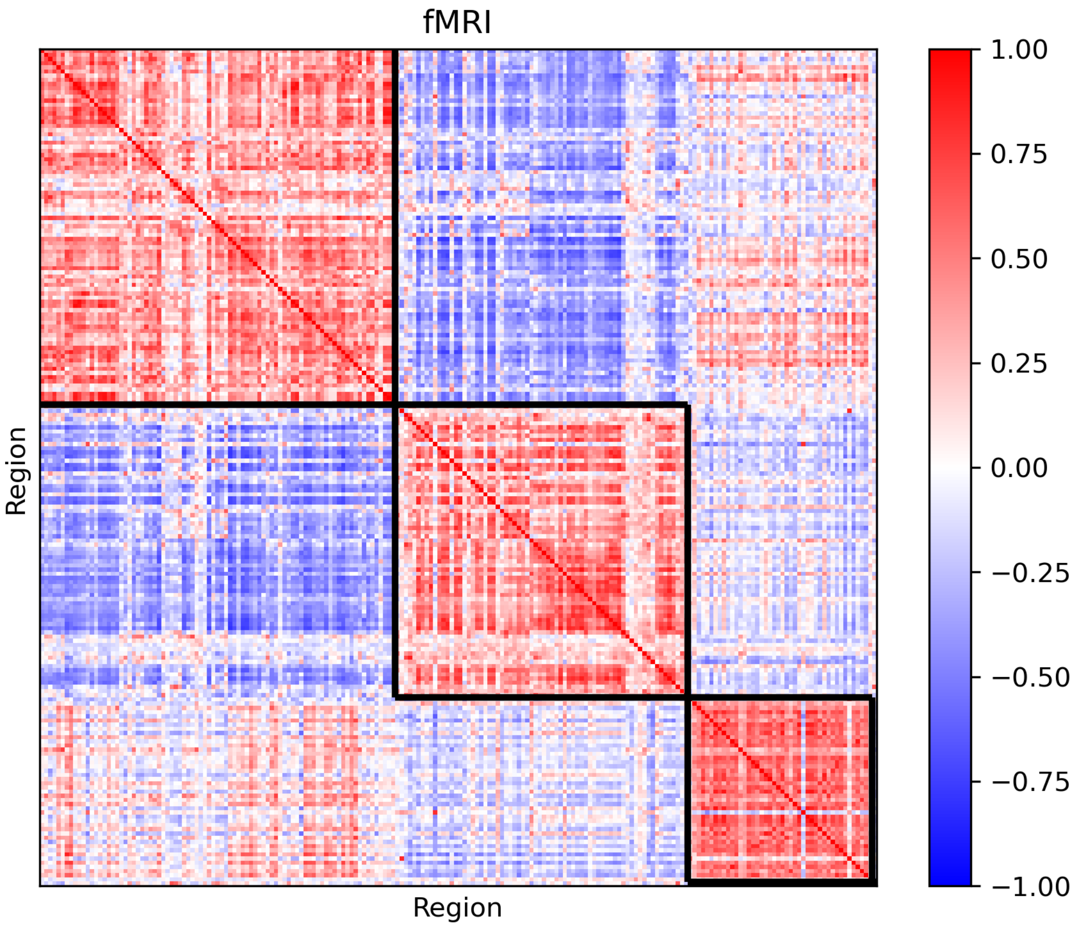
fMRI
Region
Region
1.00
0.75
0.50
0.25
0.00
−0.25
−0.50
−0.75
−1.00

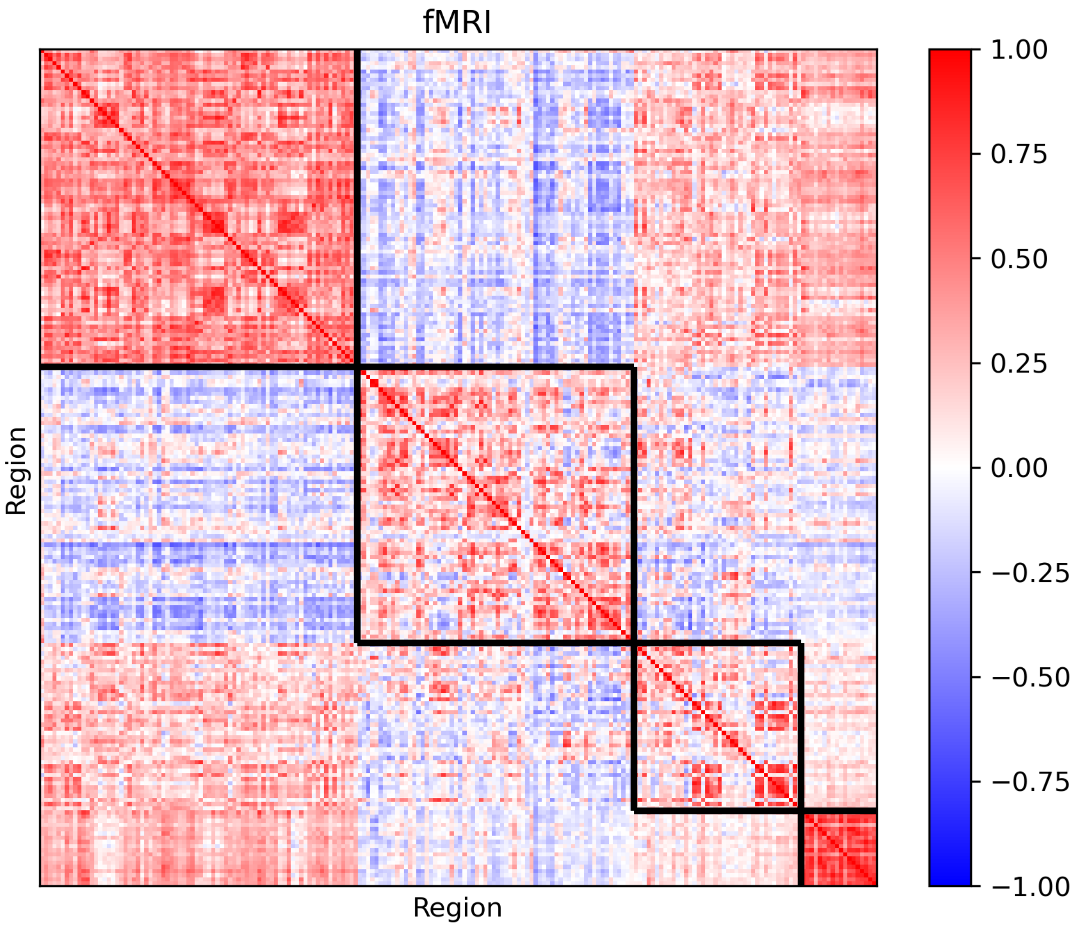
fMRI
Region
Region
1.00
0.75
0.50
0.25
0.00
−0.25
−0.50
−0.75
−1.00

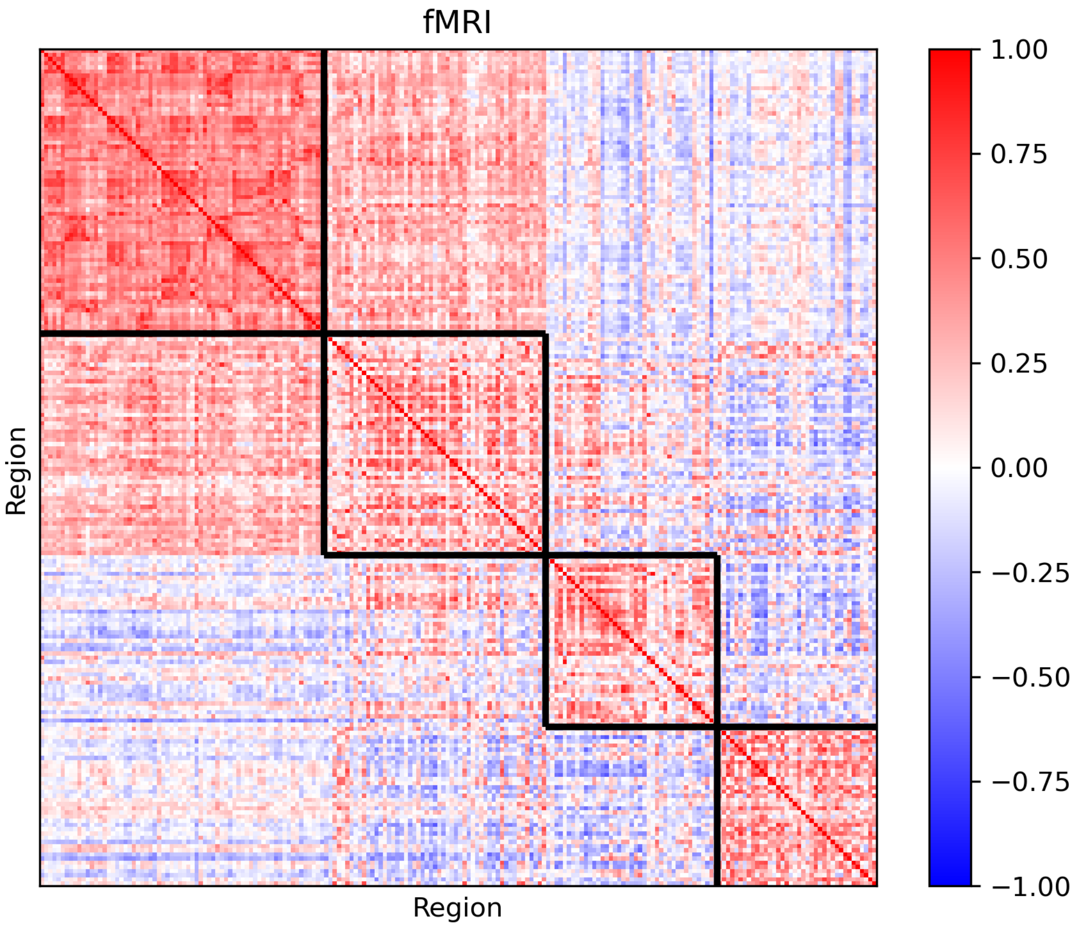
fMRI
Region
Region
1.00
0.75
0.50
0.25
0.00
−0.25
−0.50
−0.75
−1.00

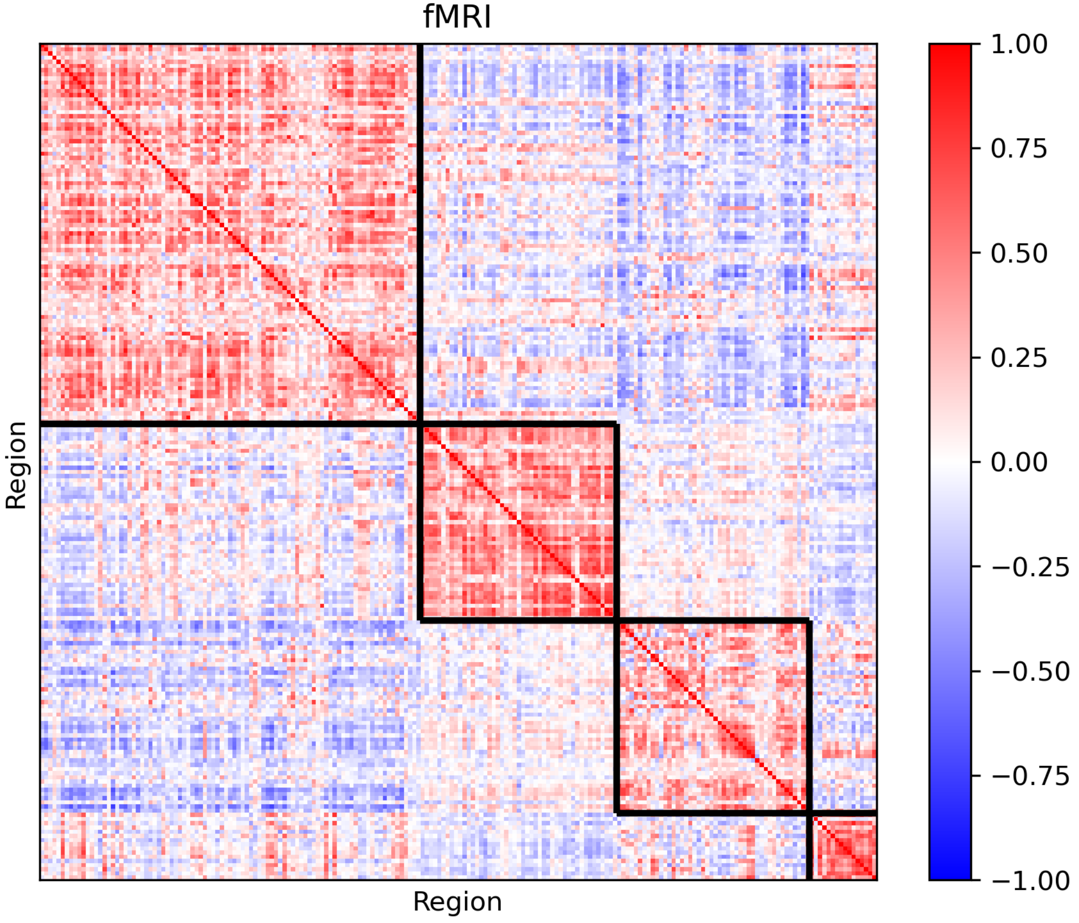
fMRI
Region
Region
1.00
0.75
0.50
0.25
0.00
−0.25
−0.50
−0.75
−1.00

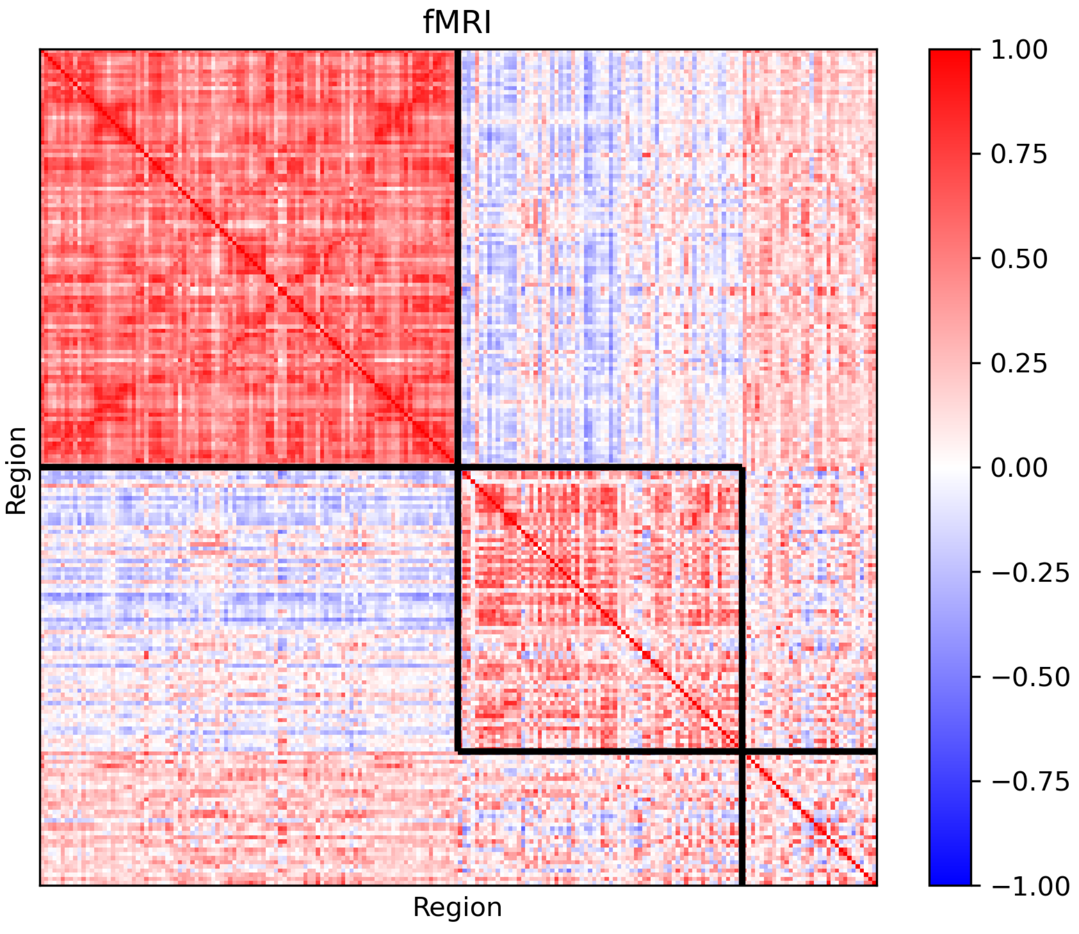
fMRI
Region
Region
1.00
0.75
0.50
0.25
0.00
−0.25
−0.50
−0.75
−1.00

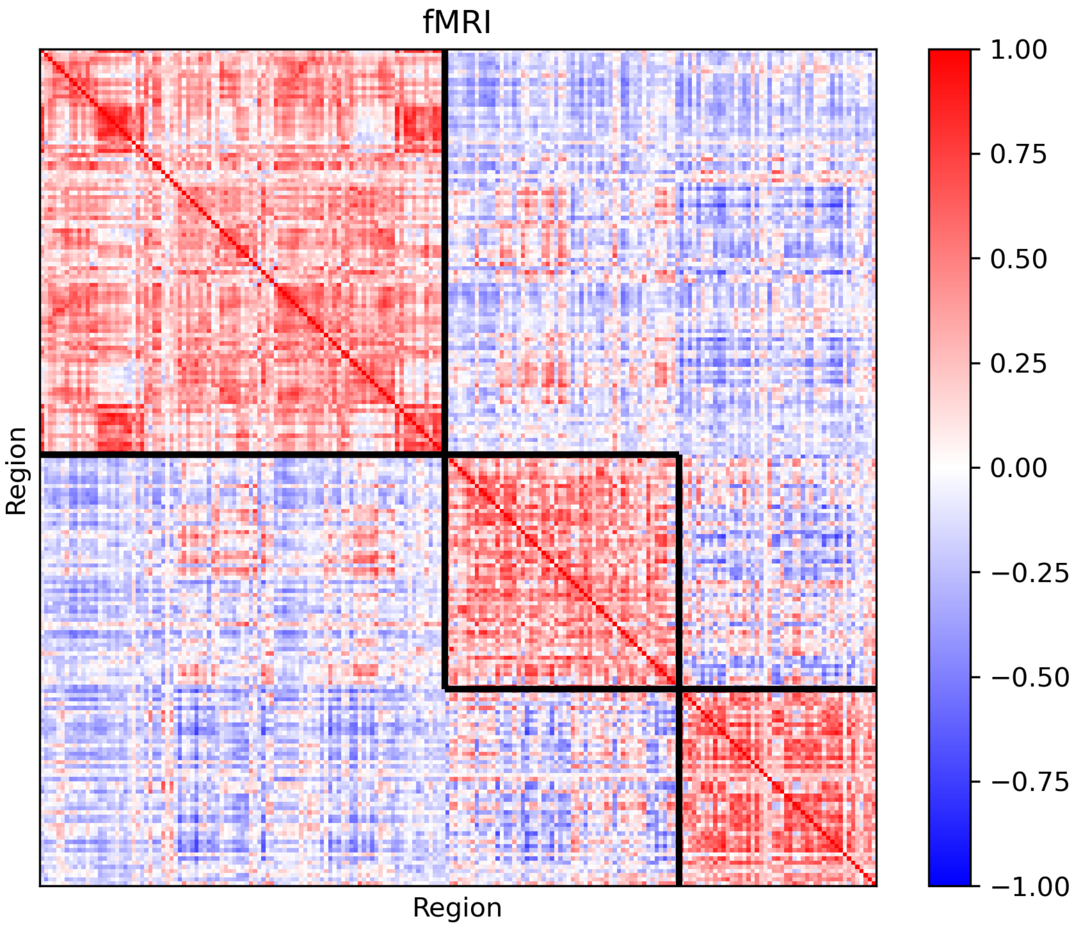

fMRI
Region
Region
1.00
0.75
0.50
0.25
0.00
−0.25
−0.50
−0.75
−1.00

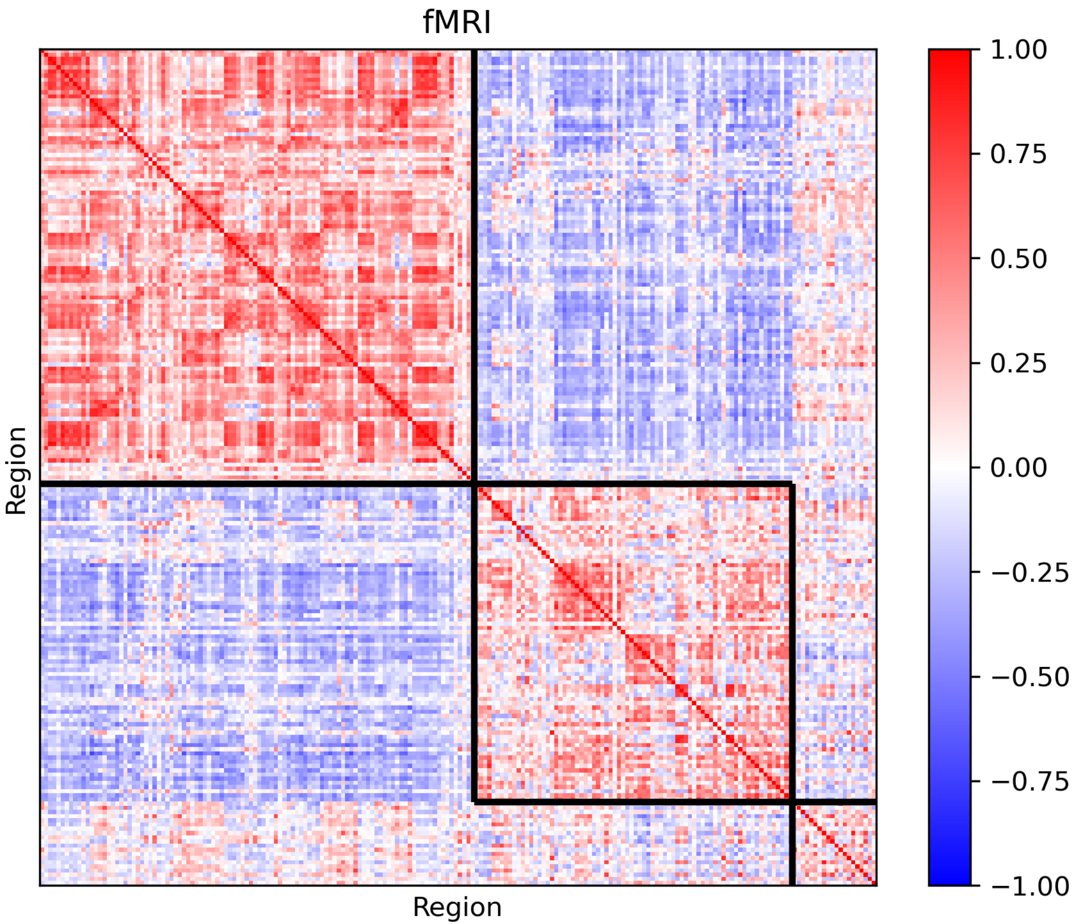
fMRI
Region
Region
1.00
0.75
0.50
0.25
0.00
−0.25
−0.50
−0.75
−1.00

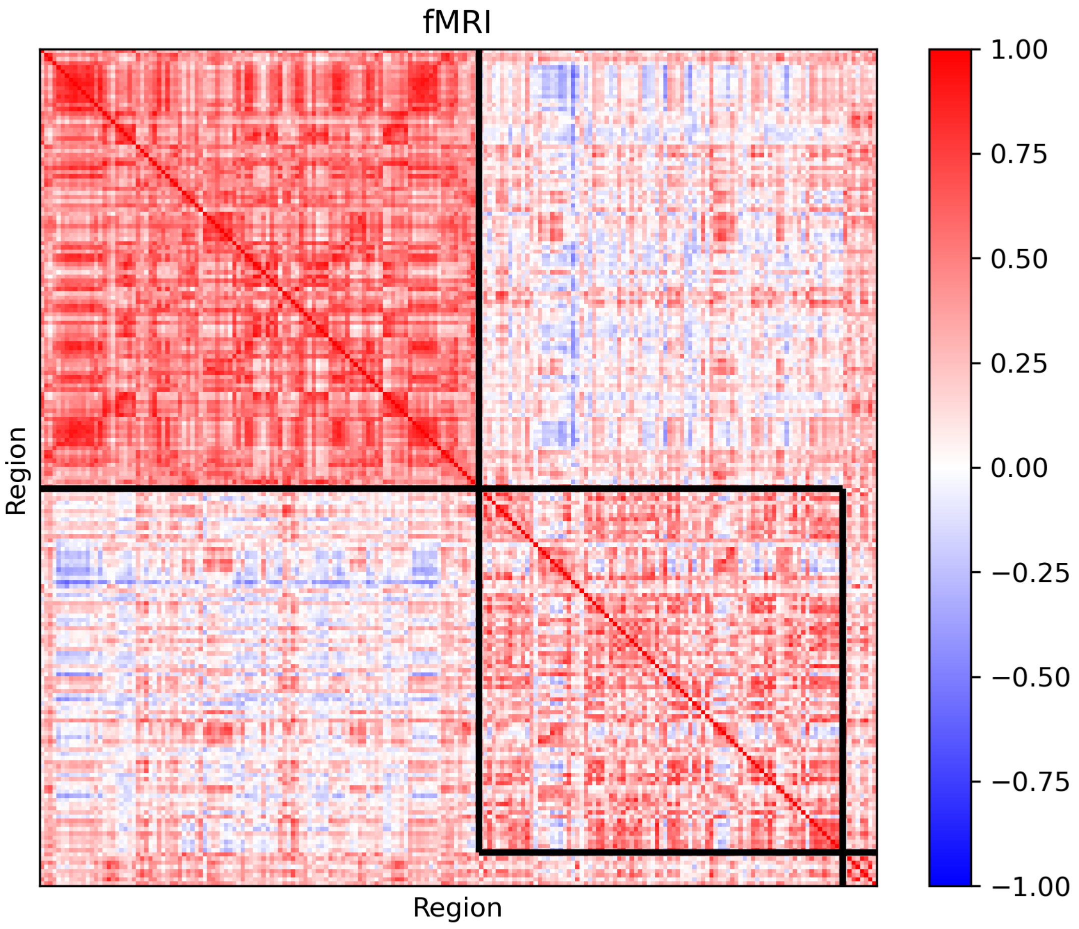
fMRI
Region
Region
1.00
0.75
0.50
0.25
0.00
−0.25
−0.50
−0.75
−1.00

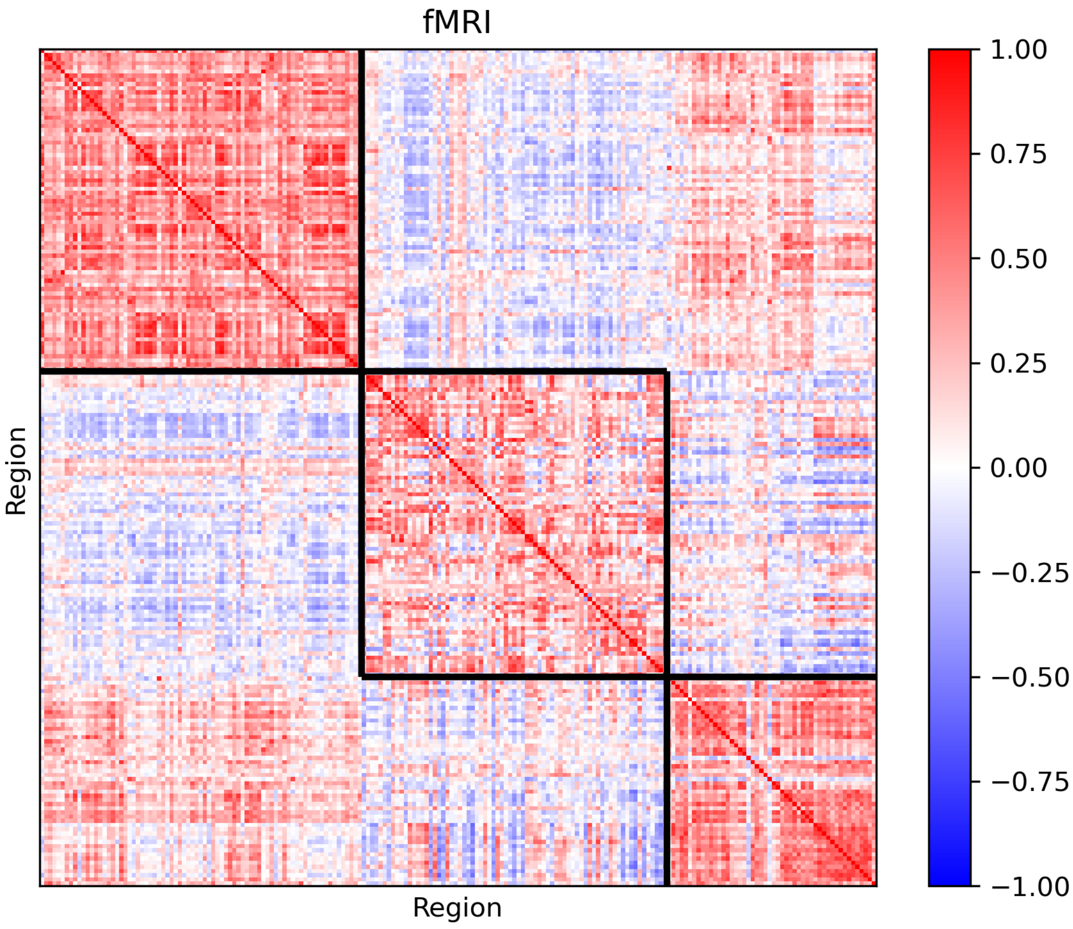
fMRI
Region
Region
1.00
0.75
0.50
0.25
0.00
−0.25
−0.50
−0.75
−1.00

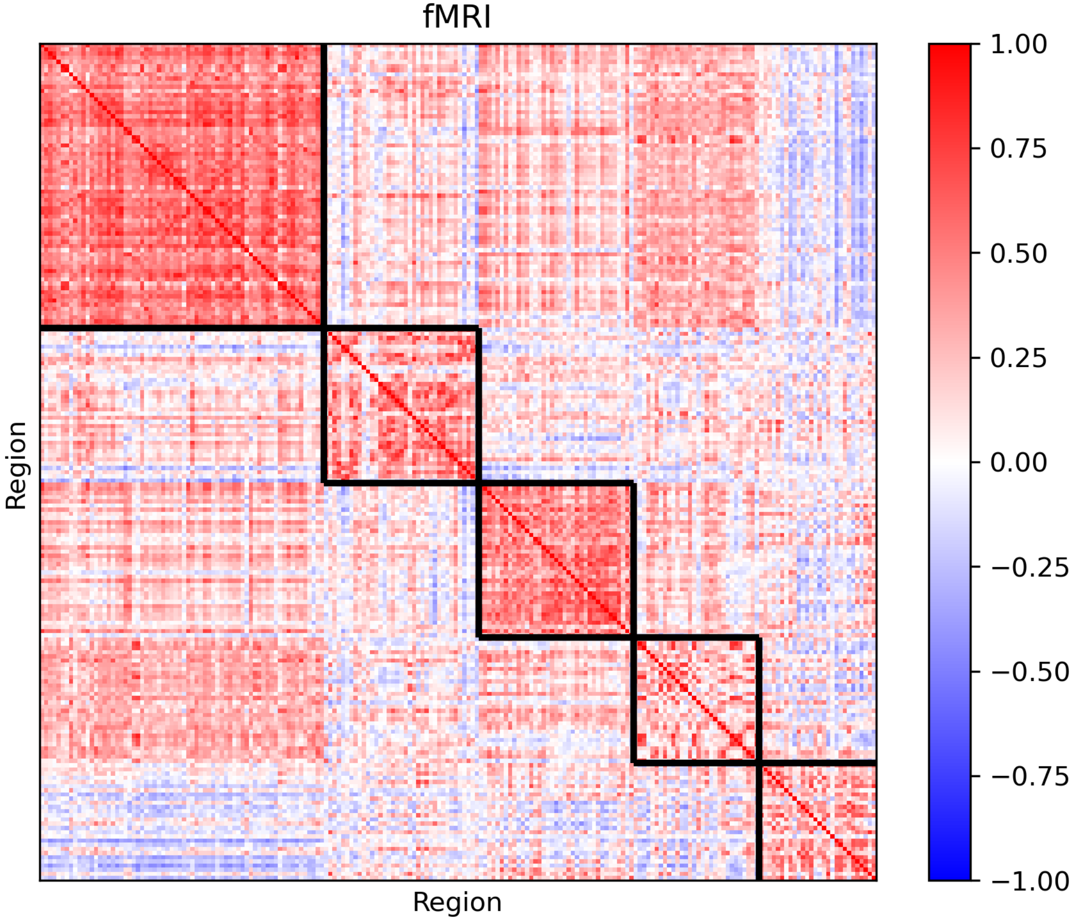
fMRI
Region
Region
1.00
0.75
0.50
0.25
0.00
−0.25
−0.50
−0.75
−1.00

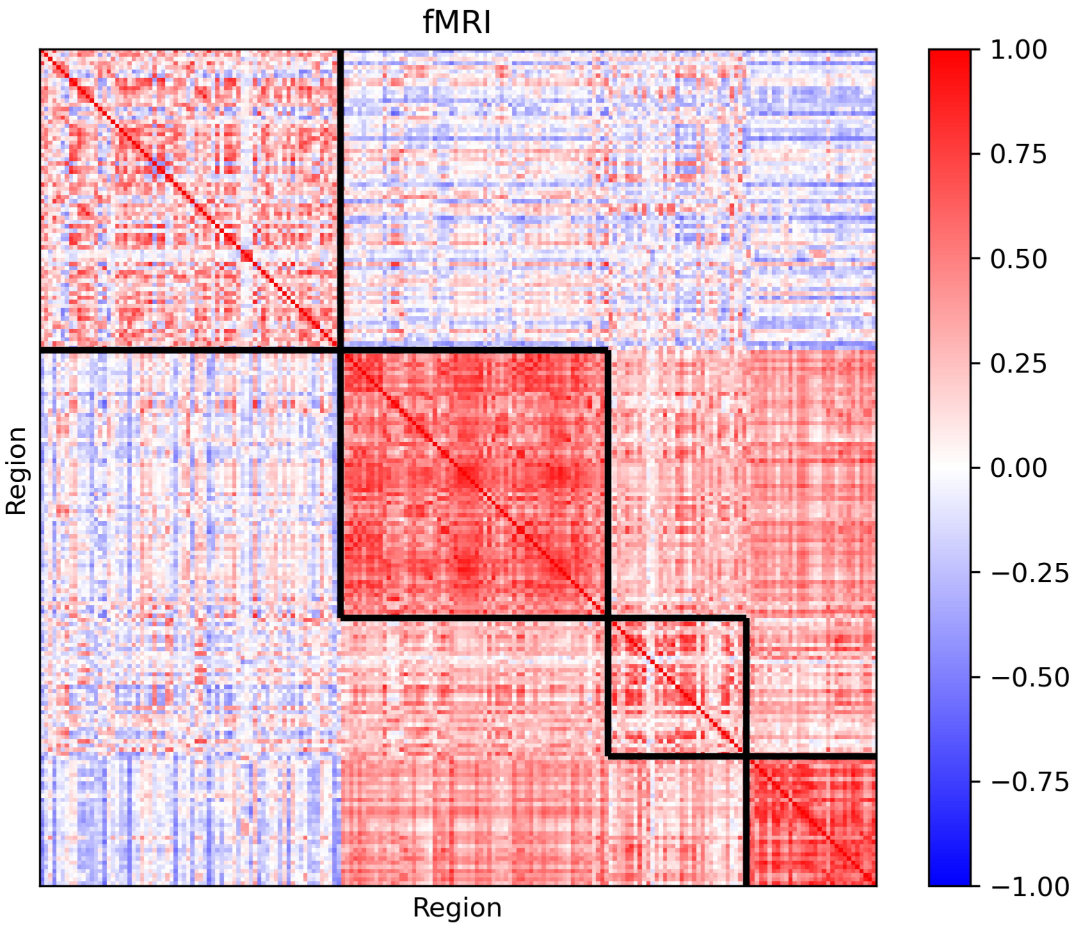
fMRI
Region
Region
1.00
0.75
0.50
0.25
0.00
−0.25
−0.50
−0.75
−1.00

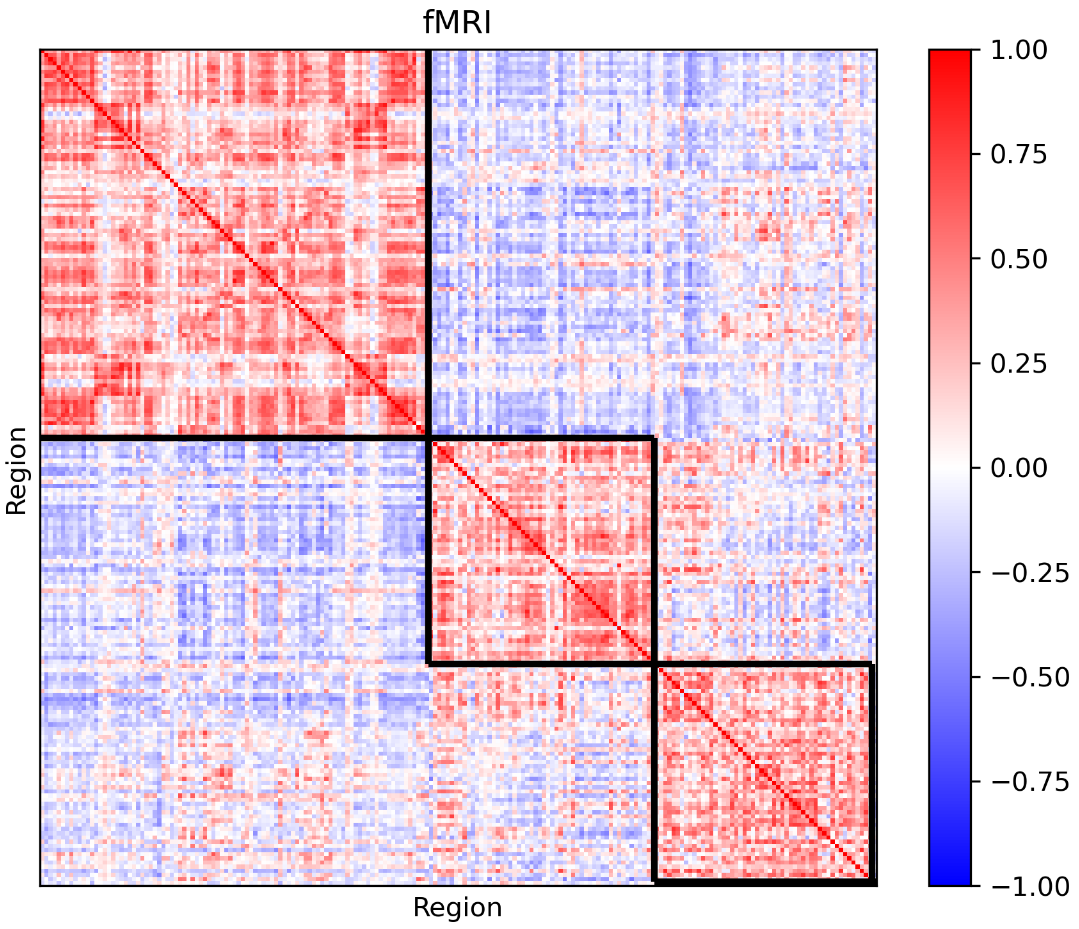
fMRI
Region
Region
1.00
0.75
0.50
0.25
0.00
−0.25
−0.50
−0.75
−1.00

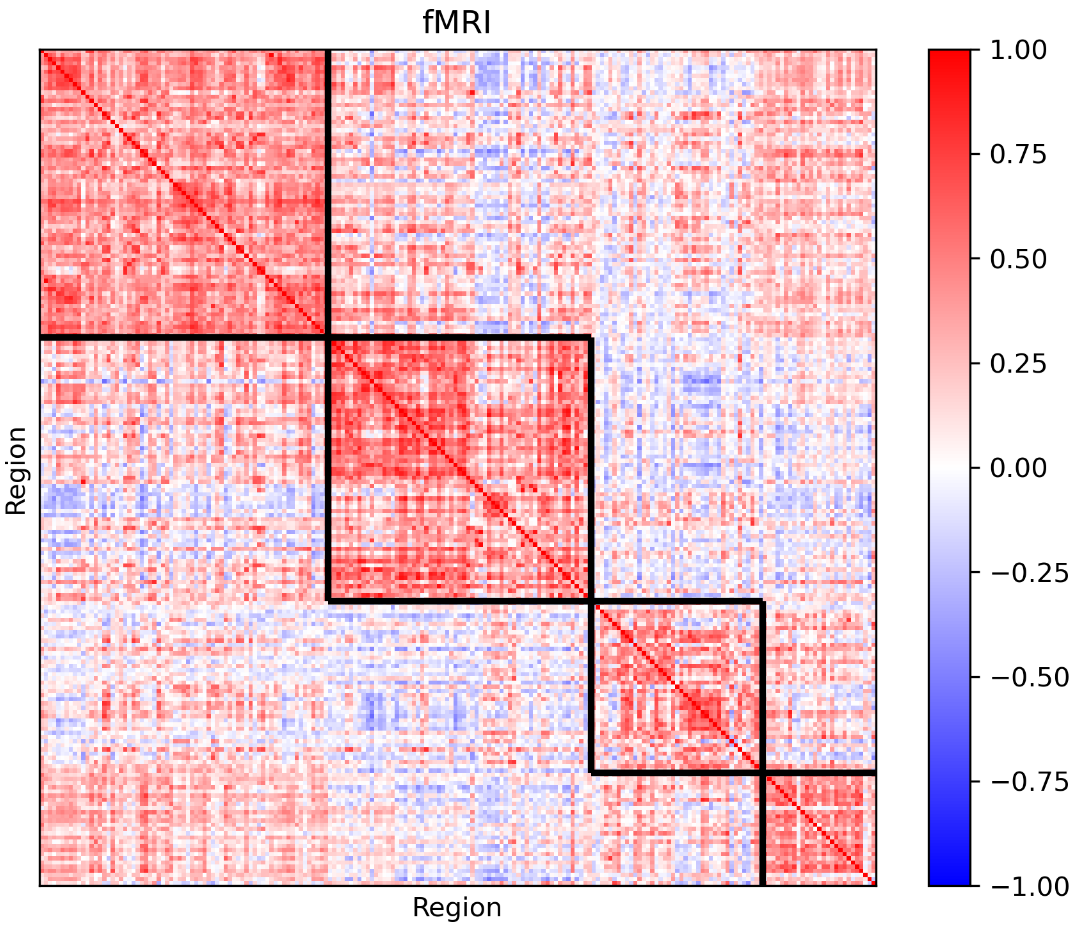
fMRI
Region
Region
1.00
0.75
0.50
0.25
0.00
−0.25
−0.50
−0.75
−1.00

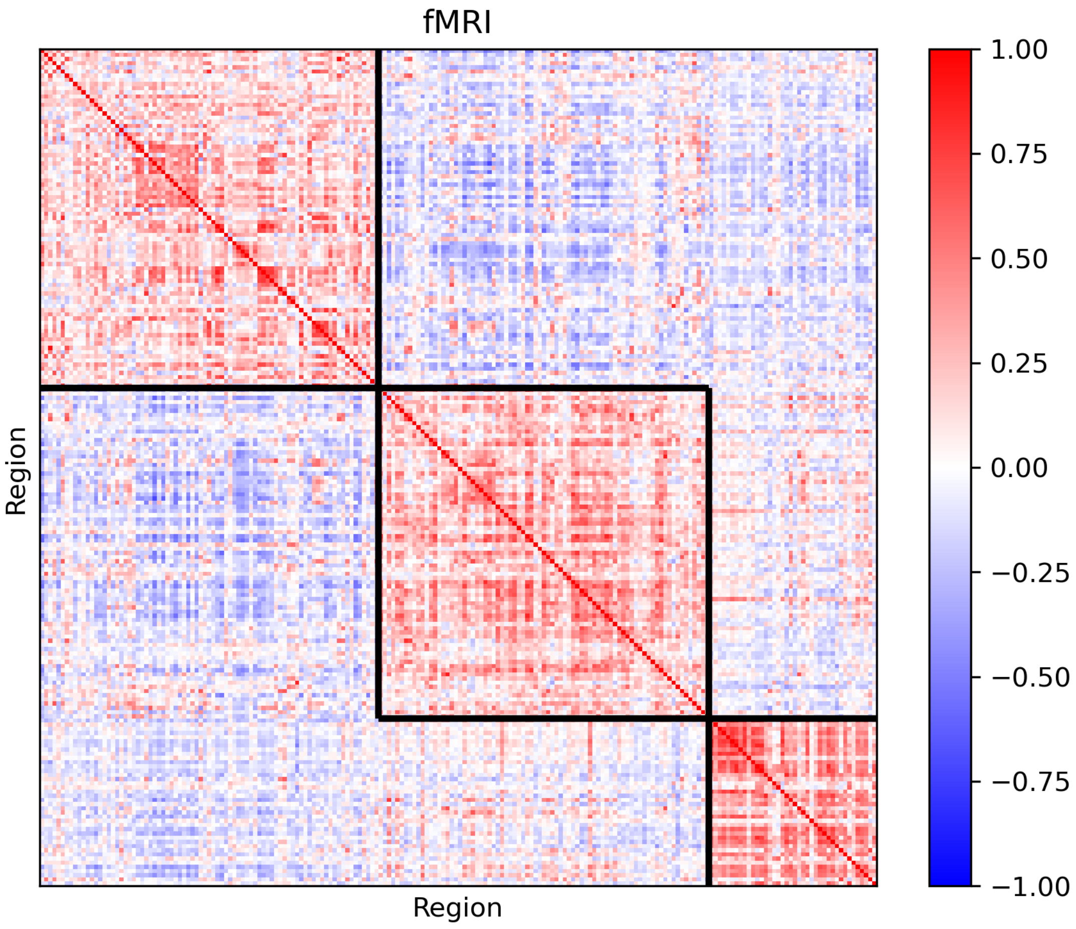
fMRI
Region
Region
1.00
0.75
0.50
0.25
0.00
−0.25
−0.50
−0.75
−1.00

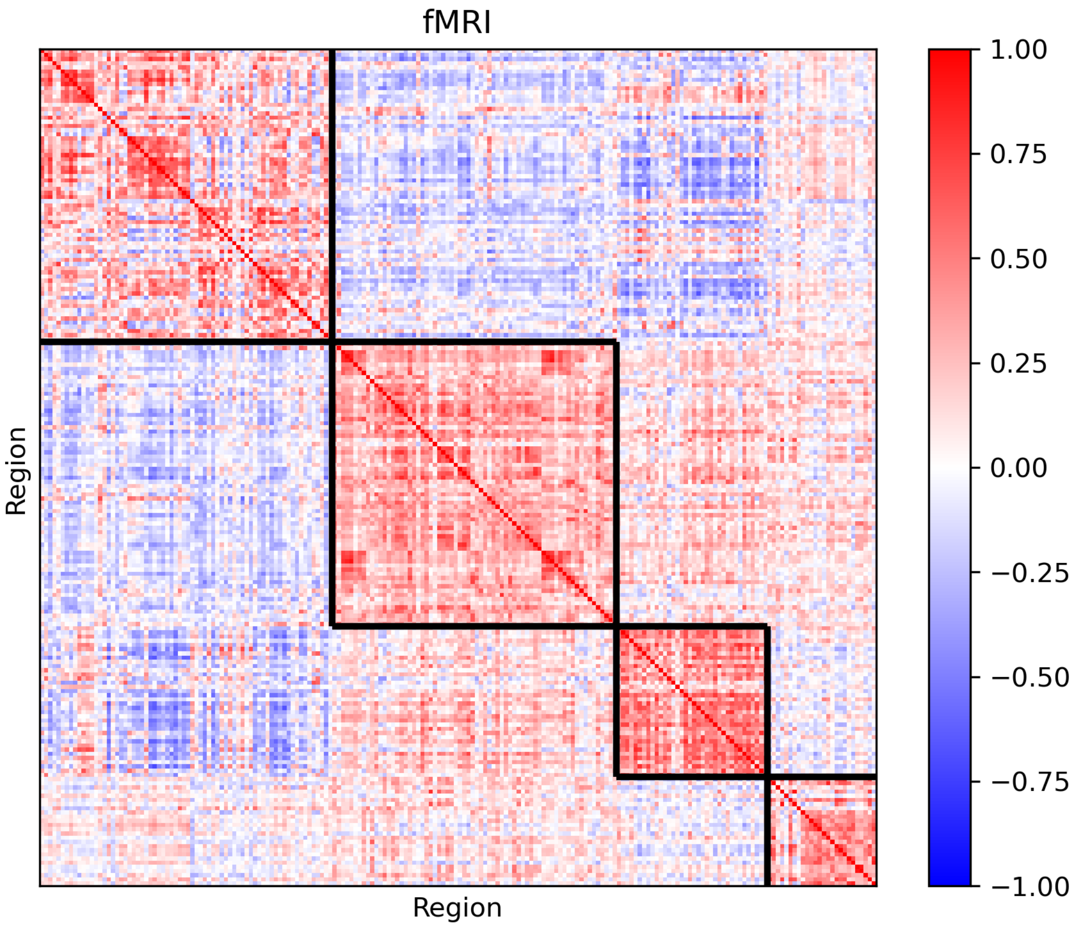
fMRI
Region
Region
1.00
0.75
0.50
0.25
0.00
−0.25
−0.50
−0.75
−1.00

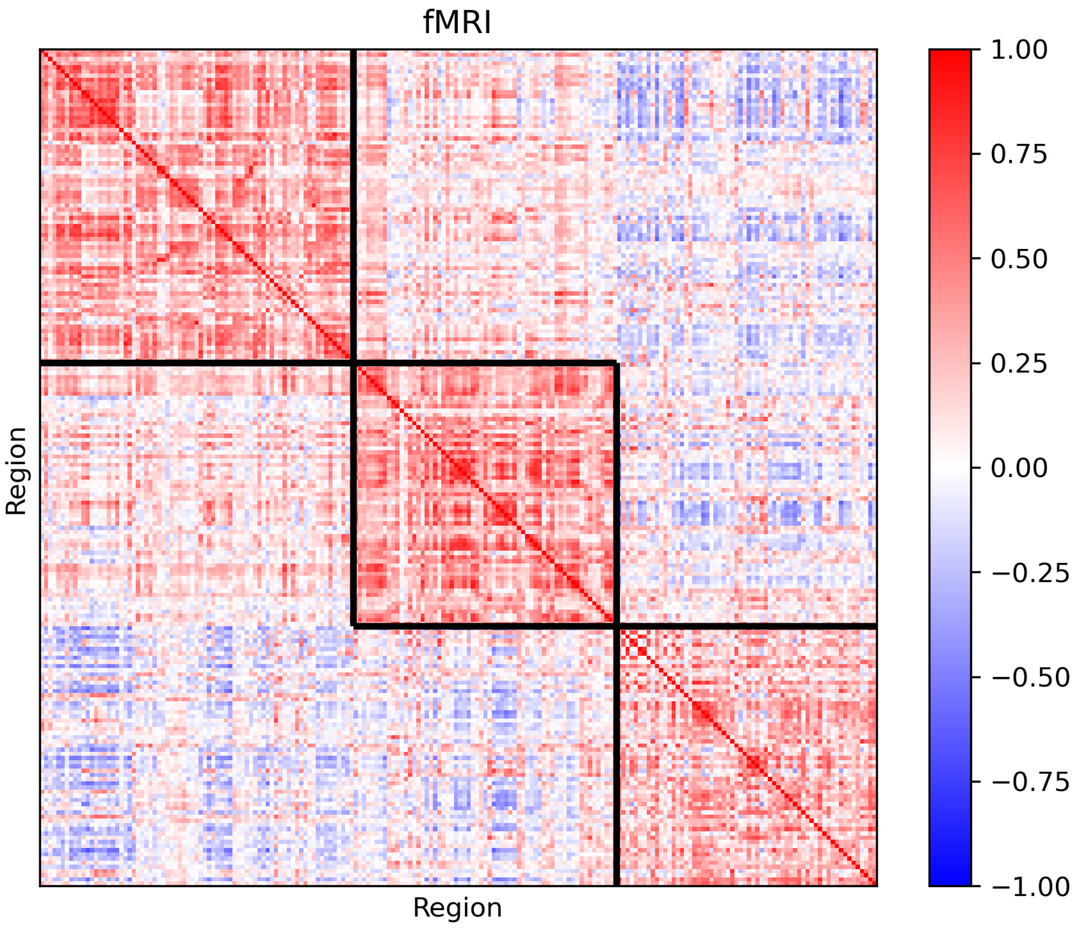
fMRI
Region
Region
1.00
0.75
0.50
0.25
0.00
−0.25
−0.50
−0.75
−1.00

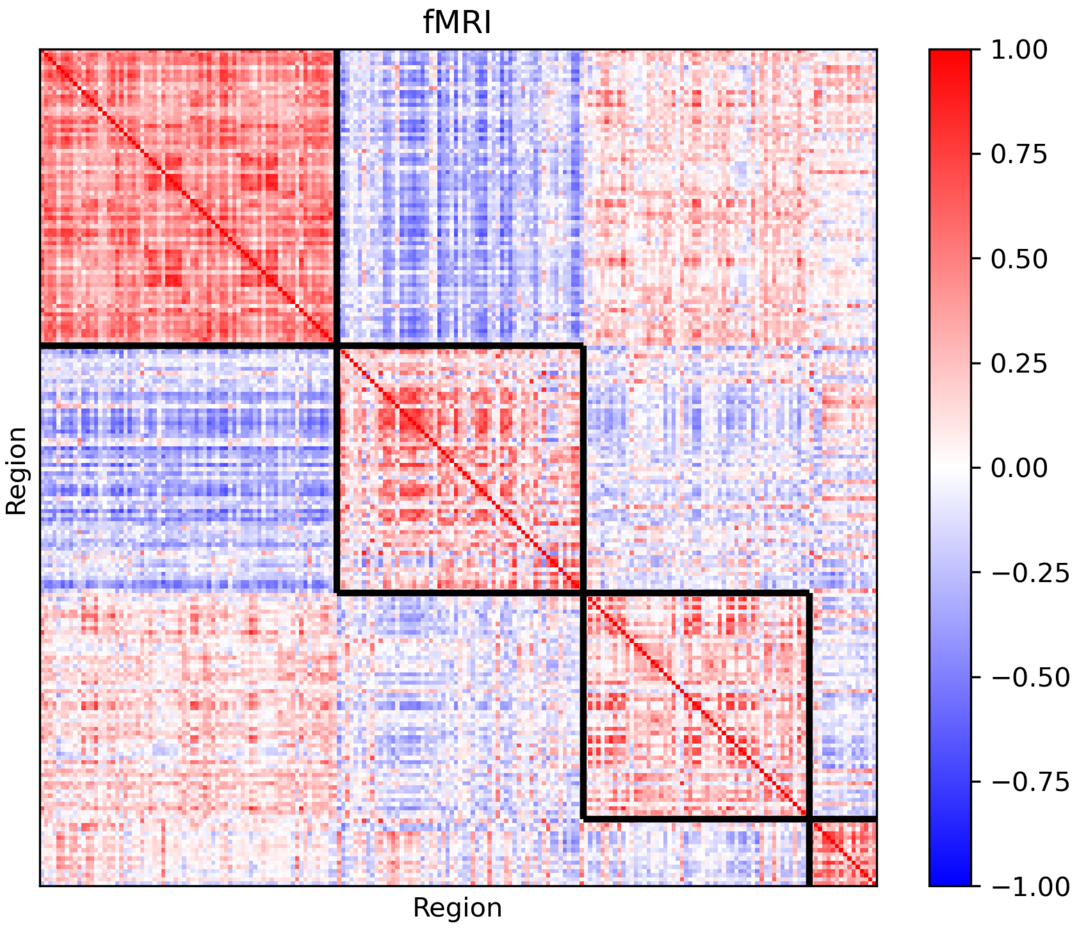
fMRI
Region
Region
1.00
0.75
0.50
0.25
0.00
−0.25
−0.50
−0.75
−1.00

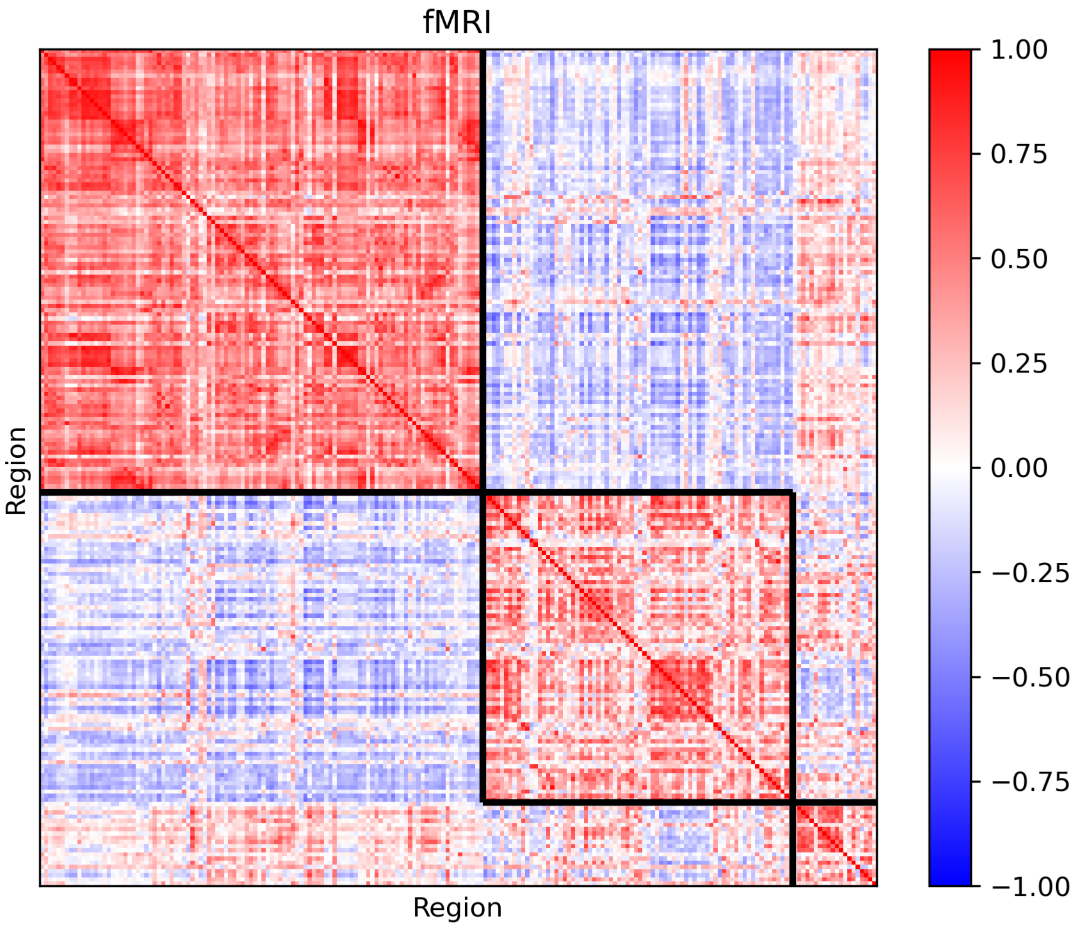
fMRI
Region
Region
1.00
0.75
0.50
0.25
0.00
−0.25
−0.50
−0.75
−1.00

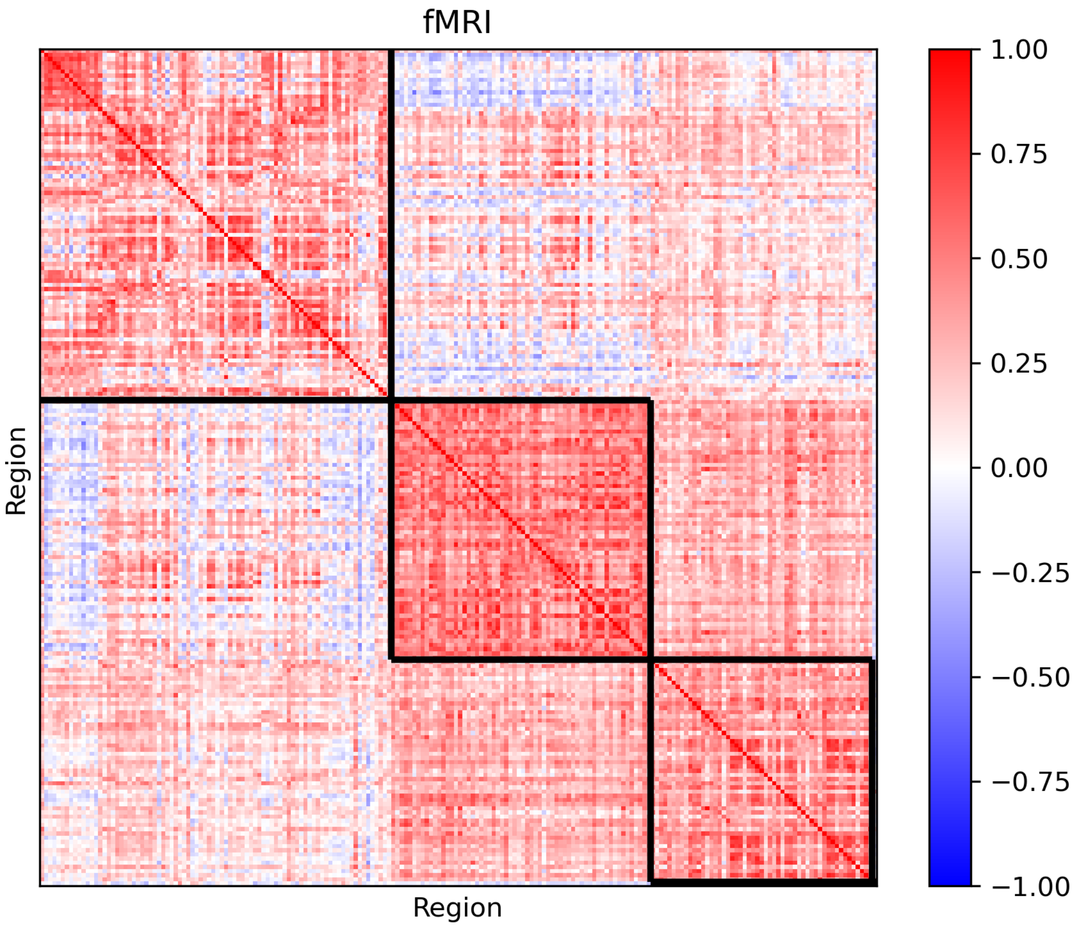

fMRI
Region
Region
1.00
0.75
0.50
0.25
0.00
−0.25
−0.50
−0.75
−1.00

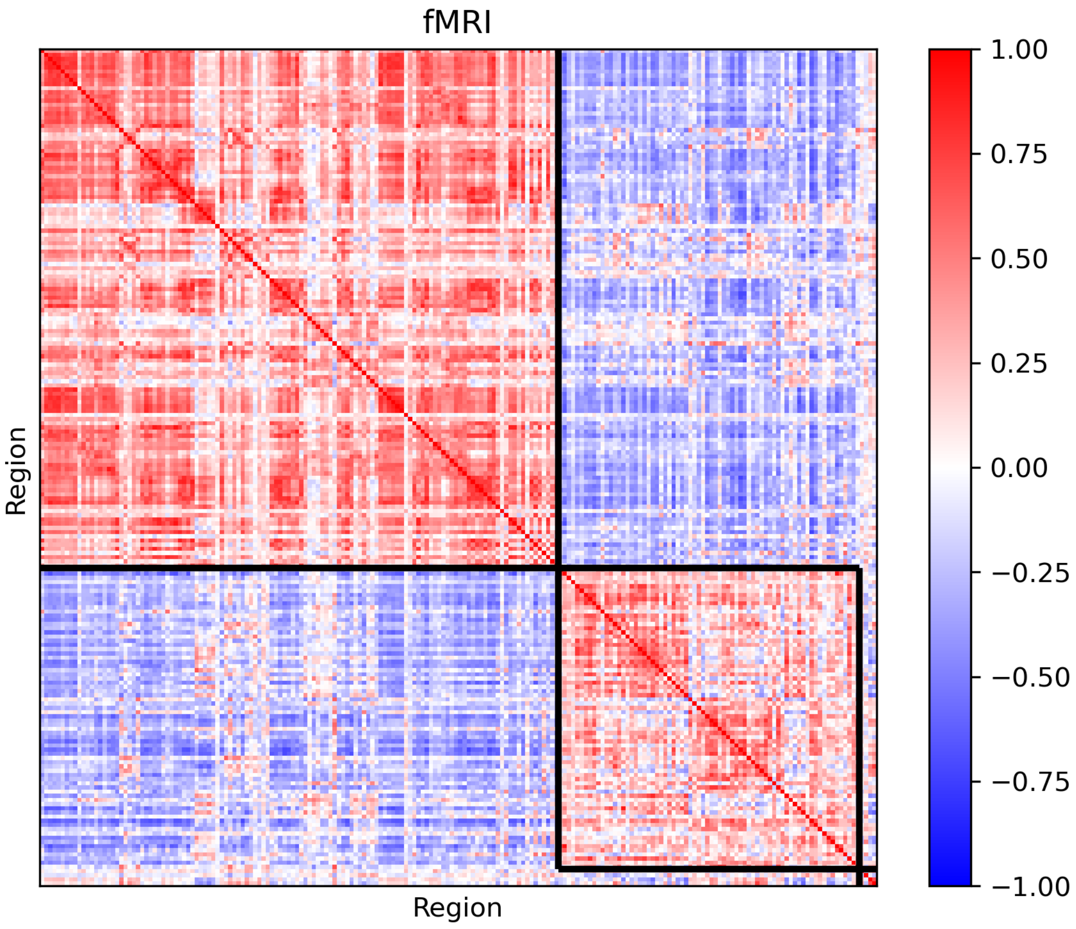
fMRI
Region
Region
1.00
0.75
0.50
0.25
0.00
−0.25
−0.50
−0.75
−1.00

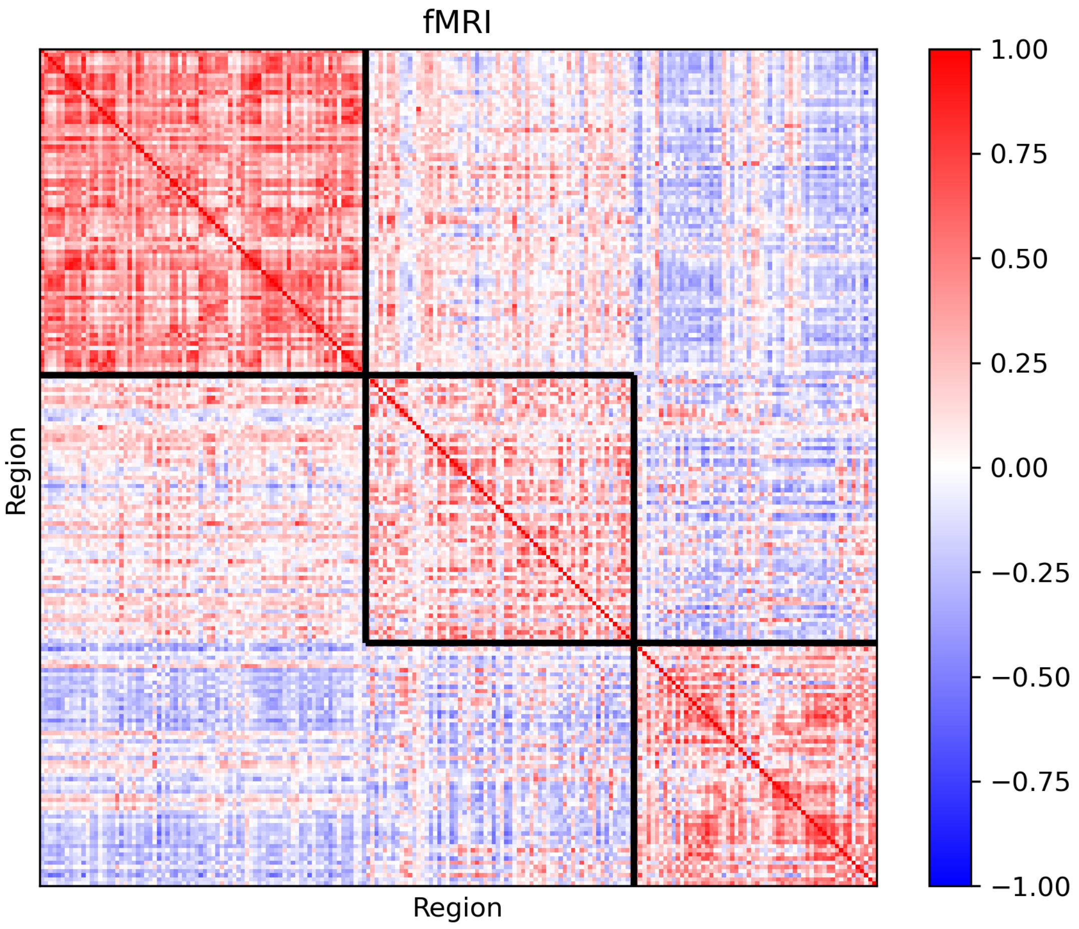
fMRI
Region
Region
1.00
0.75
0.50
0.25
0.00
−0.25
−0.50
−0.75
−1.00

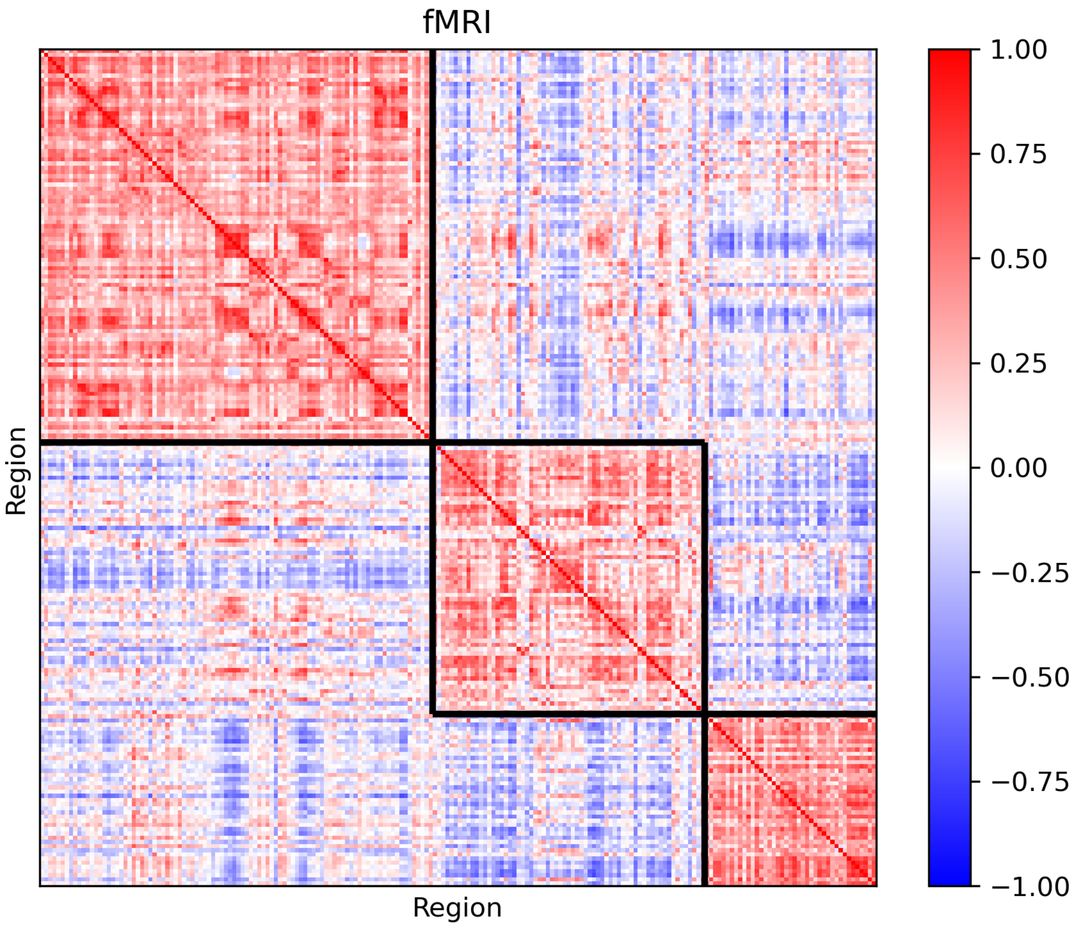
fMRI
Region
Region
1.00
0.75
0.50
0.25
0.00
−0.25
−0.50
−0.75
−1.00

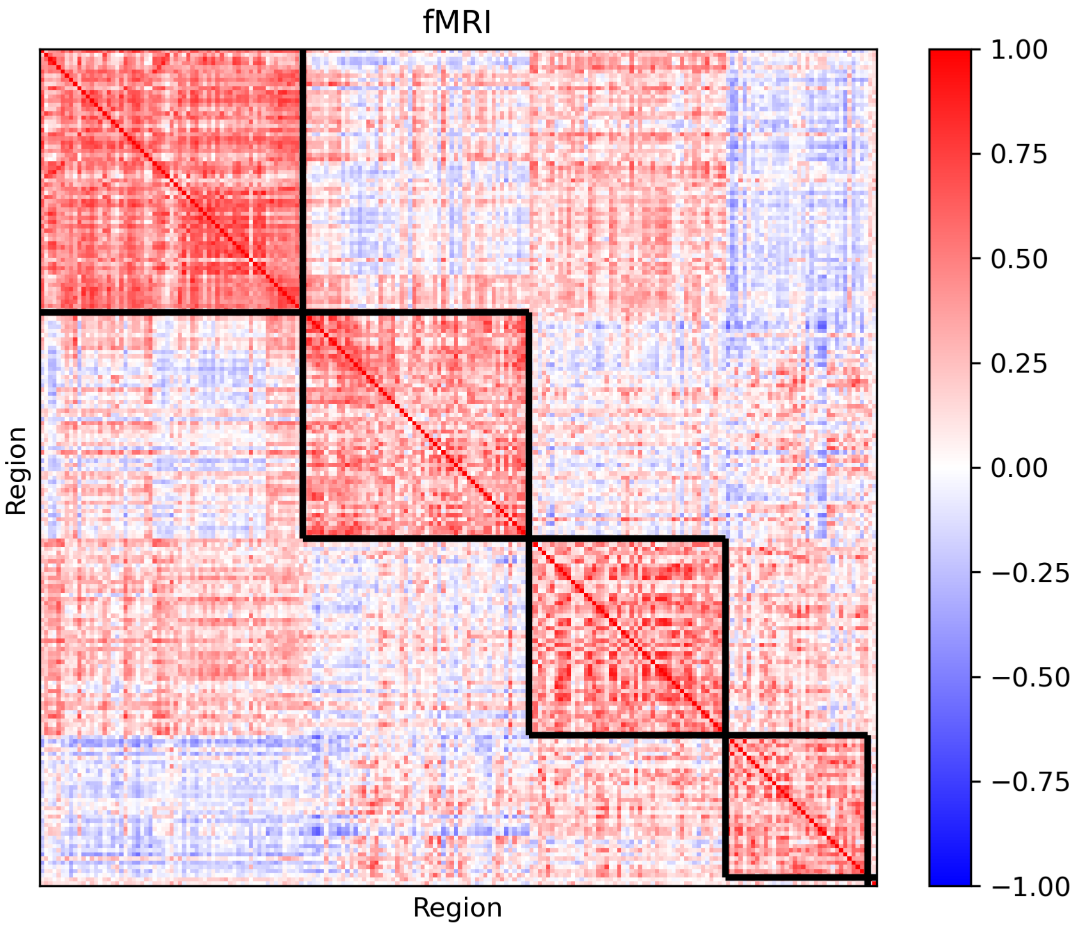
fMRI
Region
Region
1.00
0.75
0.50
0.25
0.00
−0.25
−0.50
−0.75
−1.00

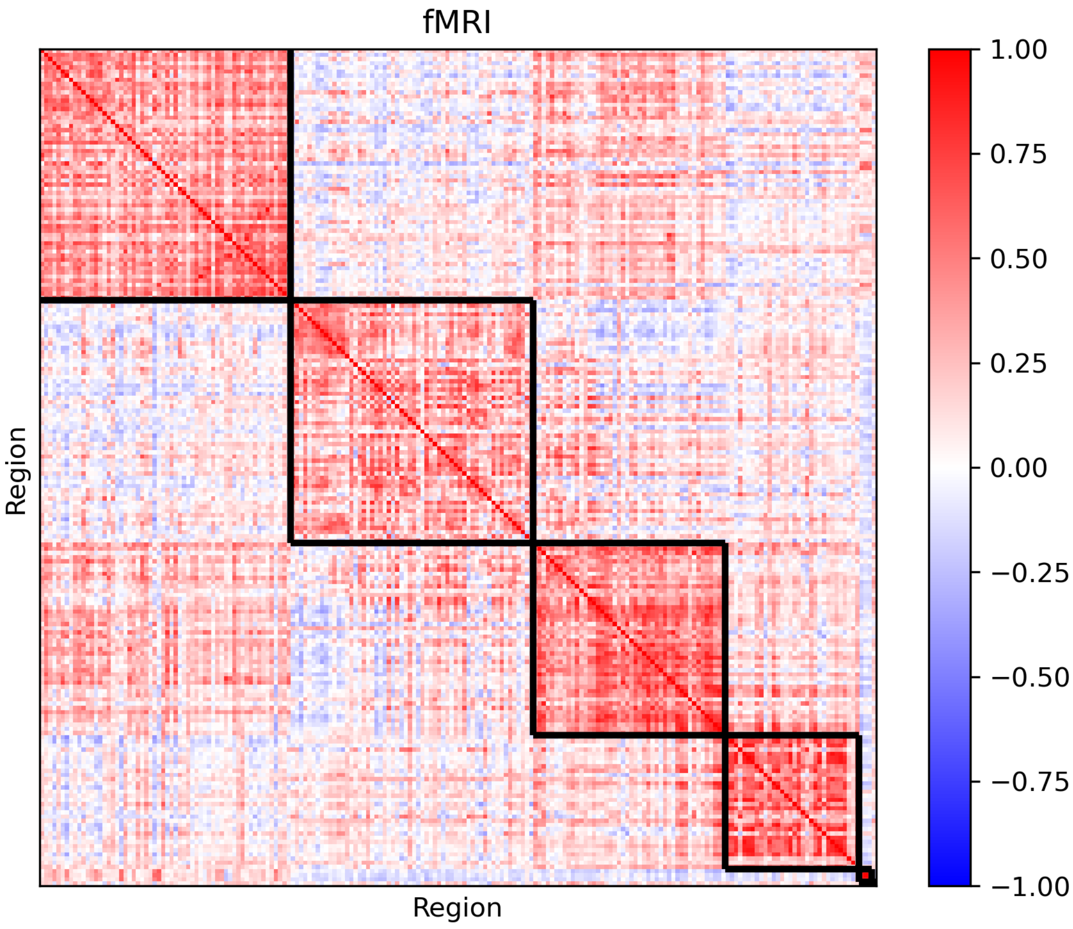
fMRI
Region
Region
1.00
0.75
0.50
0.25
0.00
−0.25
−0.50
−0.75
−1.00

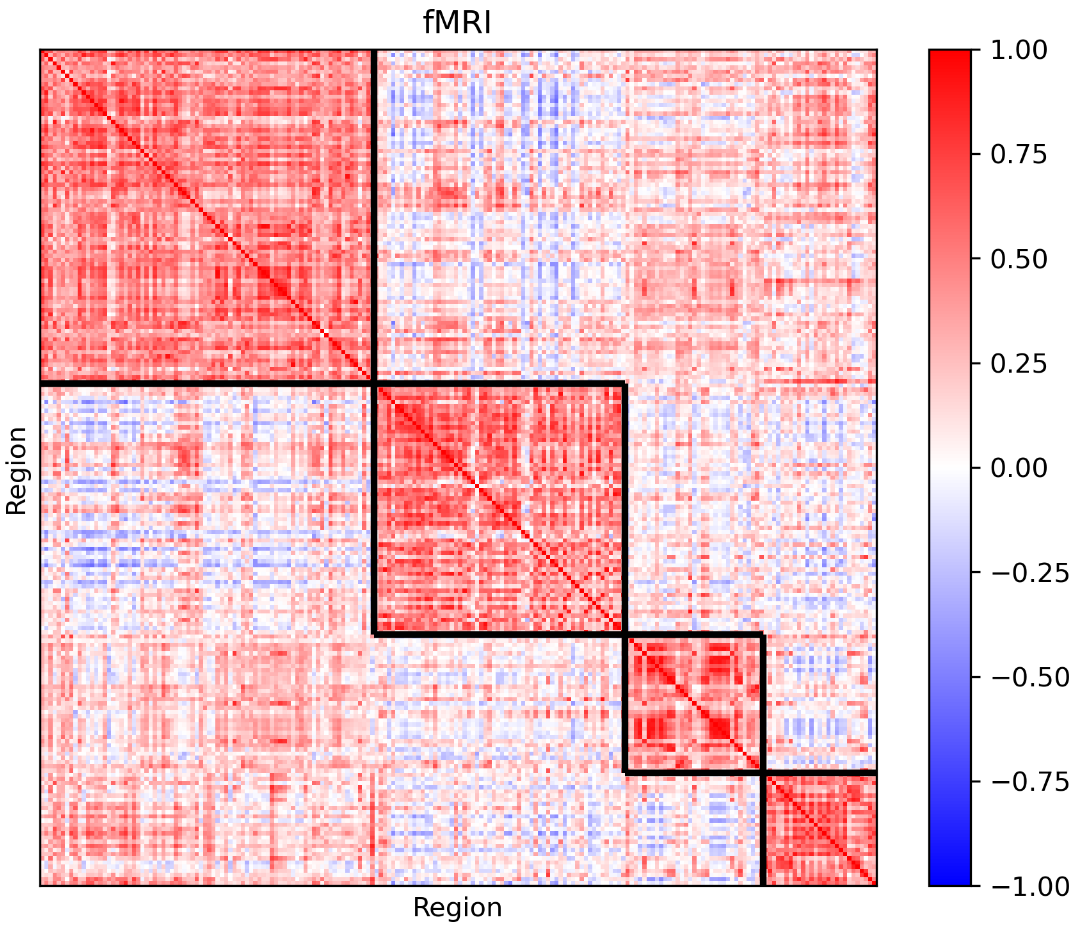
fMRI
Region
Region
1.00
0.75
0.50
0.25
0.00
−0.25
−0.50
−0.75
−1.00

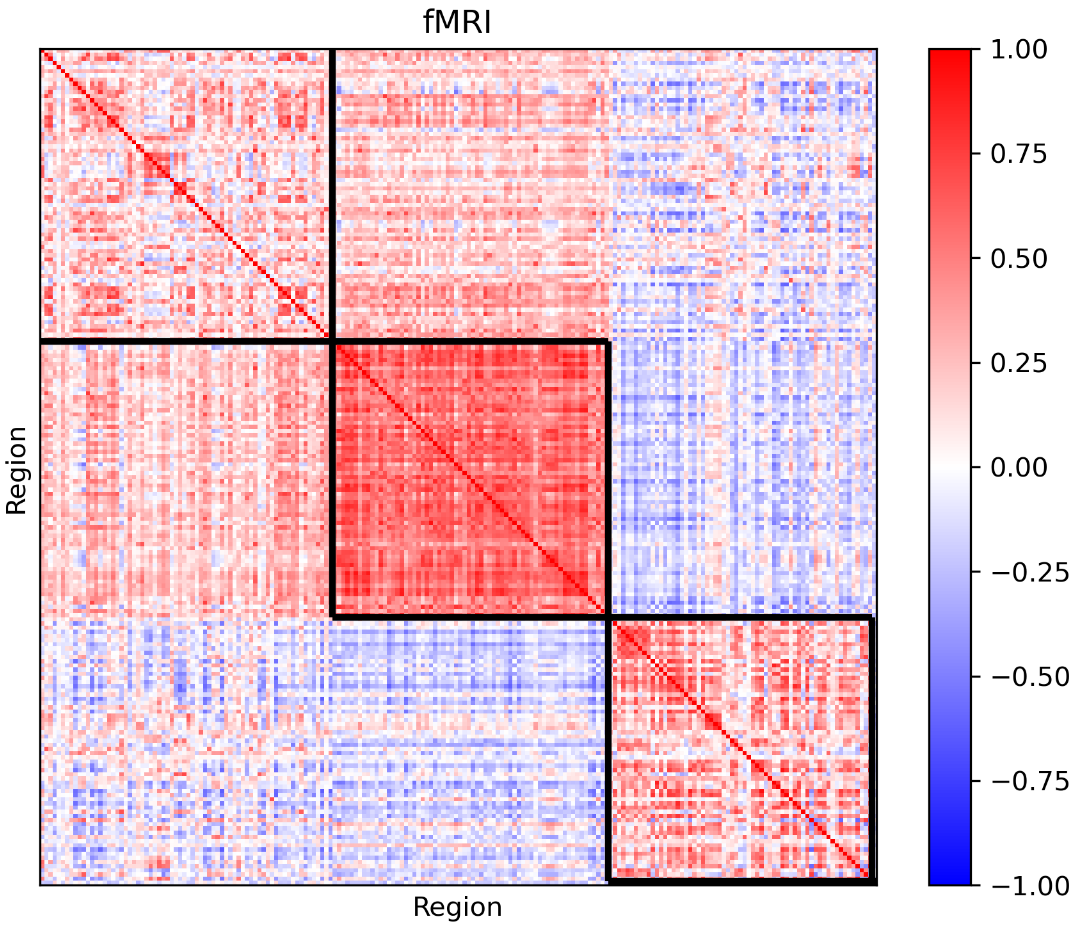
fMRI
Region
Region
1.00
0.75
0.50
0.25
0.00
−0.25
−0.50
−0.75
−1.00

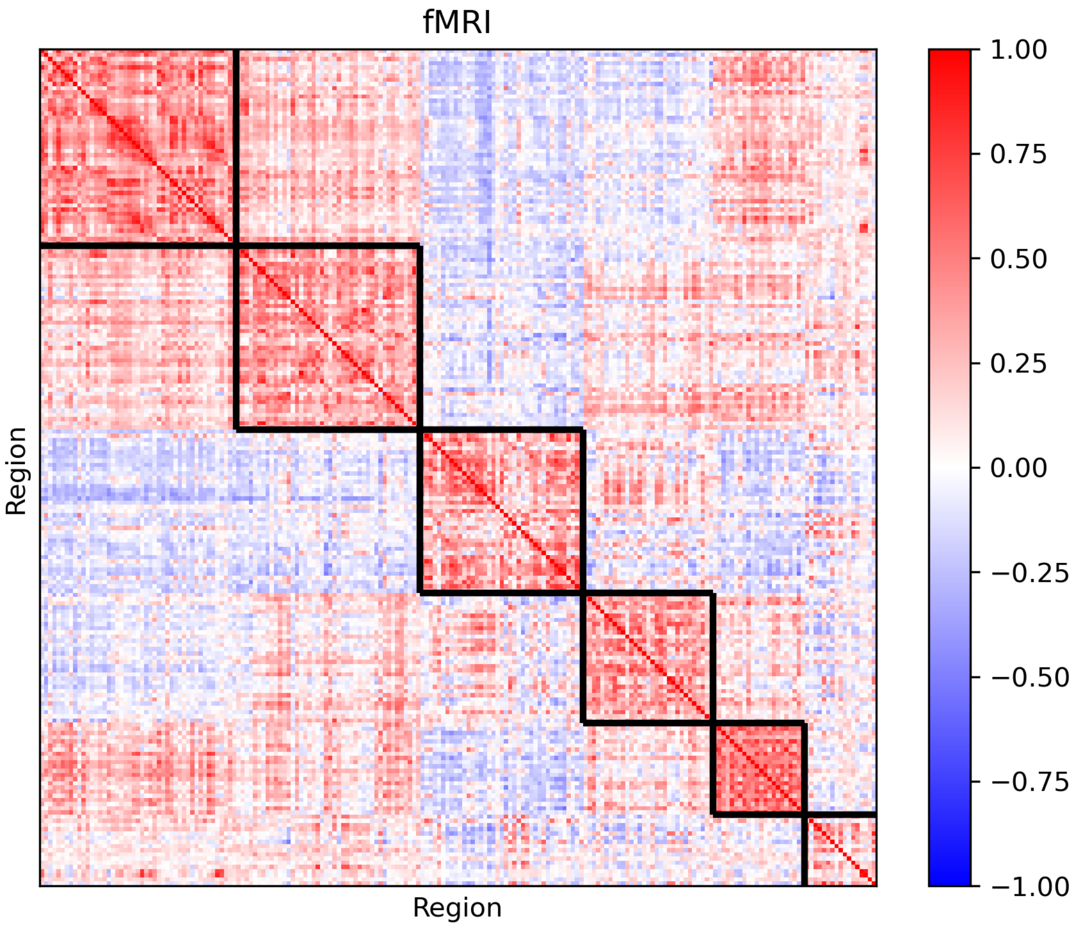
fMRI
Region
Region
1.00
0.75
0.50
0.25
0.00
−0.25
−0.50
−0.75
−1.00

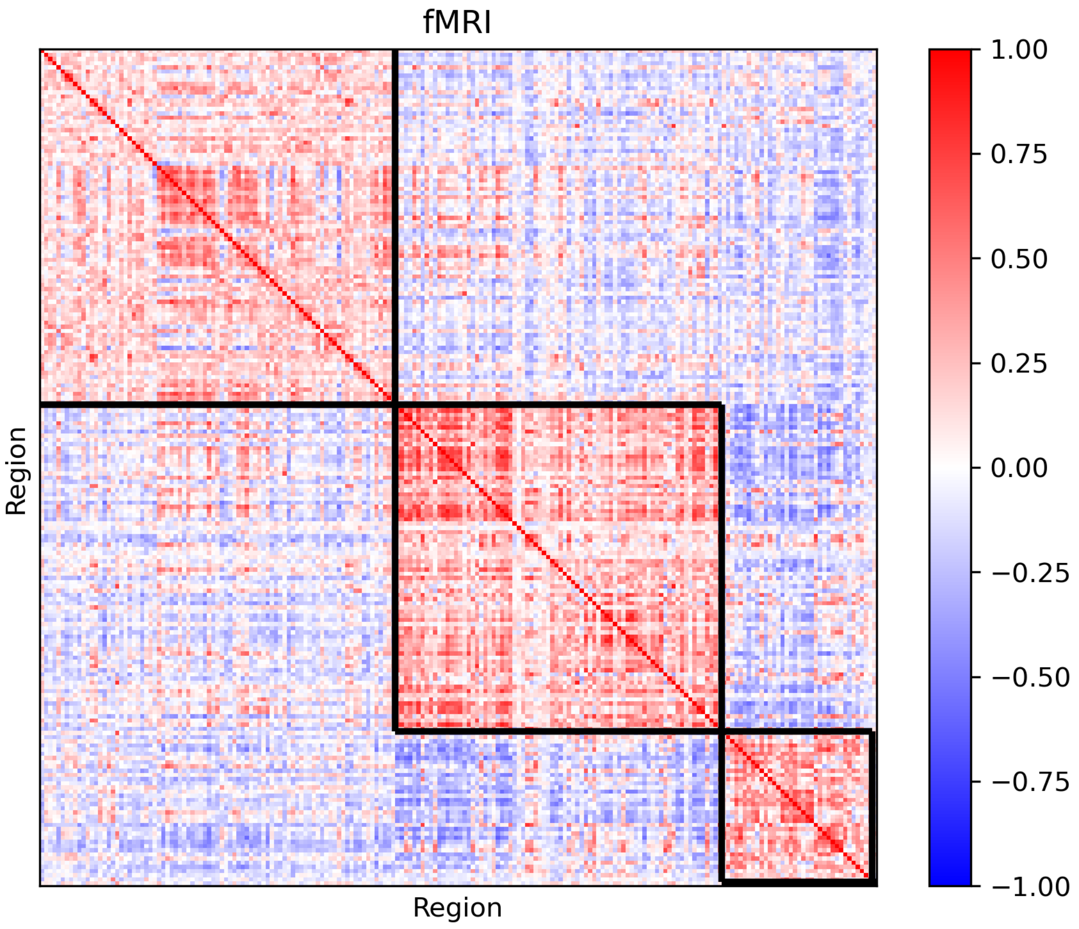
fMRI
Region
Region
1.00
0.75
0.50
0.25
0.00
−0.25
−0.50
−0.75
−1.00

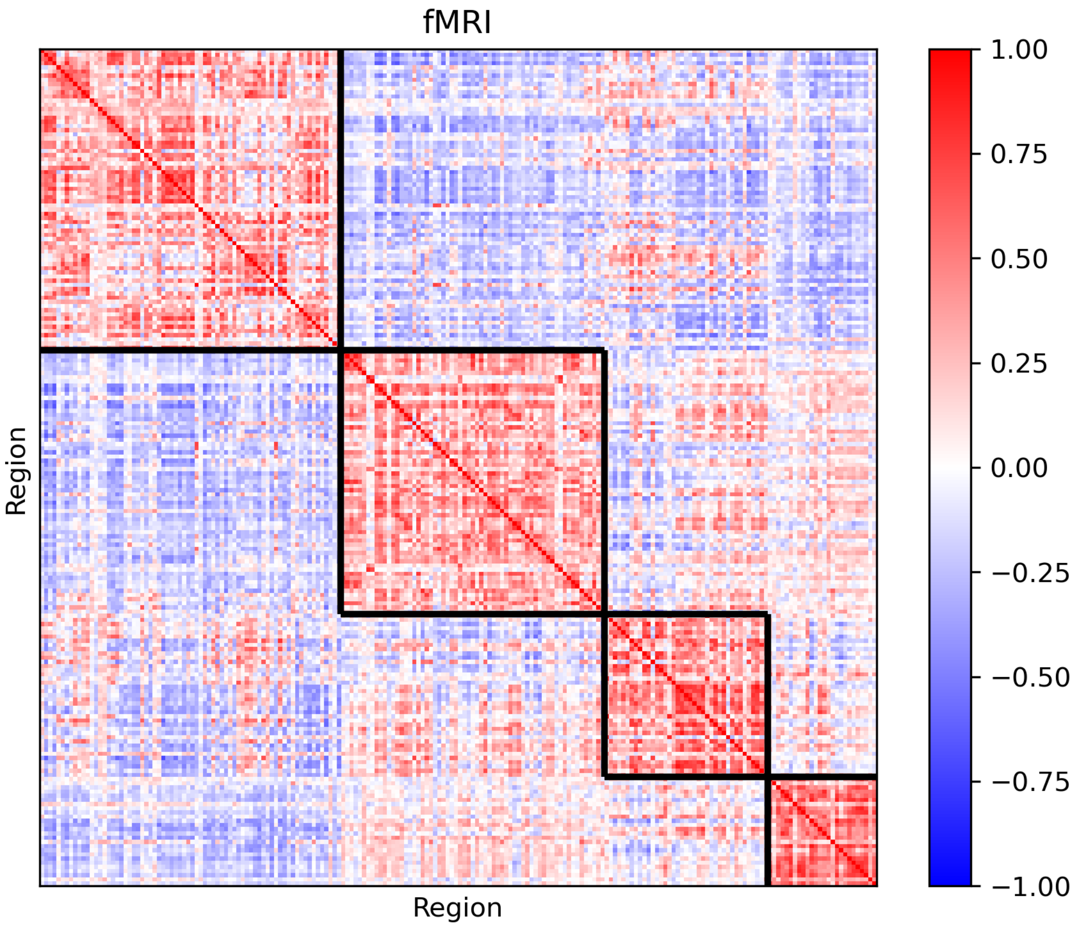
fMRI
Region
Region
1.00
0.75
0.50
0.25
0.00
−0.25
−0.50
−0.75
−1.00

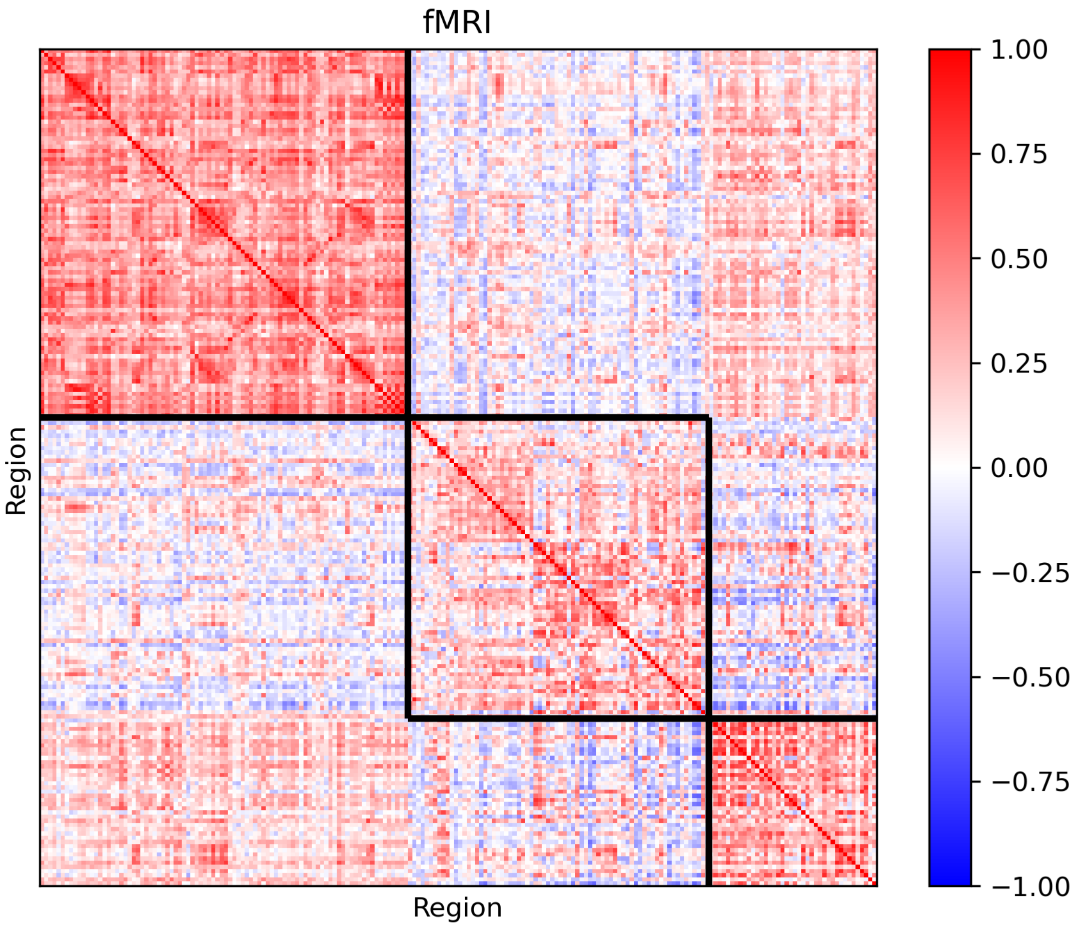
fMRI
Region
Region
1.00
0.75
0.50
0.25
0.00
−0.25
−0.50
−0.75
−1.00

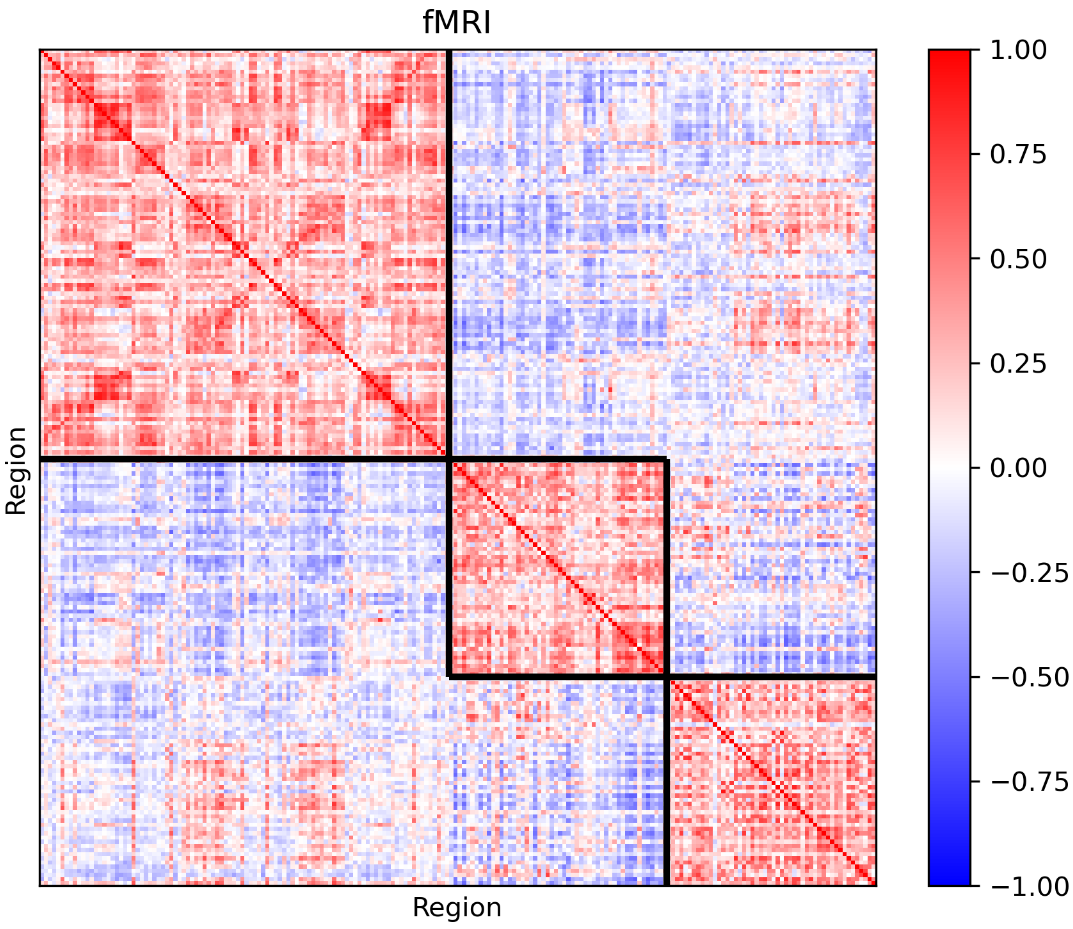
fMRI
Region
Region
1.00
0.75
0.50
0.25
0.00
−0.25
−0.50
−0.75
−1.00

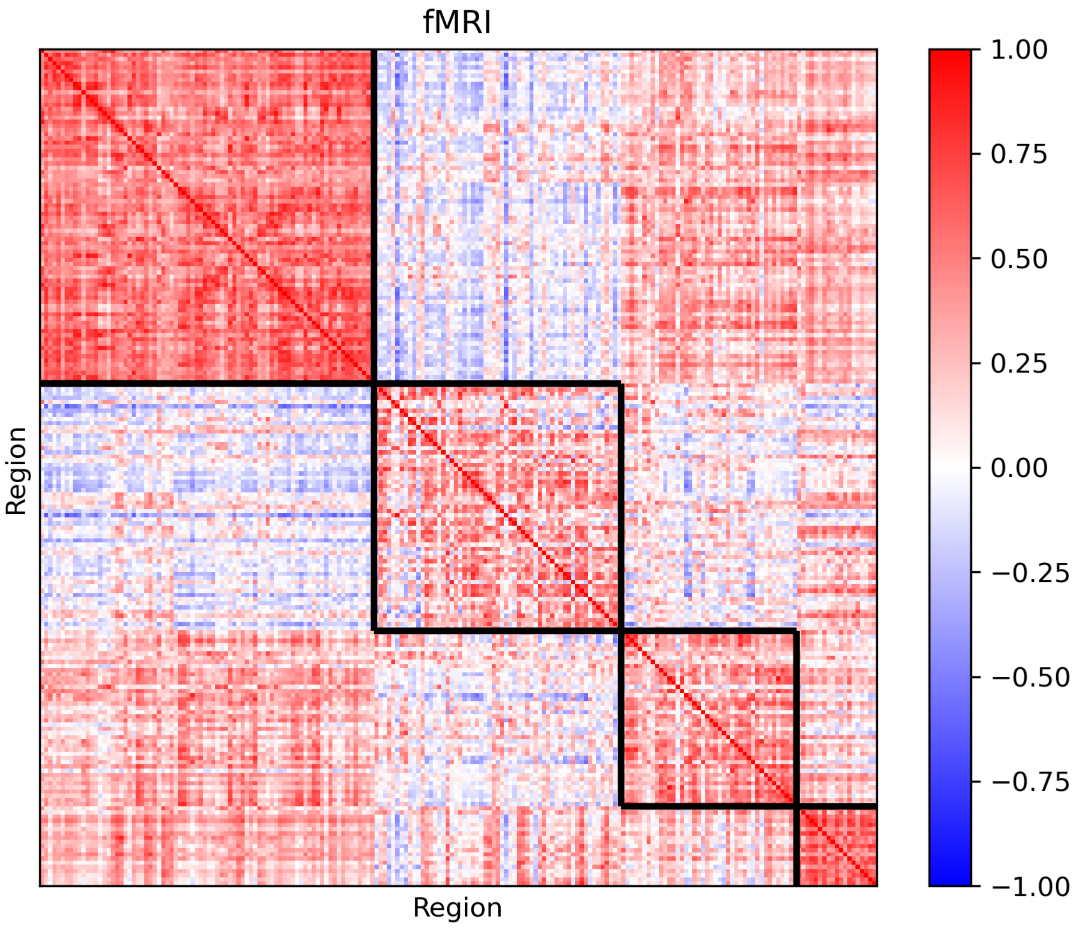

fMRI
Region
Region
1.00
0.75
0.50
0.25
0.00
−0.25
−0.50
−0.75
−1.00

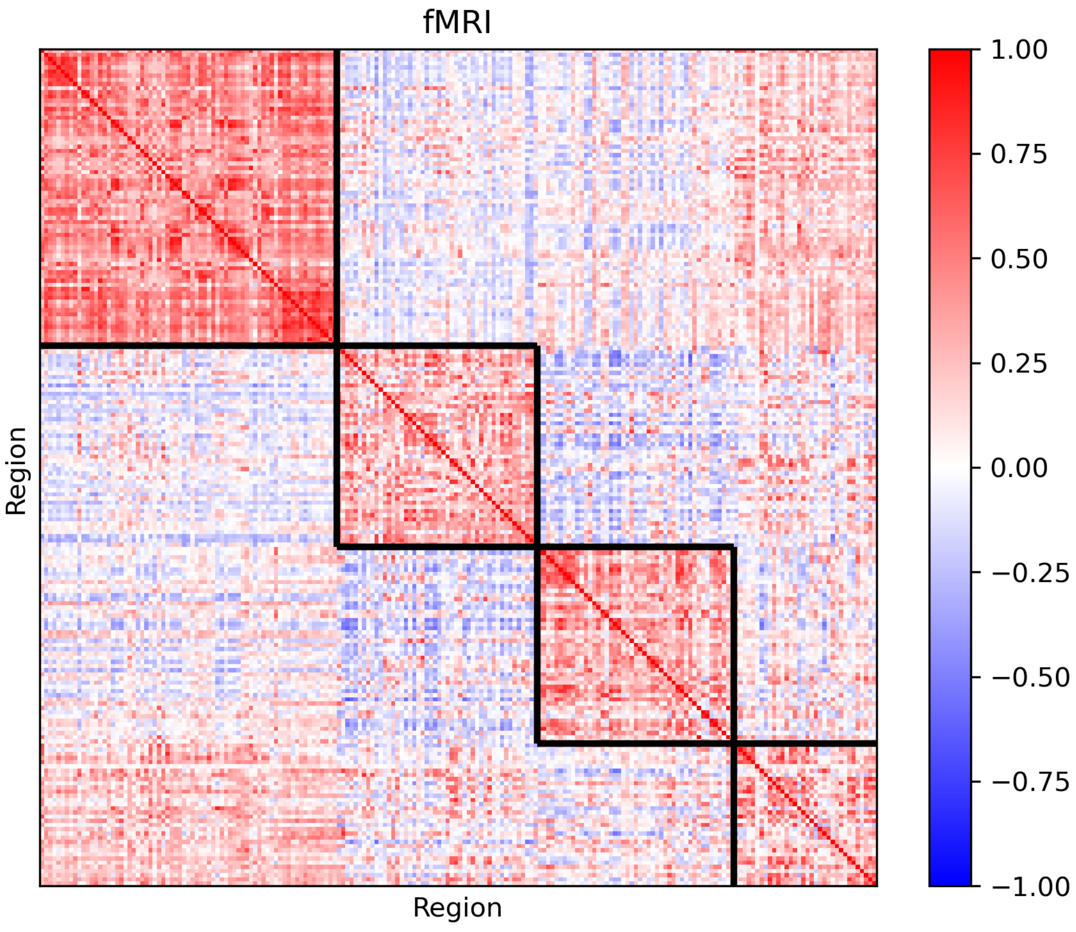
fMRI
Region
Region
1.00
0.75
0.50
0.25
0.00
−0.25
−0.50
−0.75
−1.00

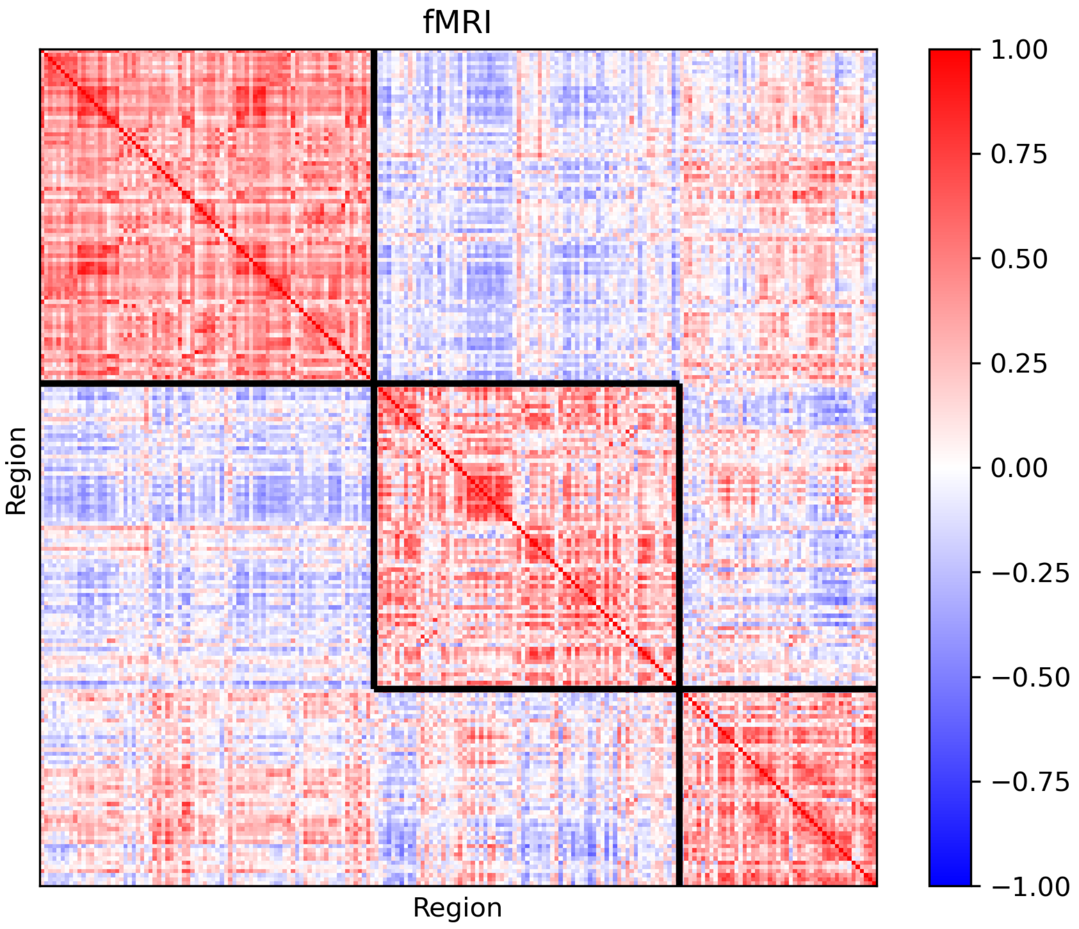
fMRI
Region
Region
1.00
0.75
0.50
0.25
0.00
−0.25
−0.50
−0.75
−1.00

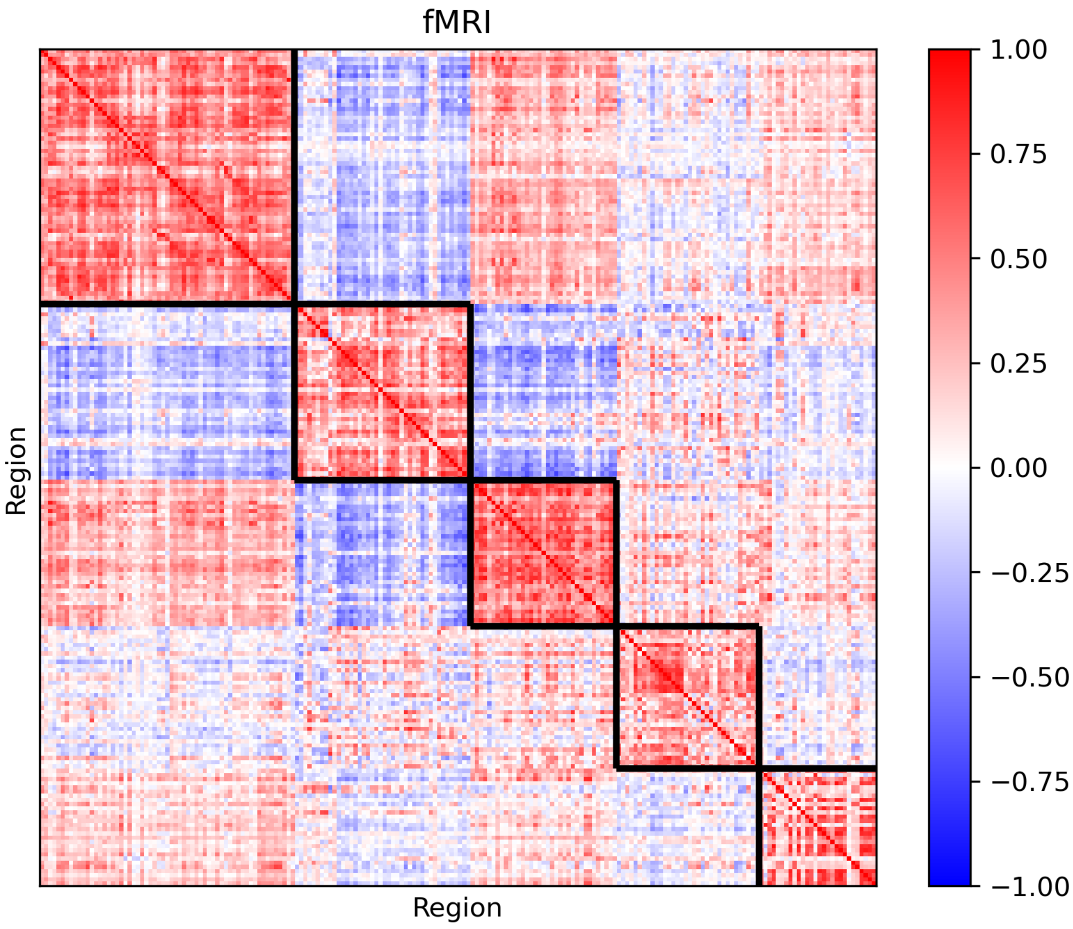
fMRI
Region
Region
1.00
0.75
0.50
0.25
0.00
−0.25
−0.50
−0.75
−1.00

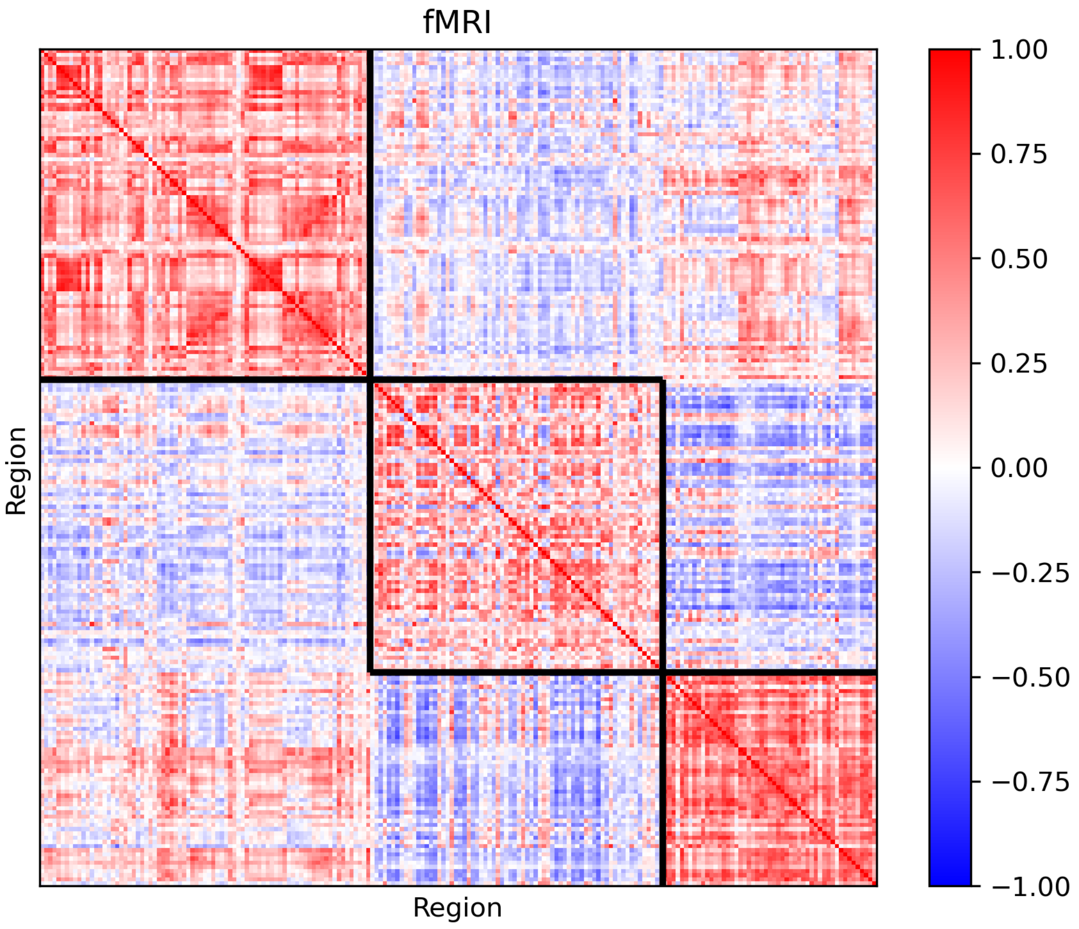
fMRI
Region
Region
1.00
0.75
0.50
0.25
0.00
−0.25
−0.50
−0.75
−1.00

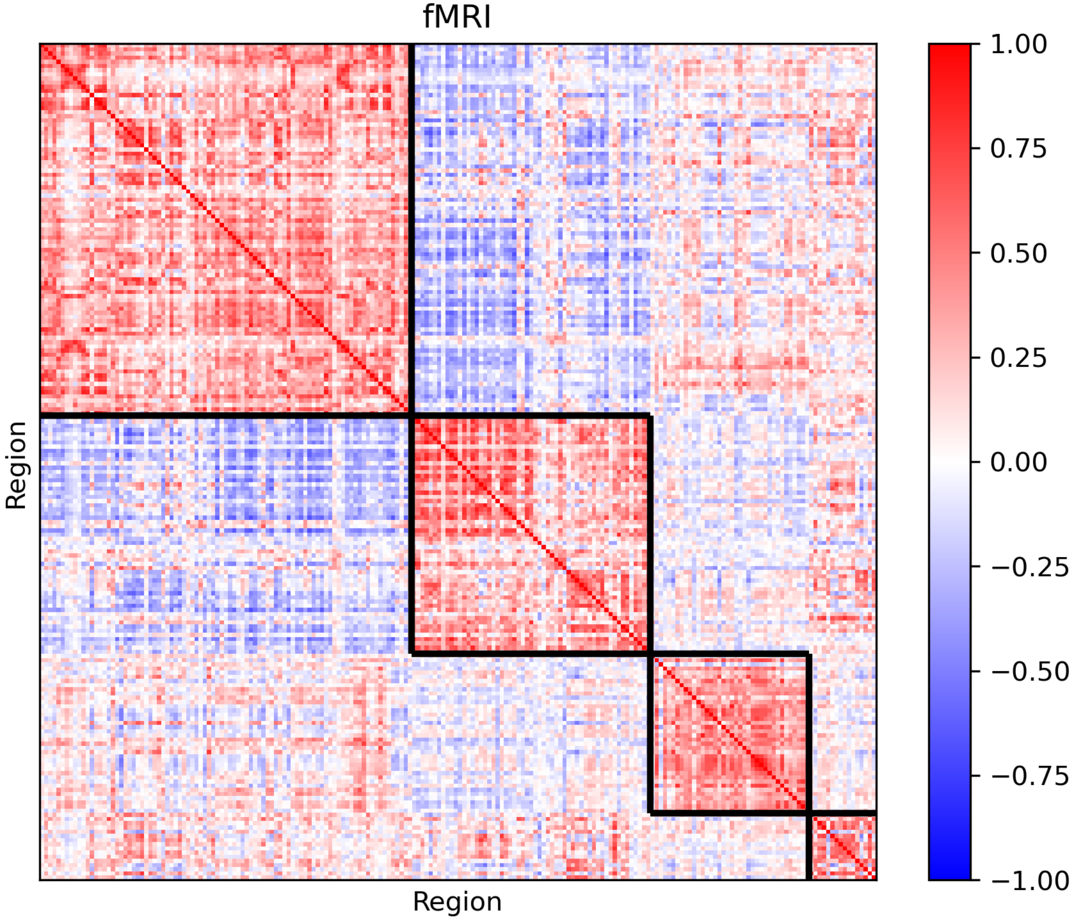
fMRI
Region
Region
1.00
0.75
0.50
0.25
0.00
−0.25
−0.50
−0.75
−1.00

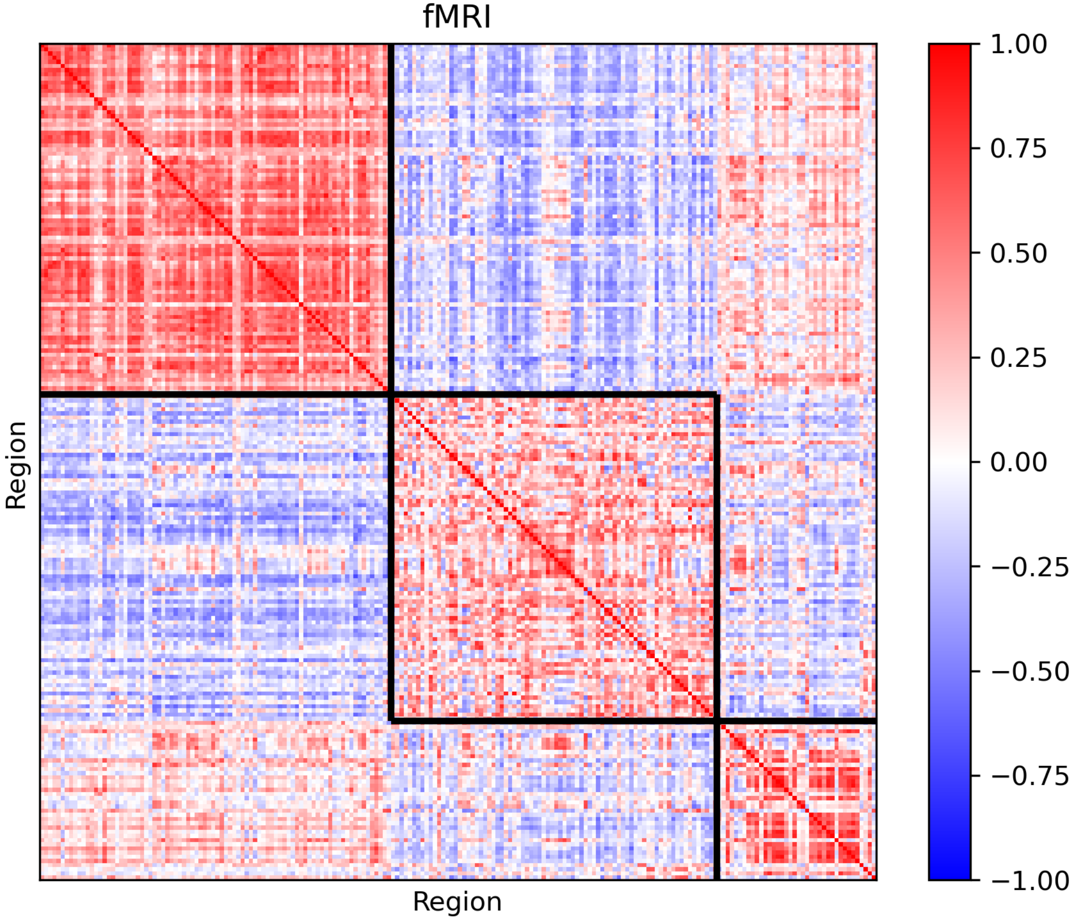
fMRI
Region
Region
1.00
0.75
0.50
0.25
0.00
−0.25
−0.50
−0.75
−1.00

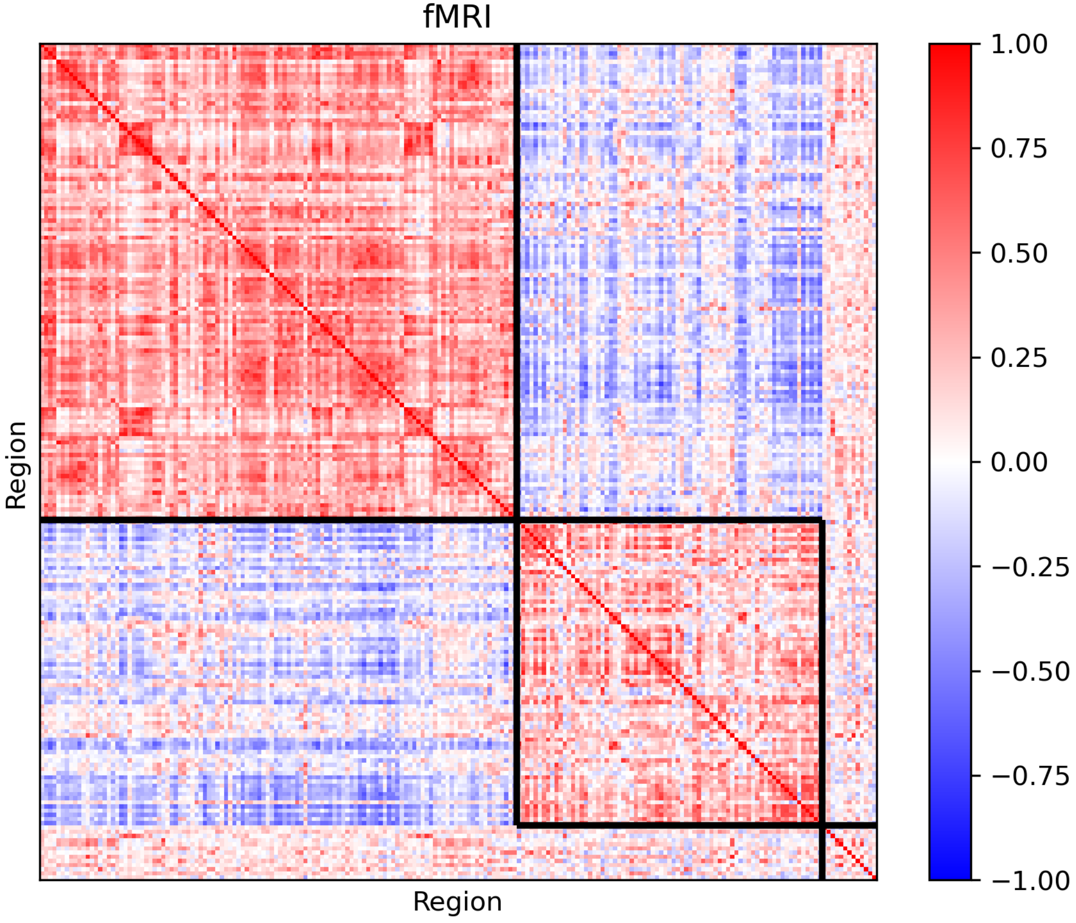
fMRI
Region
Region
1.00
0.75
0.50
0.25
0.00
−0.25
−0.50
−0.75
−1.00

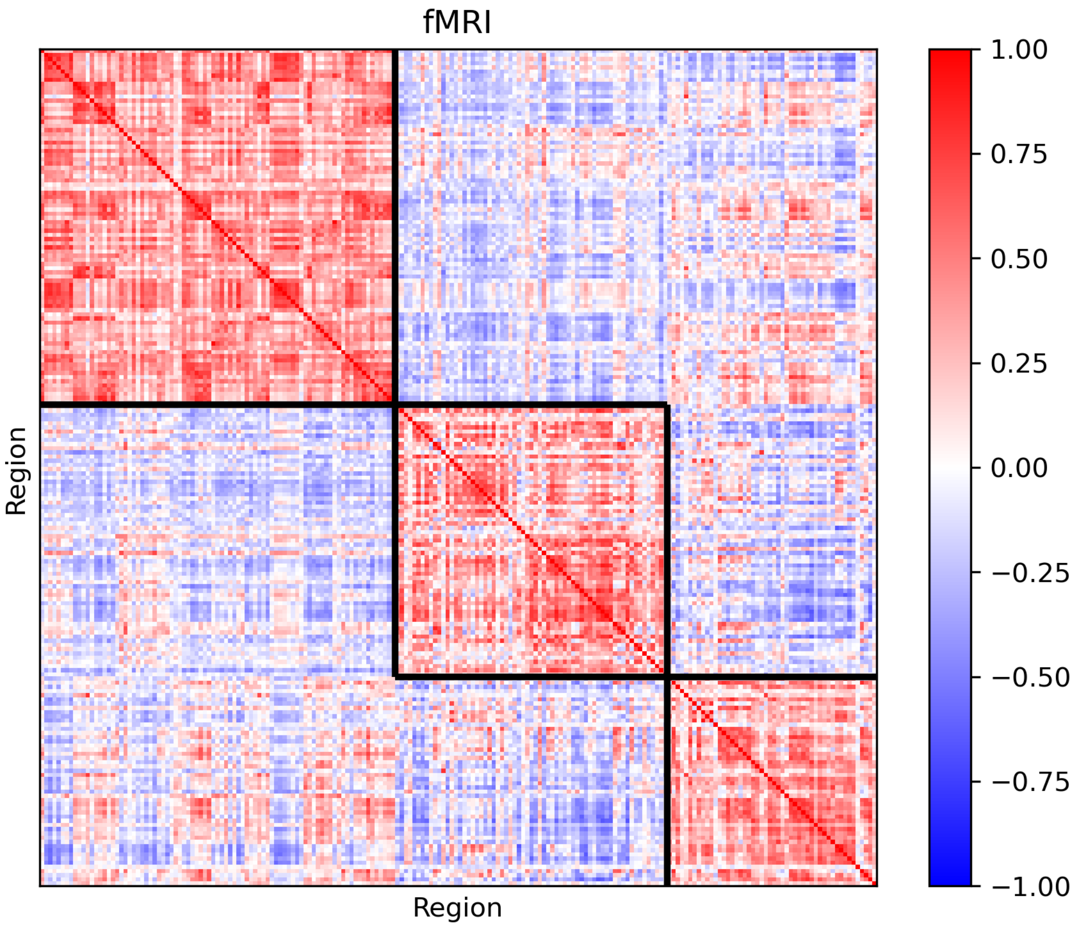
fMRI
Region
Region
1.00
0.75
0.50
0.25
0.00
−0.25
−0.50
−0.75
−1.00

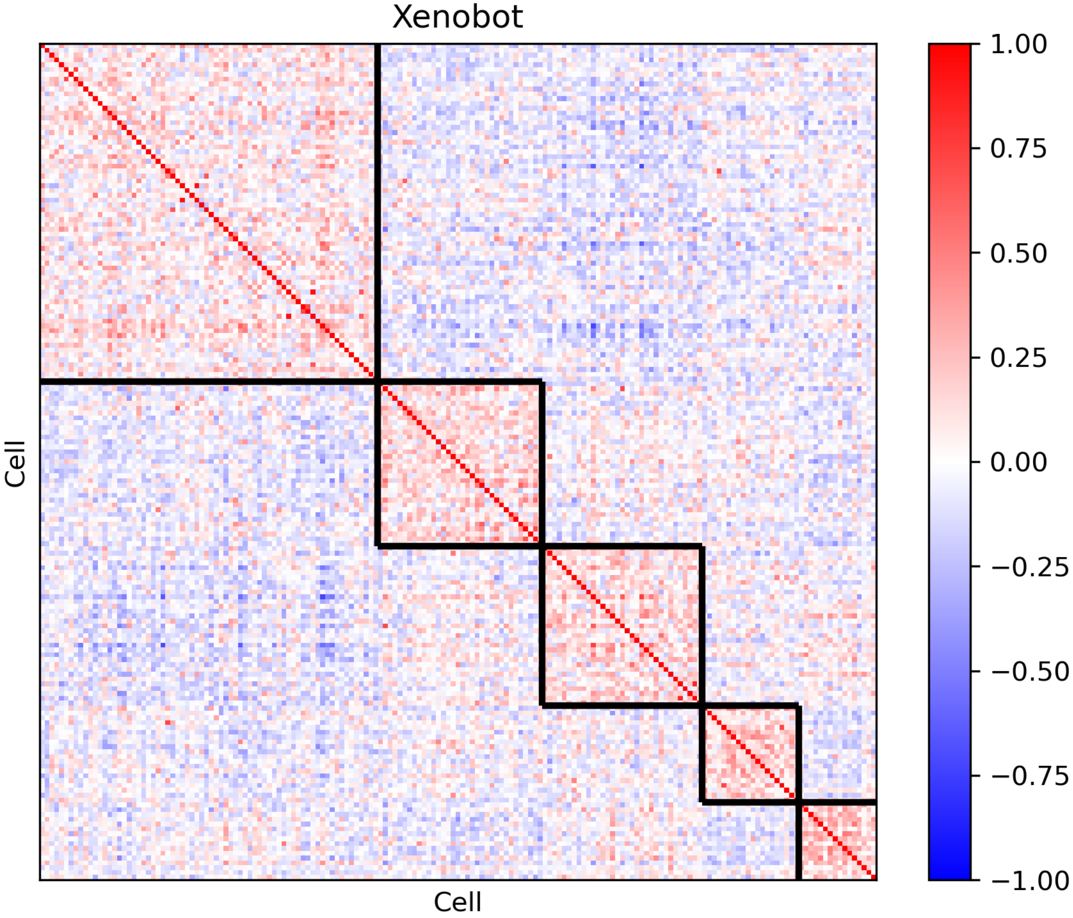
Xenobot
Cell
Cell
1.00
0.75
0.50
0.25
0.00
−0.25
−0.50
−0.75
−1.00

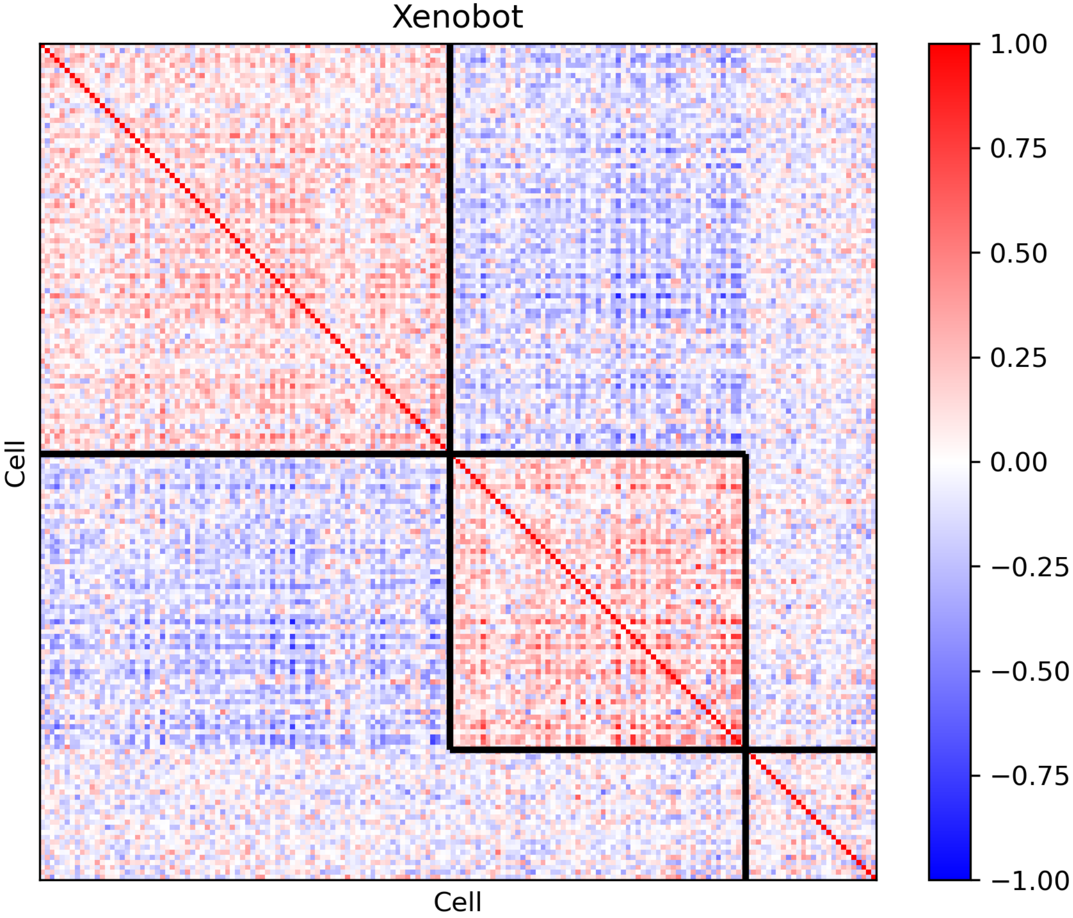
Xenobot
Cell
Cell
1.00
0.75
0.50
0.25
0.00
−0.25
−0.50
−0.75
−1.00

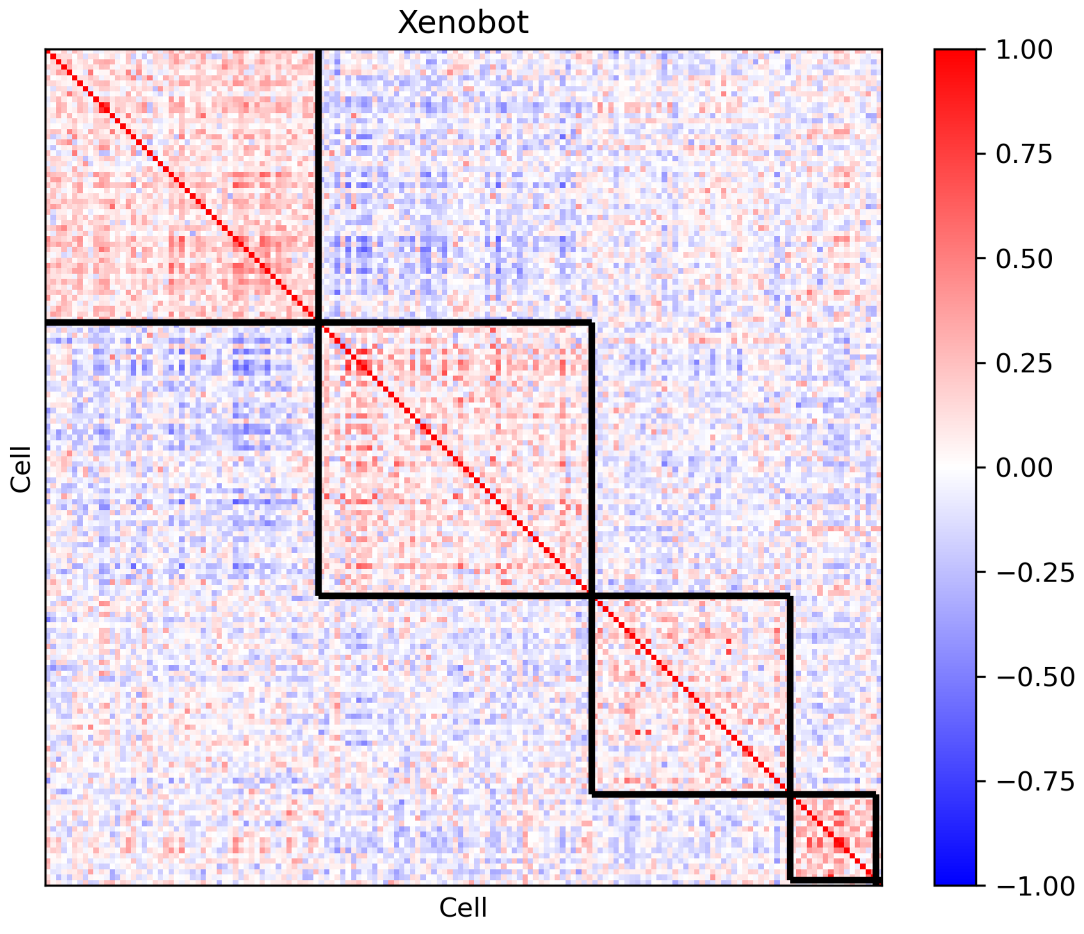

Xenobot
Cell
Cell
1.00
0.75
0.50
0.25
0.00
−0.25
−0.50
−0.75
−1.00

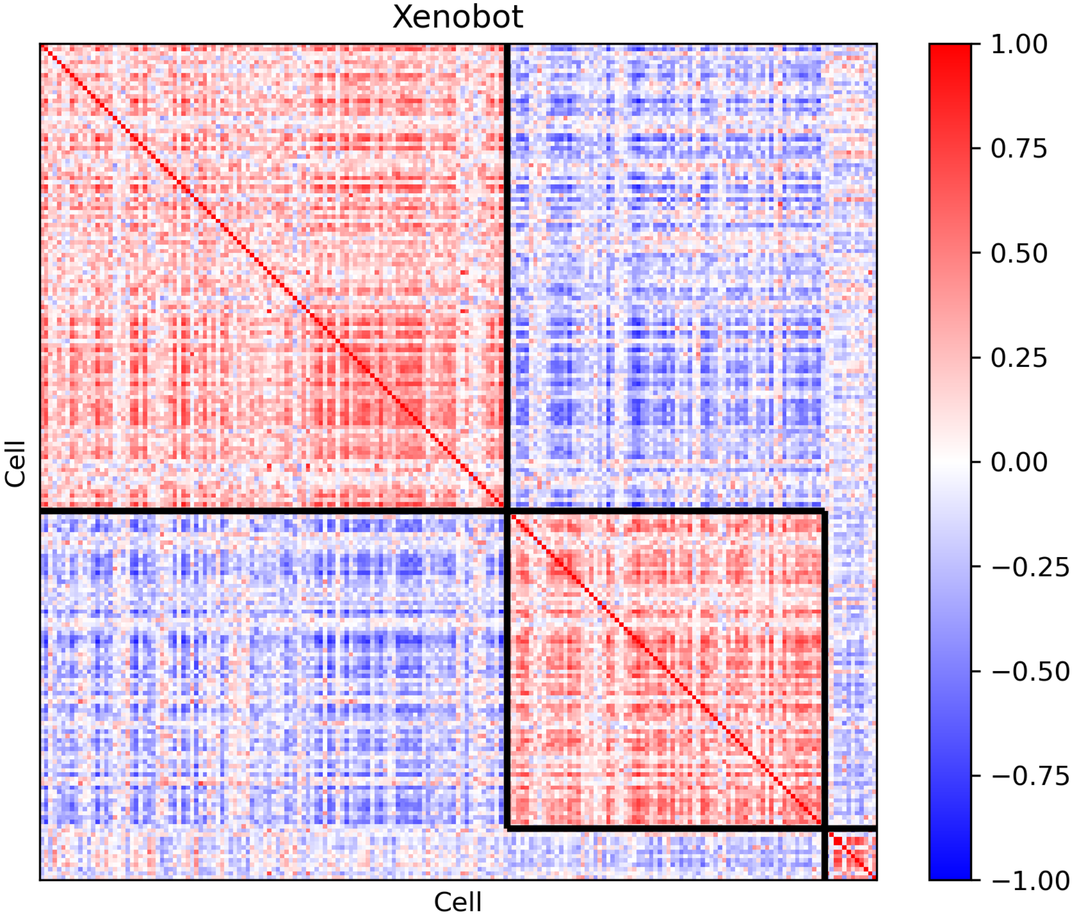

Xenobot
Cell
Cell
1.00
0.75
0.50
0.25
0.00
−0.25
−0.50
−0.75
−1.00

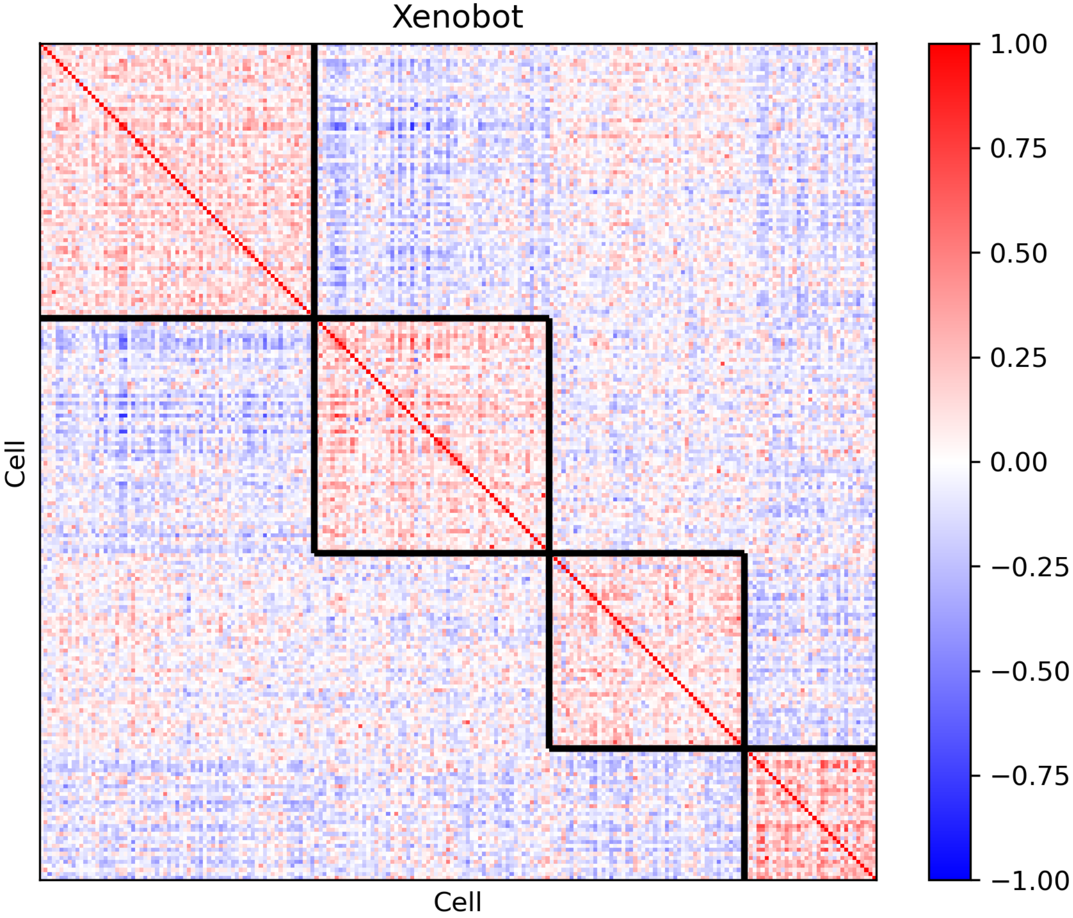
Xenobot
Cell
Cell
1.00
0.75
0.50
0.25
0.00
−0.25
−0.50
−0.75
−1.00

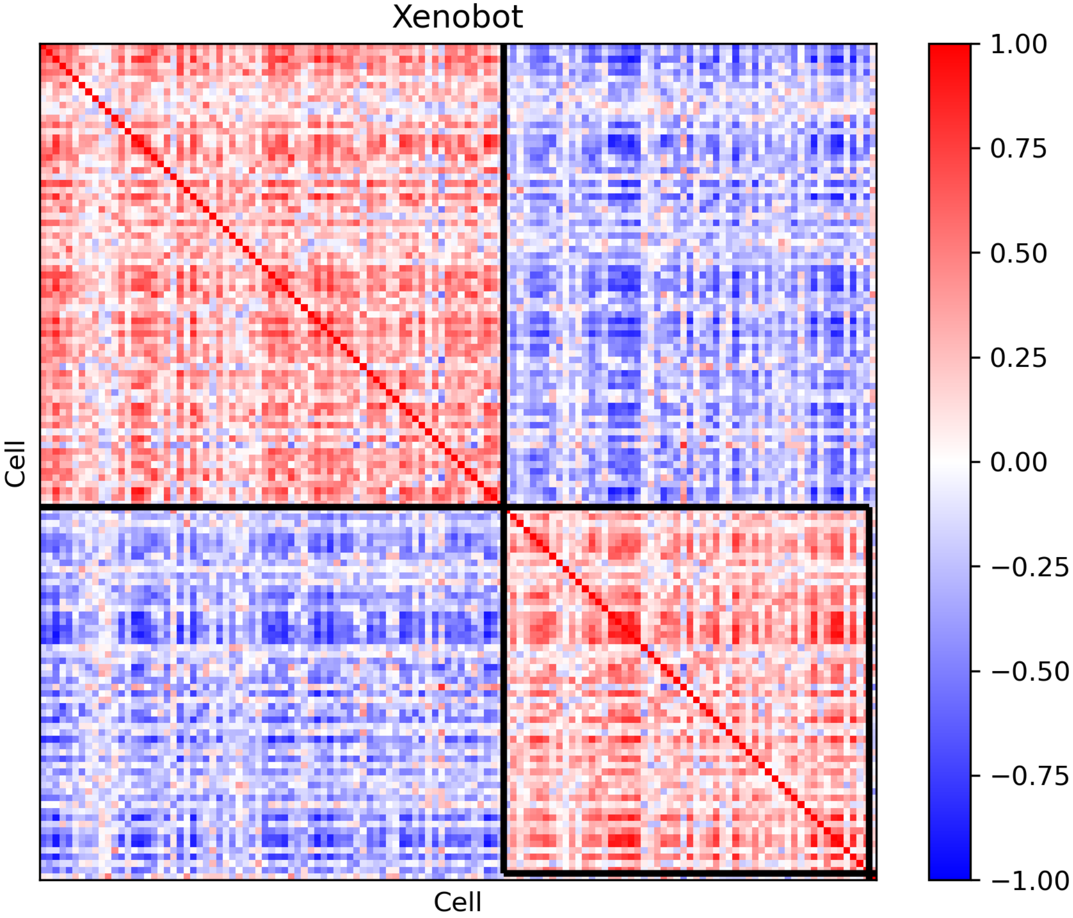
Xenobot
Cell
Cell
1.00
0.75
0.50
0.25
0.00
−0.25
−0.50
−0.75
−1.00

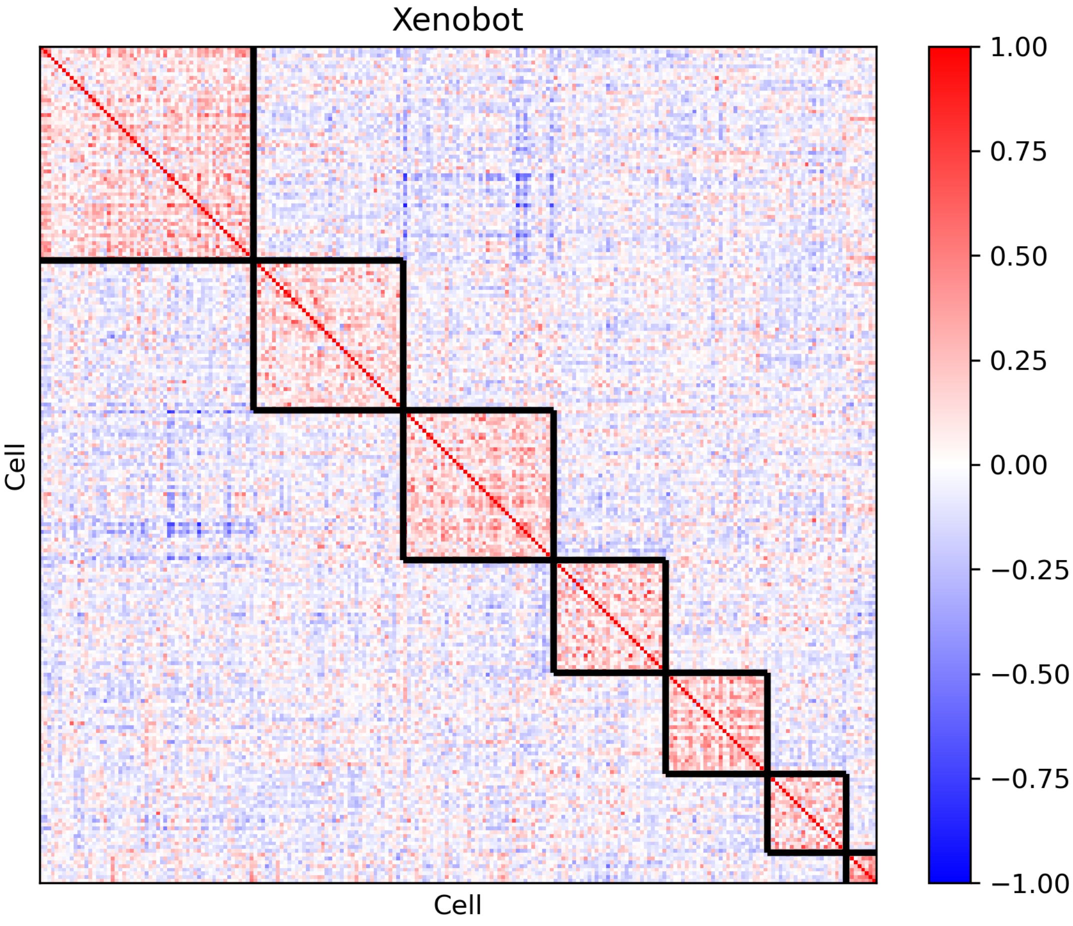
Xenobot
Cell
Cell
1.00
0.75
0.50
0.25
0.00
−0.25
−0.50
−0.75
−1.00

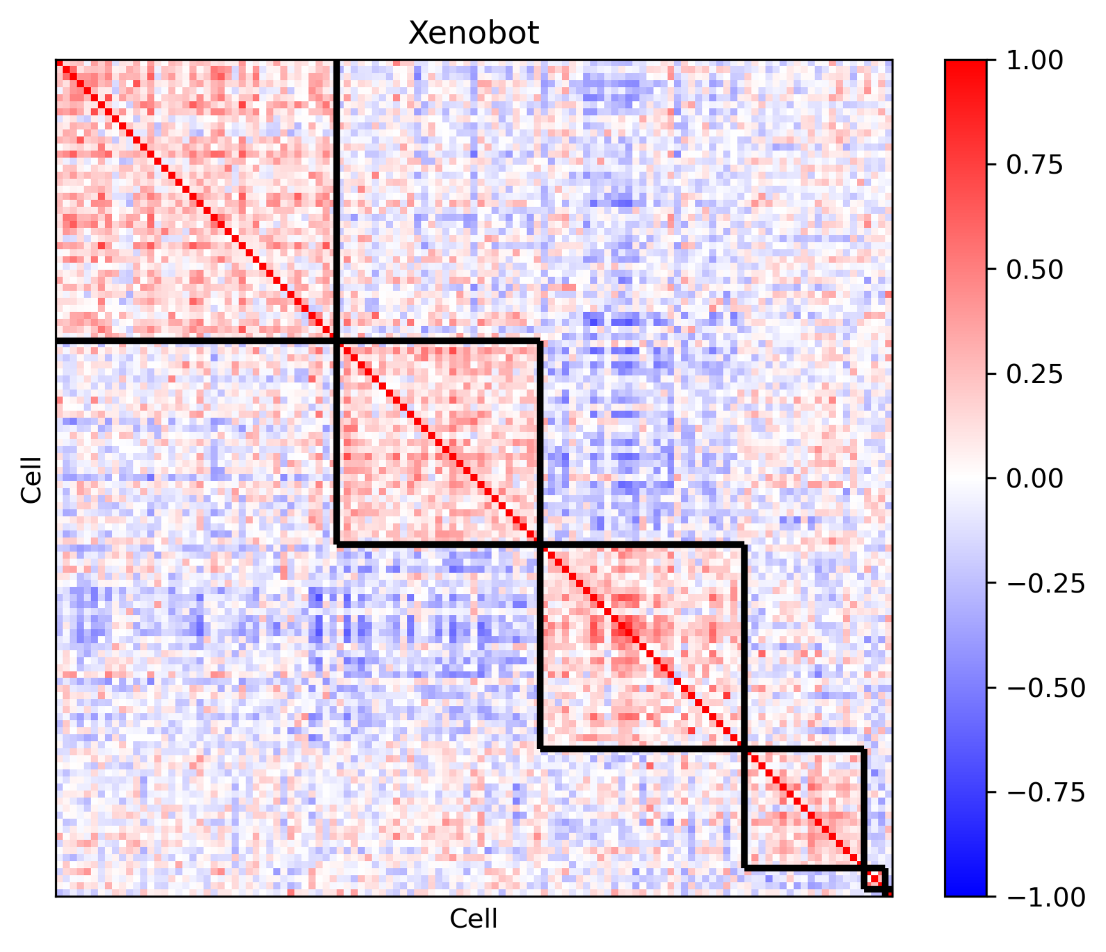
Xenobot
Cell
Cell
1.00
0.75
0.50
0.25
0.00
−0.25
−0.50
−0.75
−1.00

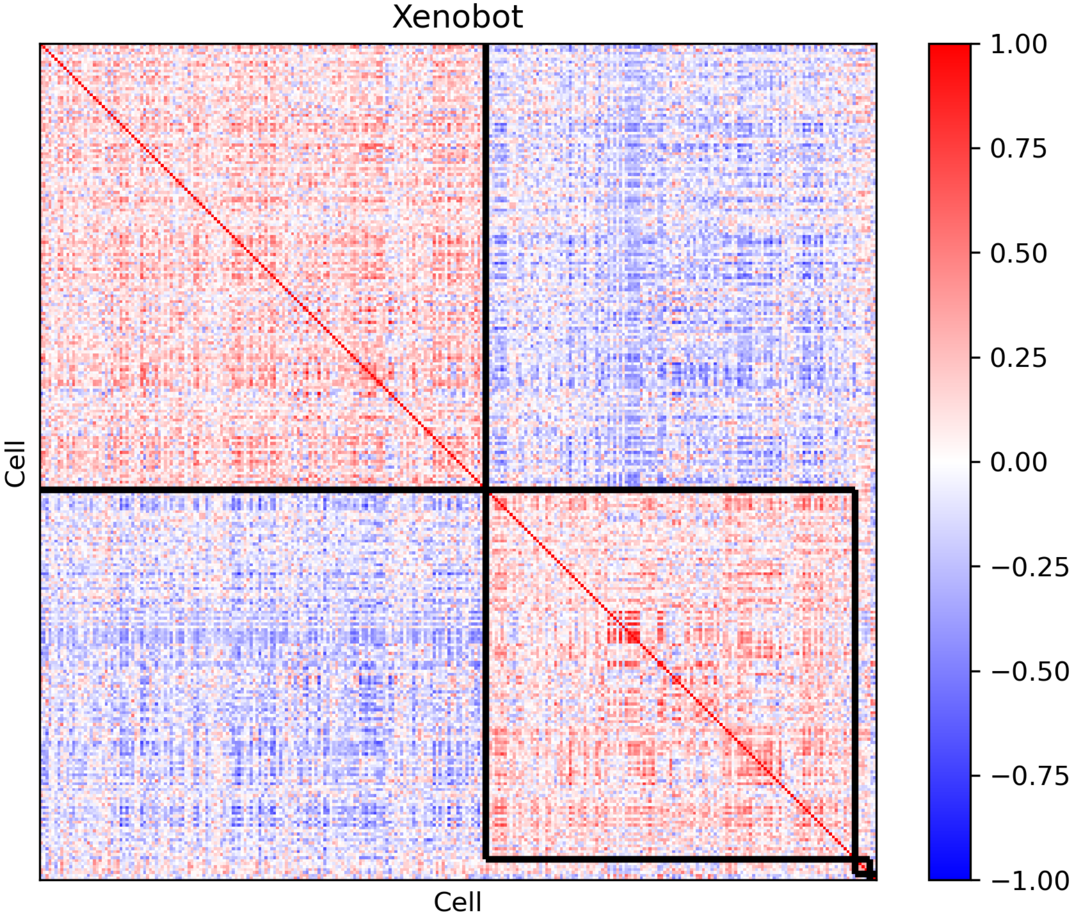

Xenobot
Cell
Cell
1.00
0.75
0.50
0.25
0.00
−0.25
−0.50
−0.75
−1.00

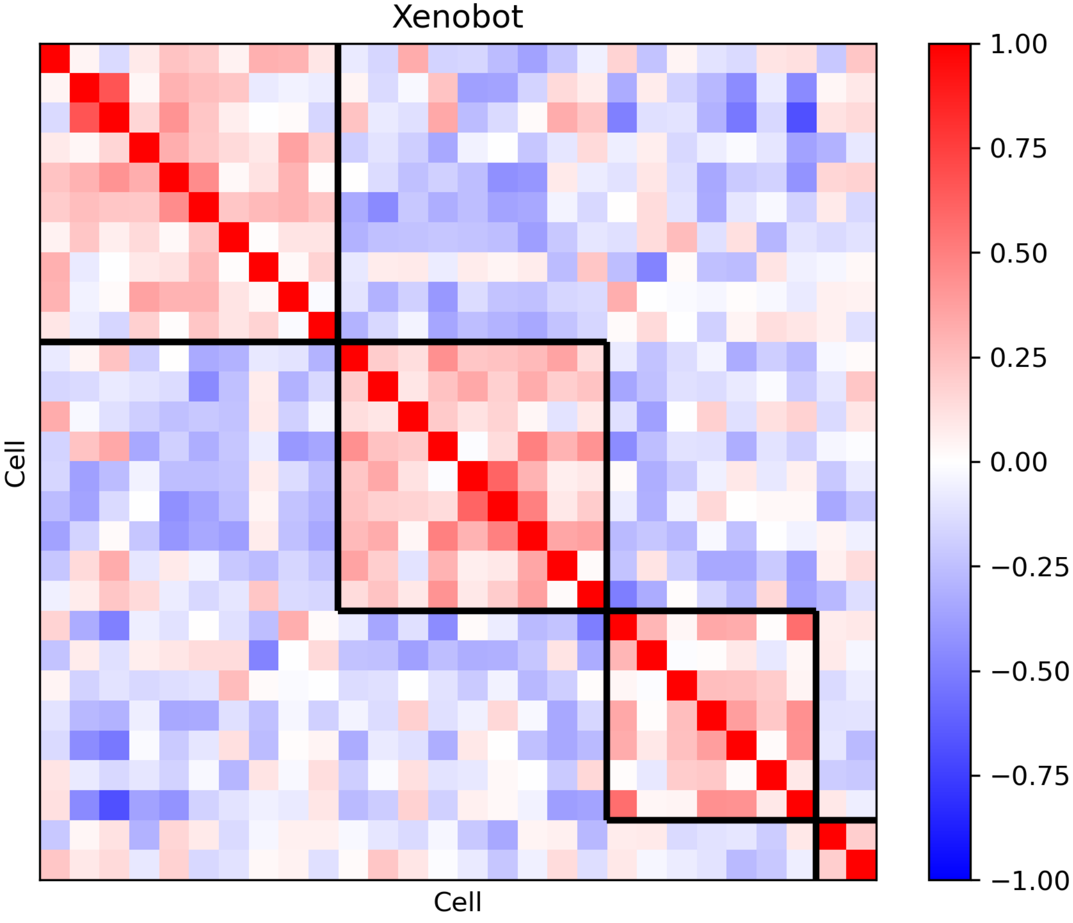
Xenobot
Cell
Cell
1.00
0.75
0.50
0.25
0.00
−0.25
−0.50
−0.75
−1.00

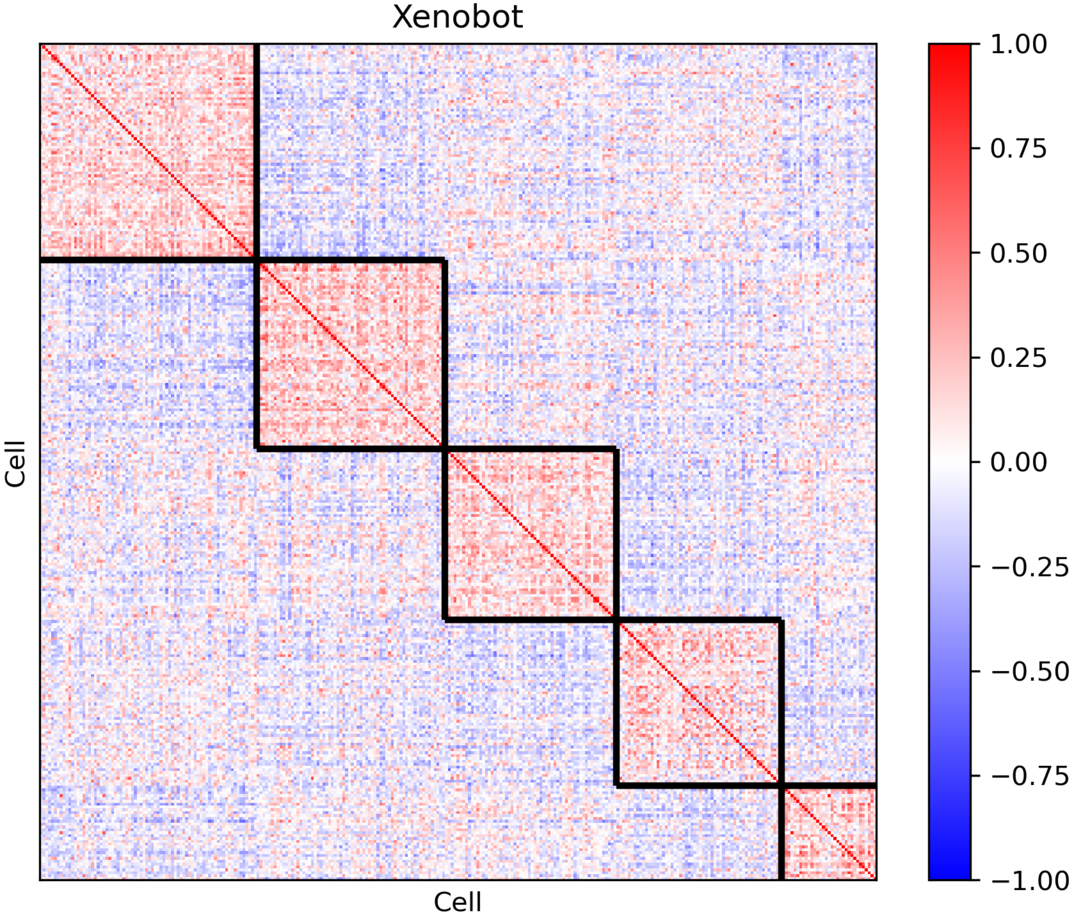
Xenobot
Cell
Cell
1.00
0.75
0.50
0.25
0.00
−0.25
−0.50
−0.75
−1.00

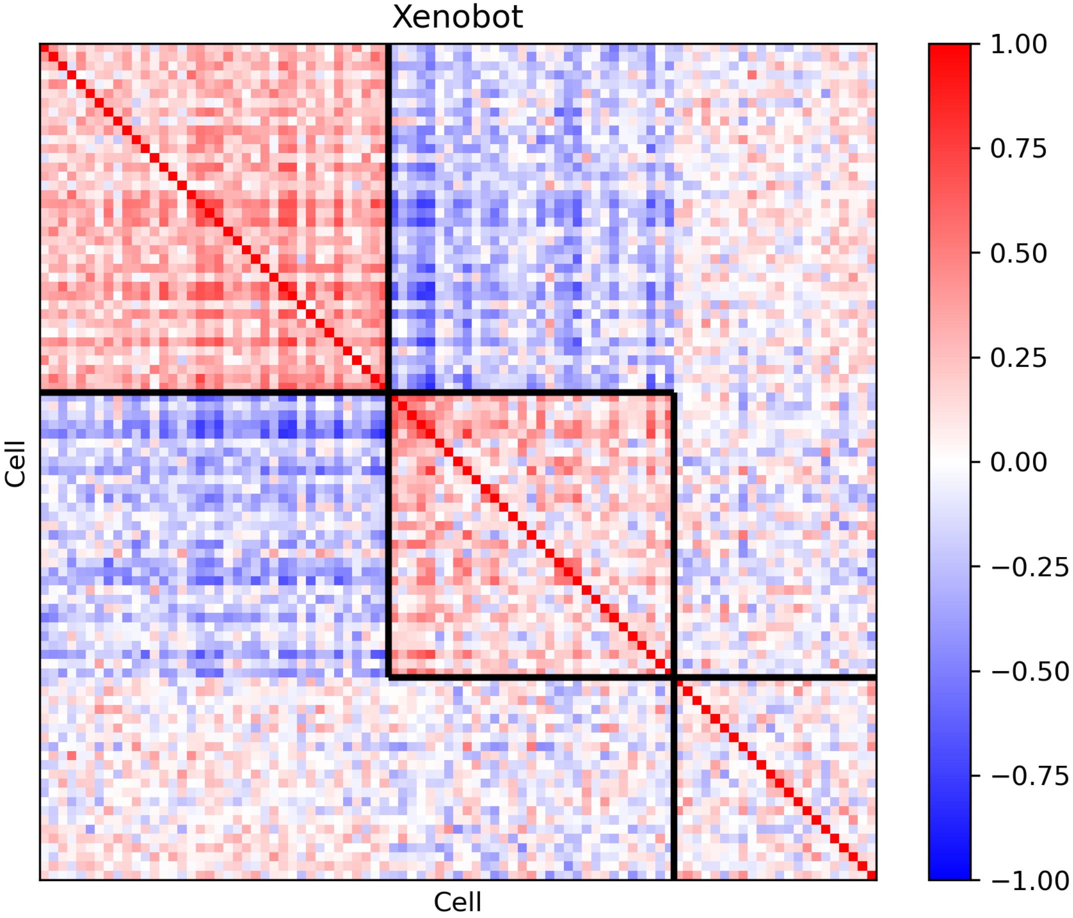
Xenobot
Cell
Cell
1.00
0.75
0.50
0.25
0.00
−0.25
−0.50
−0.75
−1.00

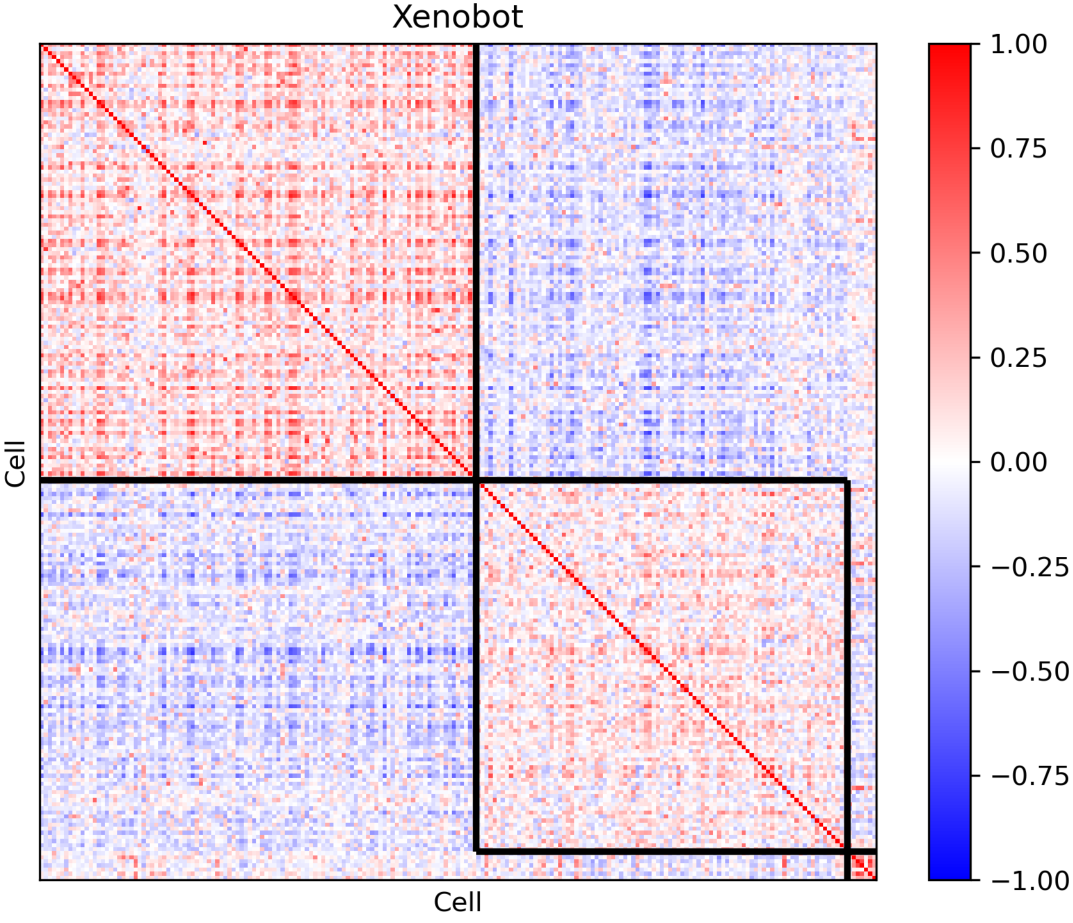
Xenobot
Cell
Cell
1.00
0.75
0.50
0.25
0.00
−0.25
−0.50
−0.75
−1.00

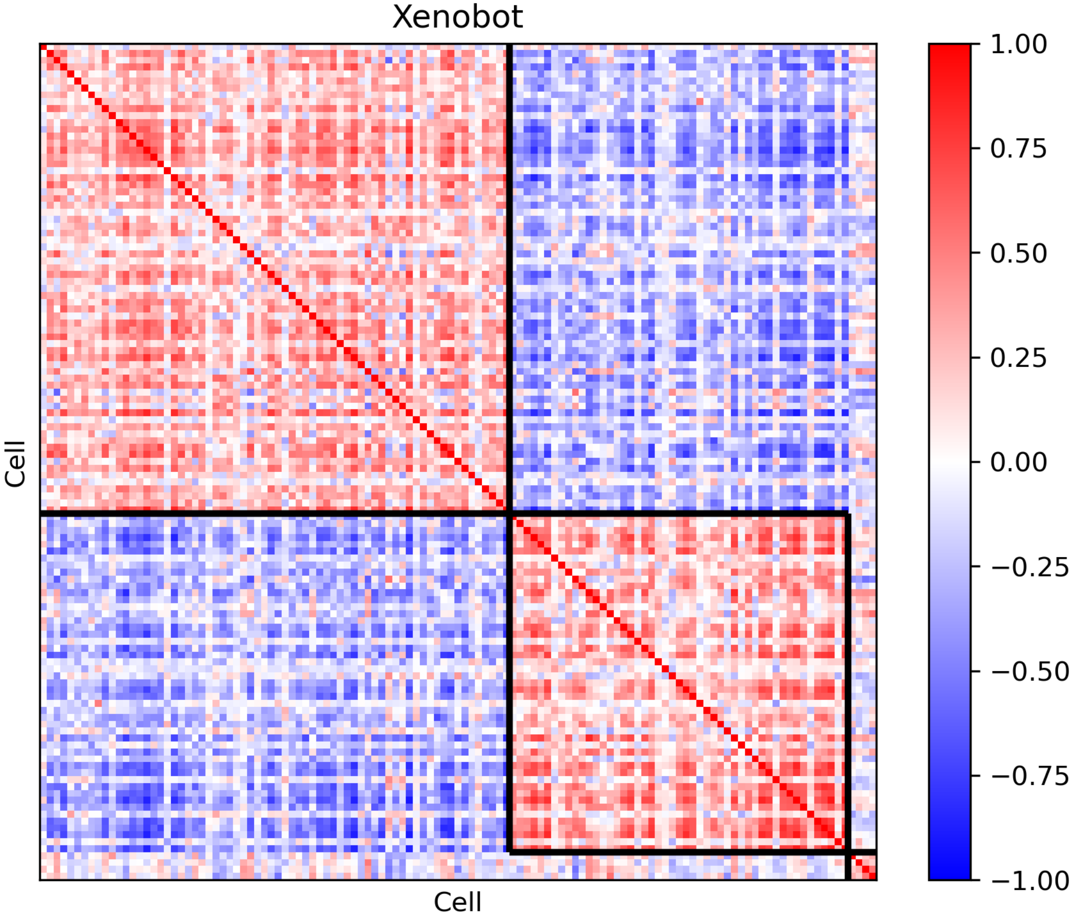
Xenobot
Cell
Cell
1.00
0.75
0.50
0.25
0.00
−0.25
−0.50
−0.75
−1.00

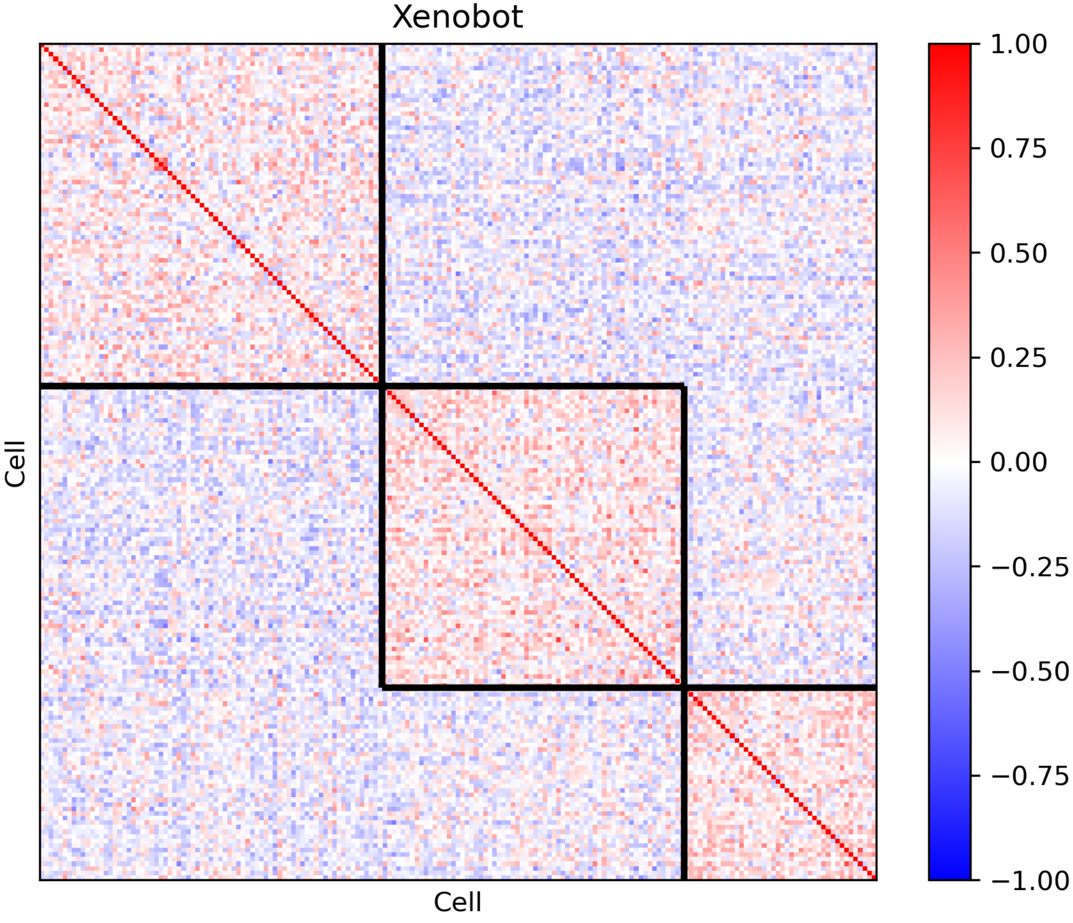
Xenobot
Cell
Cell
1.00
0.75
0.50
0.25
0.00
−0.25
−0.50
−0.75
−1.00

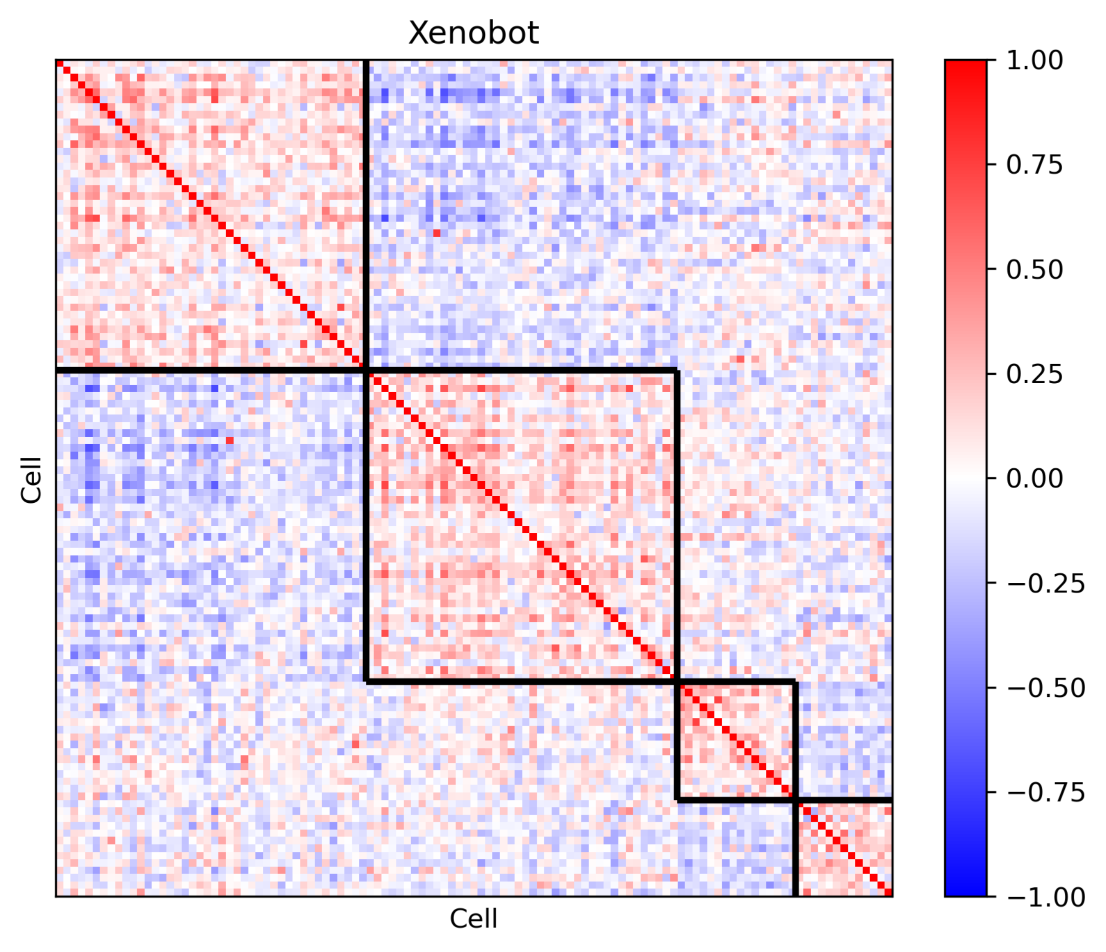
Xenobot
Cell
Cell
1.00
0.75
0.50
0.25
0.00
−0.25
−0.50
−0.75
−1.00

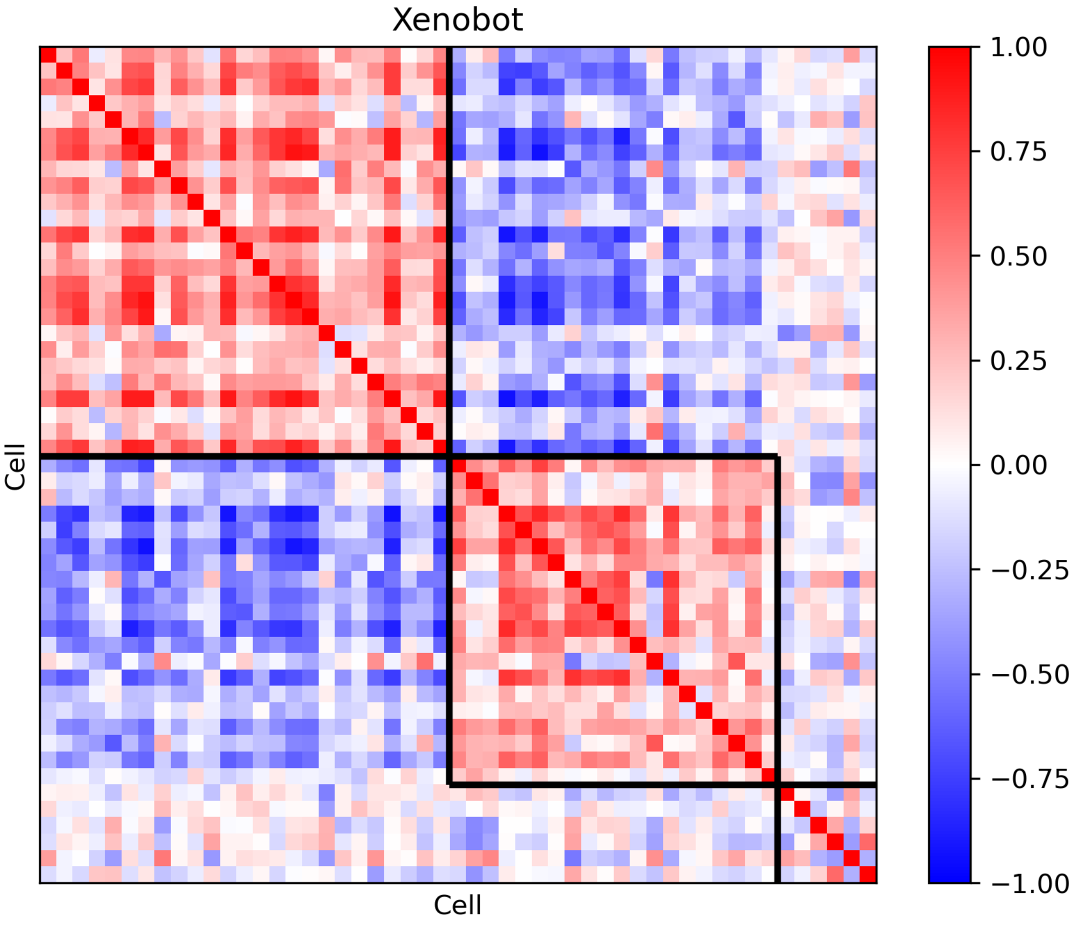
Xenobot
Cell
Cell
1.00
0.75
0.50
0.25
0.00
−0.25
−0.50
−0.75
−1.00

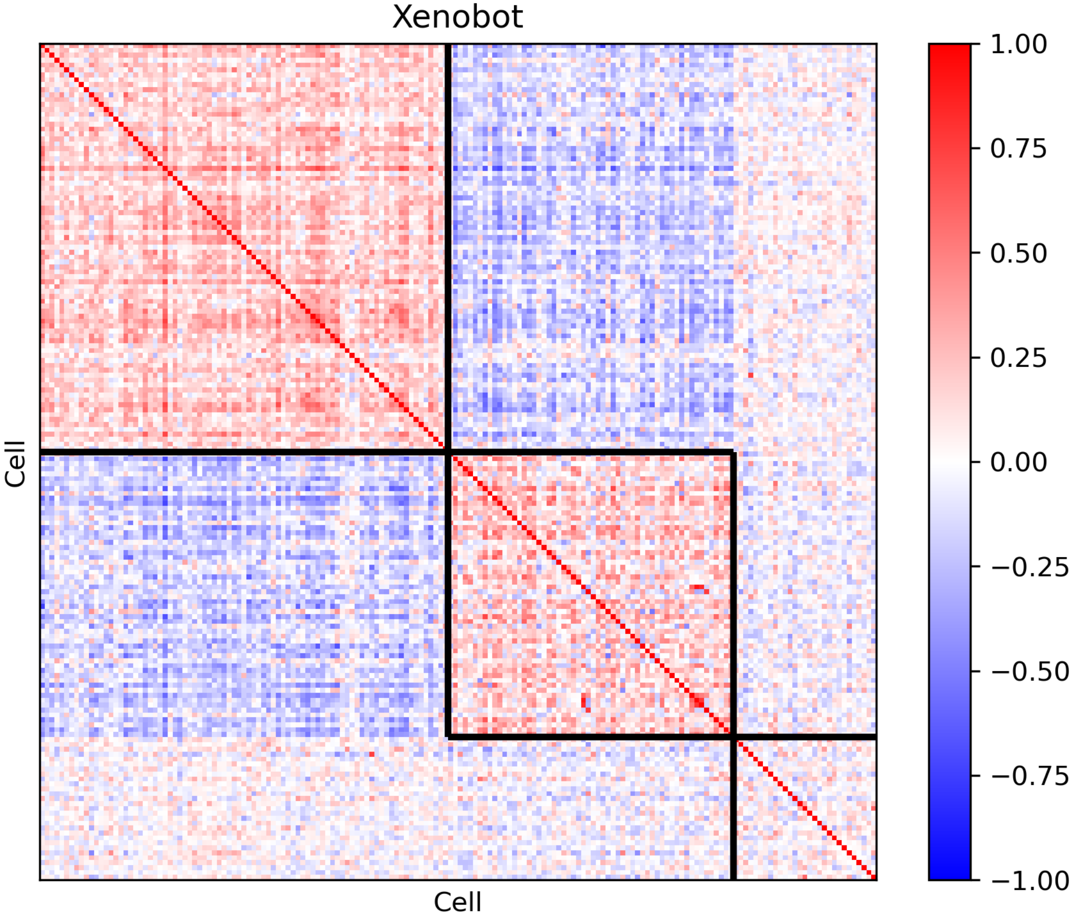
Xenobot
Cell
Cell
1.00
0.75
0.50
0.25
0.00
−0.25
−0.50
−0.75
−1.00

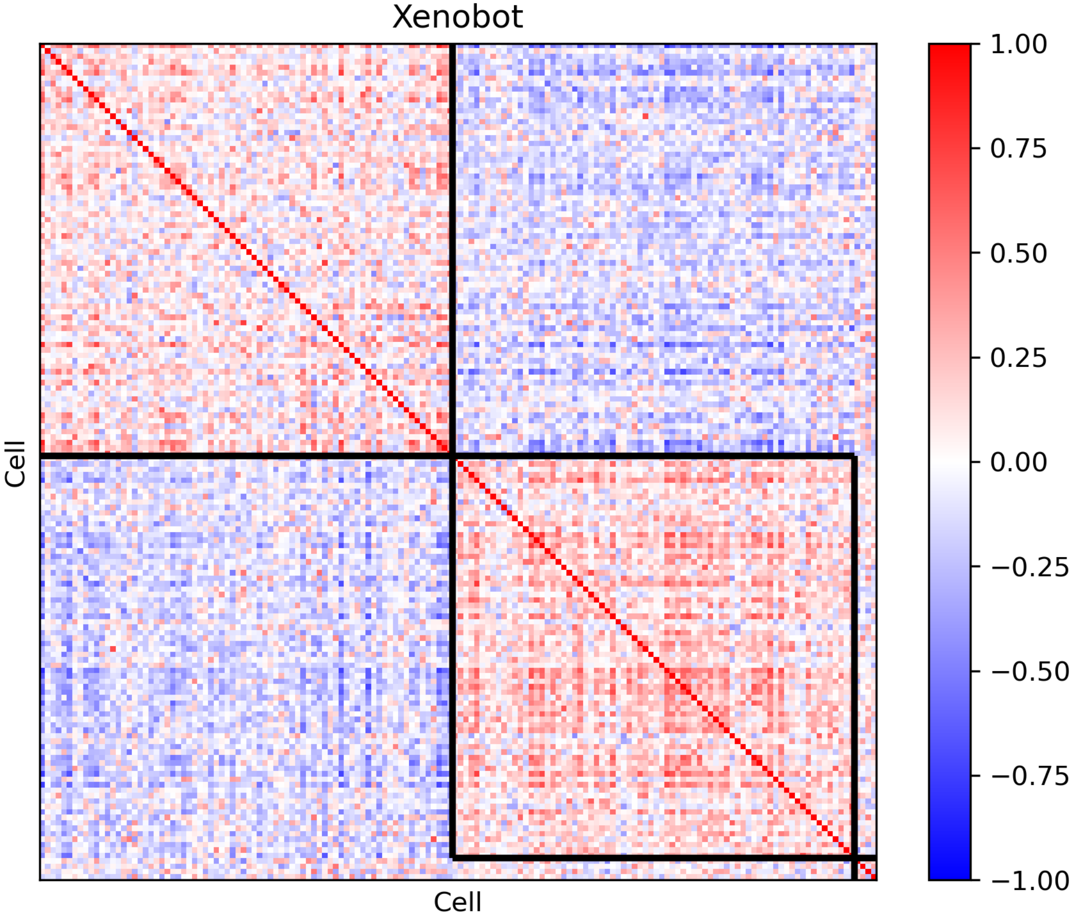
Xenobot
Cell
Cell
1.00
0.75
0.50
0.25
0.00
−0.25
−0.50
−0.75
−1.00

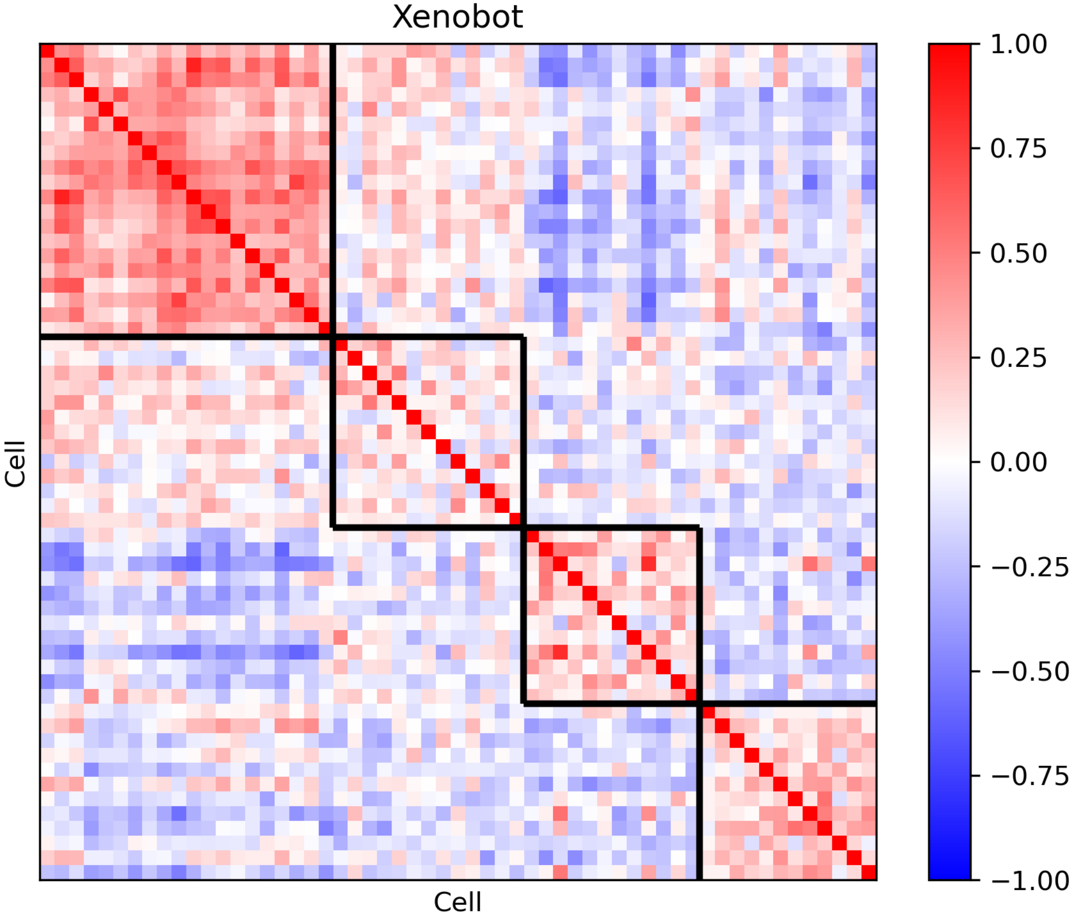
Xenobot
Cell
Cell
1.00
0.75
0.50
0.25
0.00
−0.25
−0.50
−0.75
−1.00

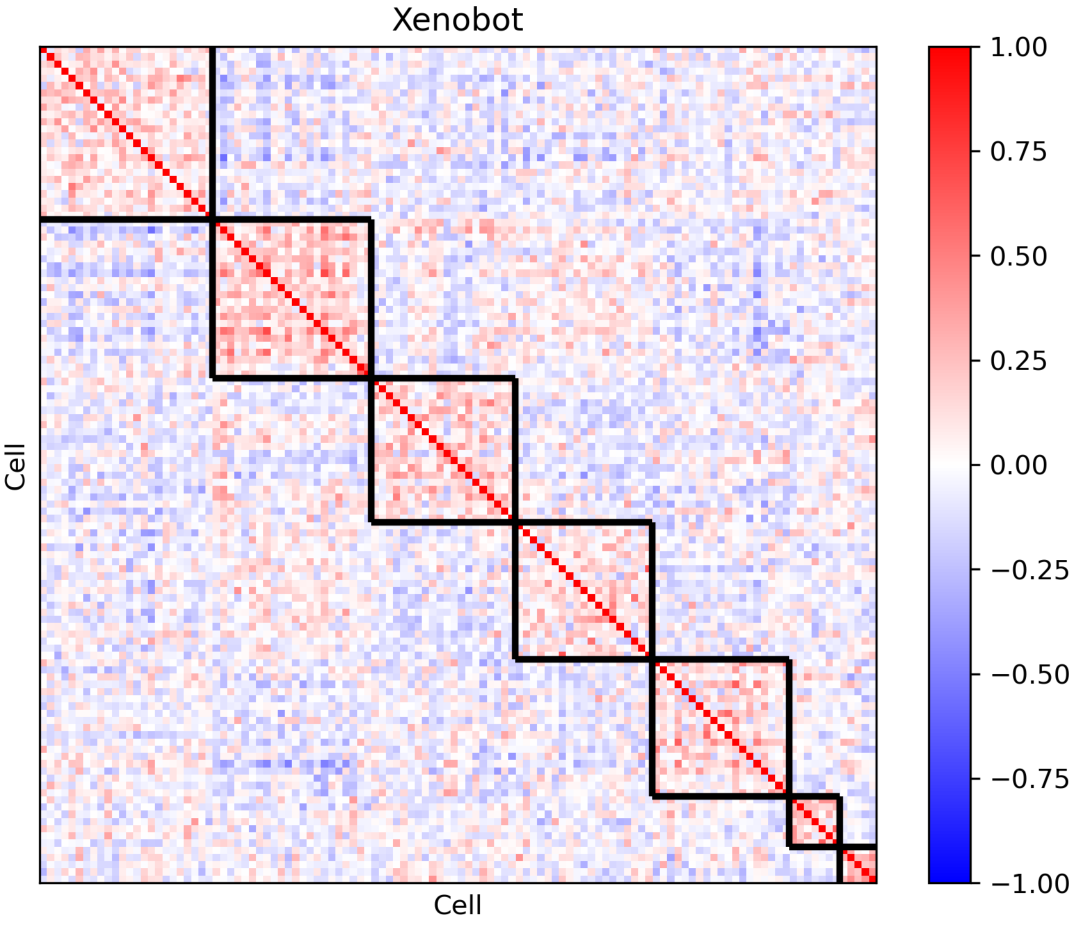

Xenobot
Cell
Cell
1.00
0.75
0.50
0.25
0.00
−0.25
−0.50
−0.75
−1.00

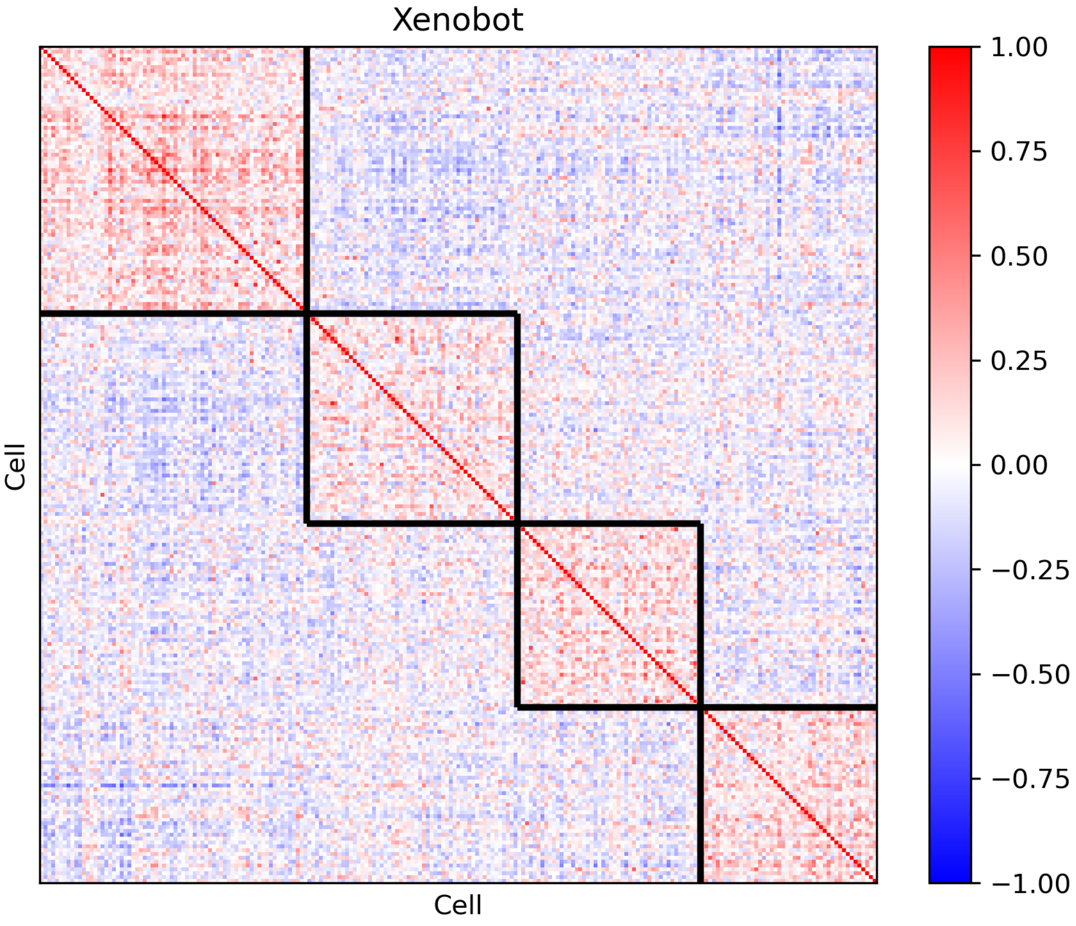
Xenobot
Cell
Cell
1.00
0.75
0.50
0.25
0.00
−0.25
−0.50
−0.75
−1.00

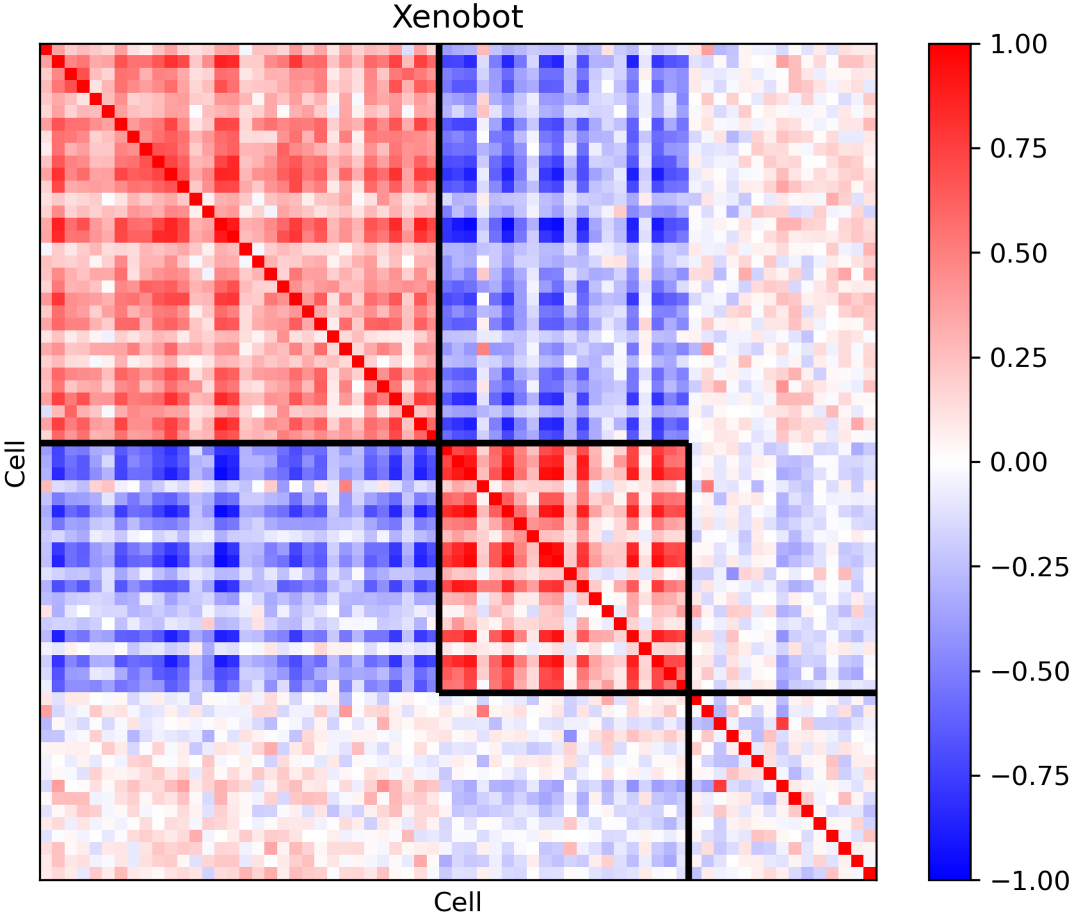
Xenobot
Cell
Cell
1.00
0.75
0.50
0.25
0.00
−0.25
−0.50
−0.75
−1.00

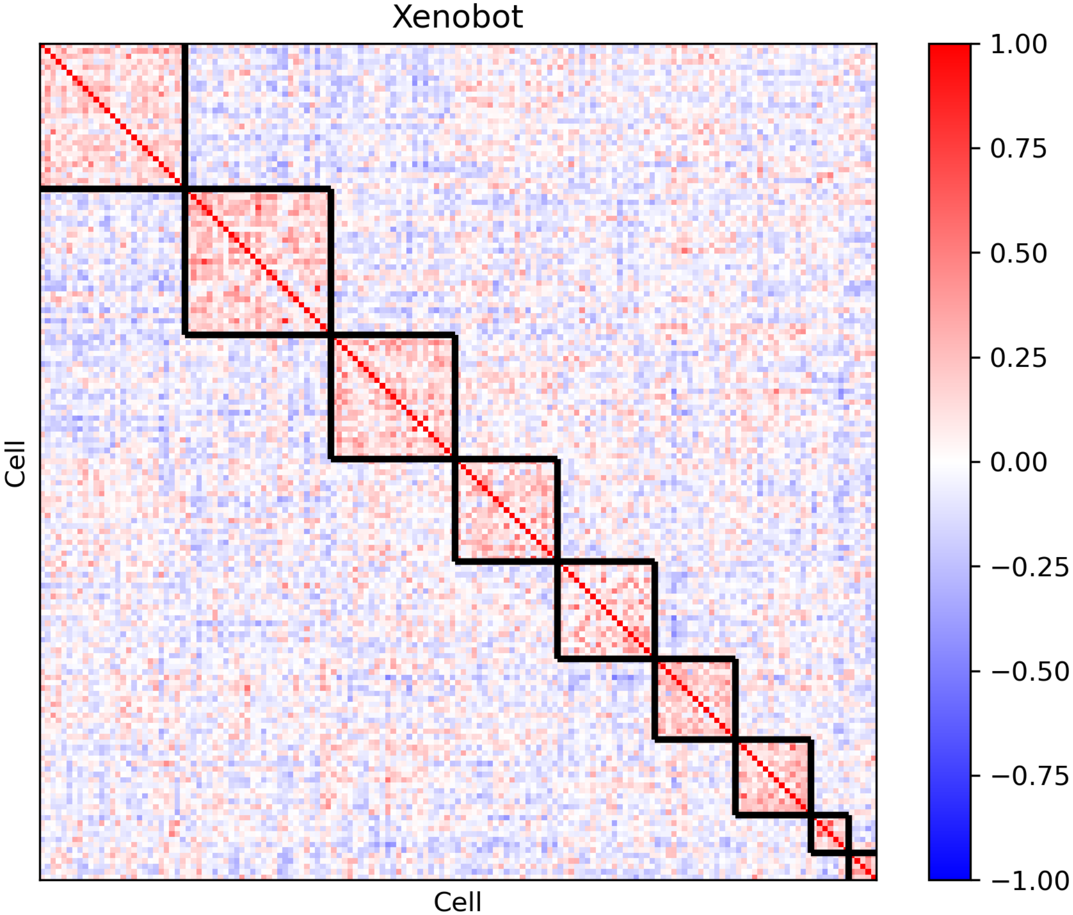
Xenobot
Cell
Cell
1.00
0.75
0.50
0.25
0.00
−0.25
−0.50
−0.75
−1.00

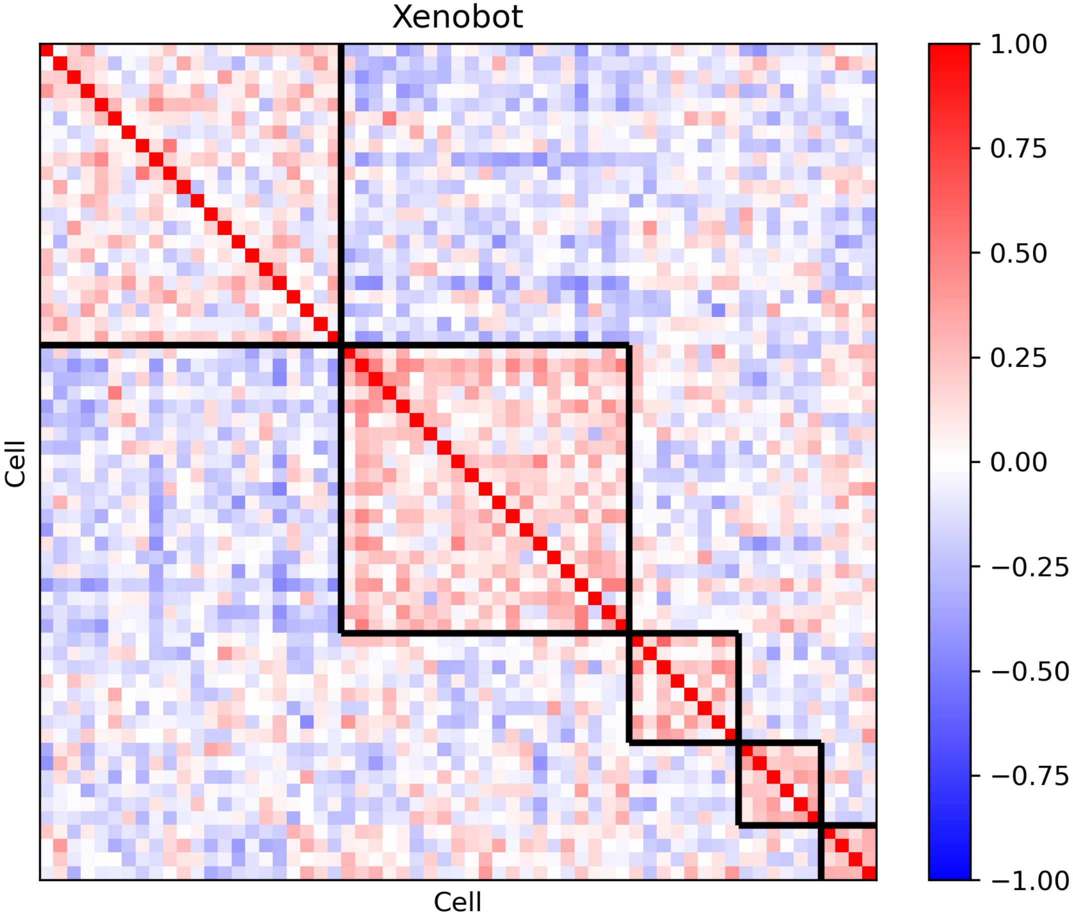

Xenobot
Cell
Cell
1.00
0.75
0.50
0.25
0.00
−0.25
−0.50
−0.75
−1.00

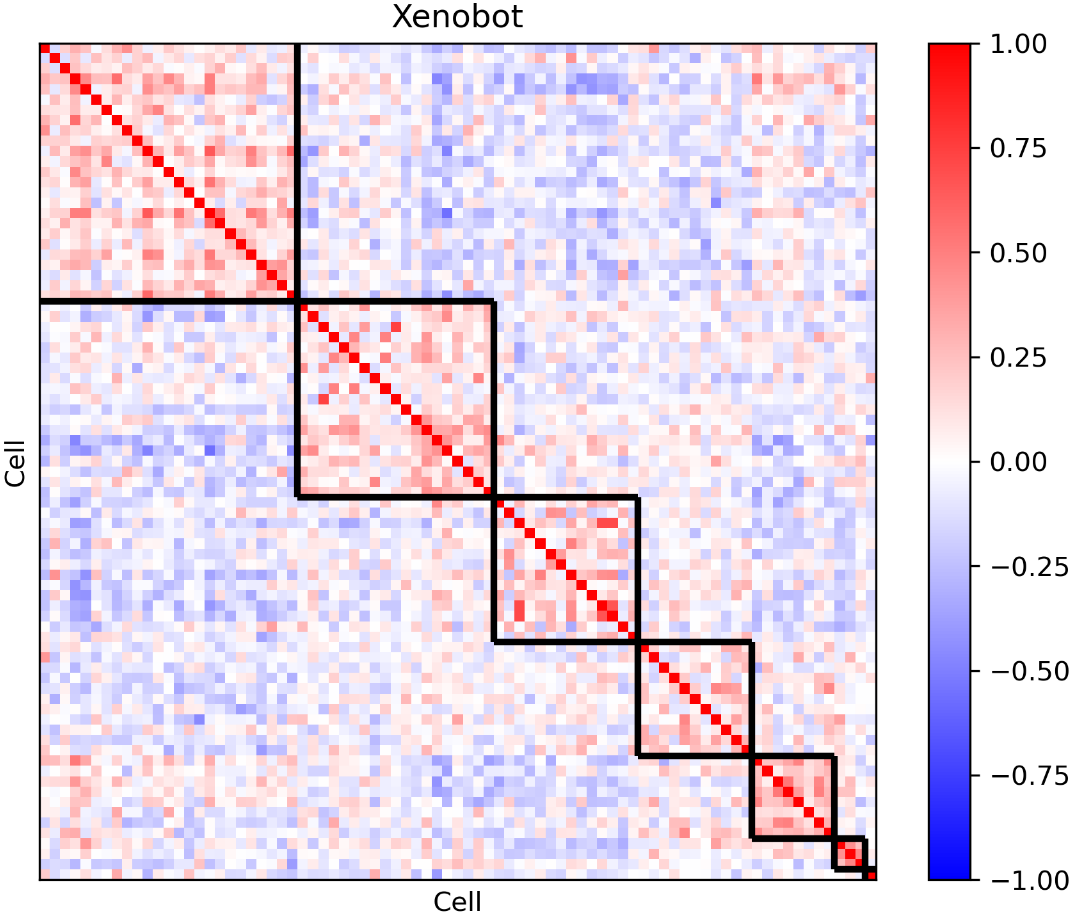
Xenobot
Cell
Cell
1.00
0.75
0.50
0.25
0.00
−0.25
−0.50
−0.75
−1.00

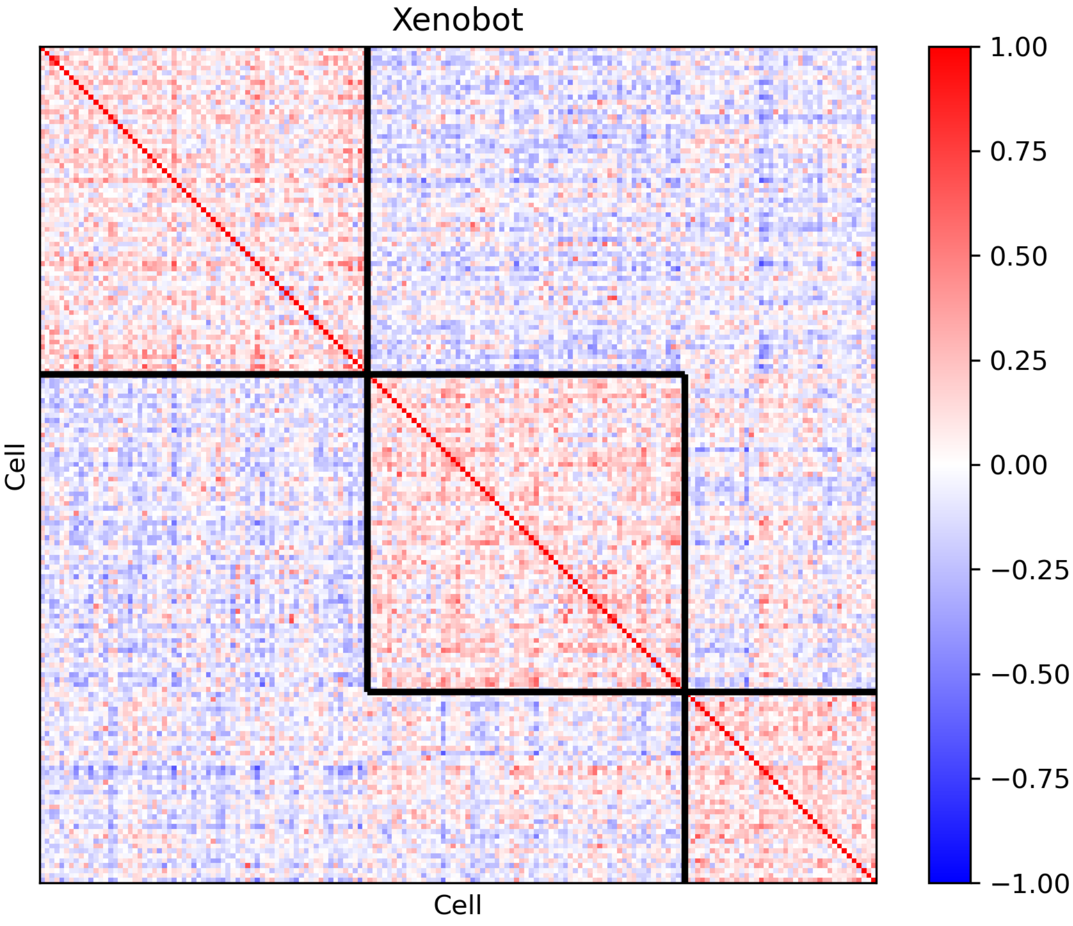
Xenobot
Cell
Cell
1.00
0.75
0.50
0.25
0.00
−0.25
−0.50
−0.75
−1.00

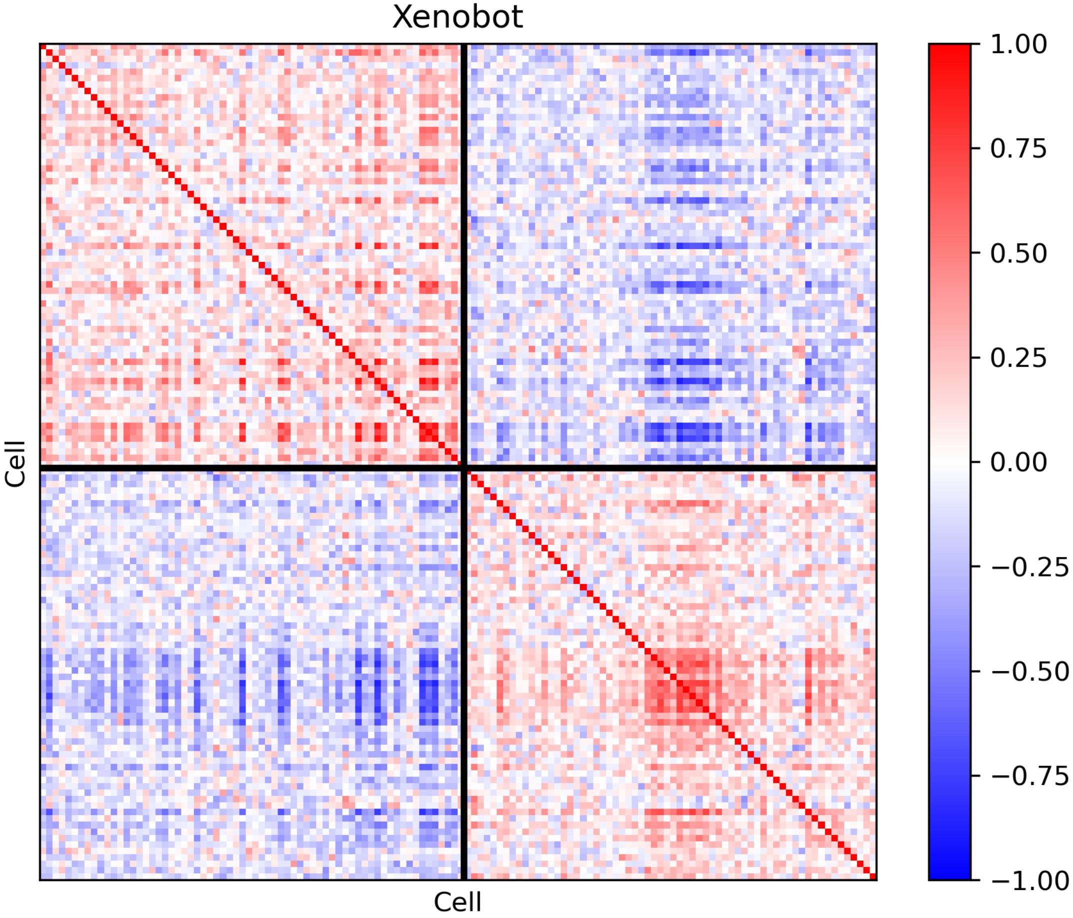
Xenobot
Cell
Cell
1.00
0.75
0.50
0.25
0.00
−0.25
−0.50
−0.75
−1.00
